# A plant-specific syntaxin-6 protein contributes to the intracytoplasmic route for begomoviruses

**DOI:** 10.1101/2020.01.10.901496

**Authors:** Bianca Castro Gouveia-Mageste, Laura Gonçalves Costa Martins, Maximiller Dal-Bianco, João Paulo Batista Machado, José Cleydson Ferreira da Silva, Alice Y Kim, Junshi Yazaki, Anésia Aparecida dos Santos, Joseph R Ecker, Elizabeth Pacheco Batista Fontes

**Affiliations:** National Institute of Science and Technology in Plant-Pest Interactions, Bioagro, Universidade Federal de Viçosa, Viçosa, Minas Gerais, Brazil; Department of Biochemistry and Molecular Biology, Universidade Federal de Viçosa, Viçosa, Minas Gerais 36570-000, Brazil; Agronomy Institute, Universidade Federal de Viçosa, Campus Florestal, Florestal, Minas Gerais, Brazil; Genomic Analysis Laboratory, Plant Biology Laboratory, Salk Institute for Biological Studies, La Jolla, CA 92037, USA; RIKEN Center for Integrative Medical Sciences, Yokohama City, Kanagawa 230-0045, Japan; Departament of General Biology, Universidade Federal de Viçosa, Viçosa, Minas Gerais, Brazil; Howard Hughes Medical Institute and Plant Biology Laboratory, The Salk Institute of Biological Studies, La Jolla, United States of America

## Abstract

Due to limited free diffusion in the cytoplasm, viruses must use active transport mechanisms to move intracellularly. Nevertheless, how the plant ssDNA begomoviruses hijacks the host intracytoplasmic transport machinery to move from the nucleus to the plasmodesmata remains enigmatic. Here, we identified nuclear shuttle protein (NSP)-interacting proteins from Arabidopsis by probing a protein microarray and demonstrated that the *Cabbage leaf curl virus* (CabLCV) NSP, a facilitator of the nucleocytoplasmic trafficking of viral (v)DNA, interacts with an endosomal vesicle-localized plant-specific syntaxin-6 protein, designated NSP-interacting syntaxin-6 domain-containing protein (NISP) *in planta*. NISP displays a pro-viral function, but not the syntaxin-6 paralog AT2G18860 that failed to interact with NSP. Consistent with these findings, *nisp-1* mutant plants were less susceptible to begomovirus infection, a phenotype reversed by NISP complementation. NISP-overexpressing lines accumulated higher levels of viral DNA than wild-type. Furthermore, NISP interacted with NIG, an NSP-interacting GTPase involved in NSP-vDNA nucleocytoplasmic translocation. The NISP-NIG interaction was enhanced by NSP. We also showed that NISP associates with vDNA and might assemble a NISP-NIG-NSP-vDNA-complex. NISP may function as a docking site for recruiting NIG and NSP into endosomes, providing a mechanism for the intracytoplasmic translocation of the NSP-vDNA complex towards to and from the cell periphery.

**Author Summary:** As viruses must use an active and directed intracellular movement, they hijack the intracellular host transport system for their own benefits. Therefore, the identification of interactions between host proteins and begomovirus movement proteins should target the intracellular transport machinery. This work focused on the identification of these protein-protein interactions; it addressed the molecular bases for the intracellular transport of begomoviruses. We used a protein microarray to identify cellular partners for the movement protein (MP) and the viral nuclear shuttle protein (NSP), which is a facilitator of the nucleocytoplasmic trafficking of viral (v)DNA. We identified relevant protein-protein interaction (PPI) hubs connecting host and viral proteins. We revealed a novel NSP-interacting protein, which functions in the intracytoplasmic transport of proteins and DNA from begomoviruses and was designated NSP-interacting syntaxin domain-containing protein (NISP). Our data suggest an intracellular route connecting the release of newly synthesized begomoviral DNA in the cytosol with the cell surface. Resolving viral DNA-host protein complexes led to the identification of a novel class of components of the cell machinery and a representative member, NISP, that functions as a susceptibility gene against begomoviruses. As geminiviruses pose a severe threat to agriculture and food security, this recessive gene can now be exploited as a target for engineering resistance by gene editing in crops.

## Introduction

As obligate intracellular parasites, viruses must enter the host cells and hijack the cell machinery to complete their life cycle. Independent on the molecular mechanisms for virus entry into cells, which differ entirely between animal and plant viruses, once in the cytoplasm, both plant and animal viral genomes move intracellularly to specific cellular sub-compartments for replication, transcription or encapsidation [1,2]. Due to the limitation of free diffusion in the cytoplasm, viruses have invoked the endogenous host movement mechanisms for an active and directed intracytoplasmic translocation. In fact, almost all viruses are capable of subverting the microtubule transport system to some extent to replicate and spread. Plant viruses must also use an active mechanism to move the viral genome from the replication site to the plasmodesmata, which are intercellular cytoplasmic bridges between plant cells [2,3]. To promote this typical cell-to-cell movement, plant viruses have evolved movement proteins that increase the size exclusion limit of the plasmodesmata and actively promote the viral nucleic acid translocation into uninfected, adjacent cells [4]. In the case of viruses that replicate in the nuclei of infected cells, they have to be directed to the nuclear periphery, followed by nuclear entry. Likewise, the newly-synthesized viral proteins and viral genomes must translocate throughout the cytoplasm during exit. In animal cells, the retroviruses and DNA viruses deploy the microtubule motor system and microtubule cytoskeleton to move viral proteins and genomes into the nucleus and back to the plasma membrane [1, 5]. Geminiviruses are typical examples of plant DNA viruses that replicate in the nuclei of infected cells, but the underlying mechanisms for the intracytoplasmic movement of viral proteins and viral DNA (vDNA) are far less understood.

Geminiviruses are circular single-stranded (ss) DNA viruses, which replicate via double-stranded (ds) DNA intermediates in the nucleus of infected cells, infect a broad spectrum of economically important crops and cause significant yield losses worldwide [6]. The *Geminiviridae* family includes nine genera, but approximately 75% of the identified species belong to the genus *Begomovirus*, which are whitefly-transmitted, infect dicotyledonous species, and display a genome configuration that can be either monopartite or bipartite [7,8]. The genomic components of the bipartite begomoviruses are designated DNA-A and DNA-B. The DNA-A encodes proteins required for viral replication (AC1 or Rep, REn or AC3), transactivation of viral genes (AC2 or TRaP), encapsidation of viral genome (AV1 or CP) and suppression of host defenses (AC4 and TrAP). DNA-B encodes the nuclear shuttle protein (NSP or BV1) and the movement protein (MP or BC1) involved in intracellular and cell-to-cell movement of vDNA.

Although the role of MP-NSP complex formation in mediating intracellular and intercellular transport of begomoviruses is widely accepted, the molecular mechanisms underlying the action of these viral proteins at individual stages of the intracellular and intercellular movement are still debated. Some studies indicated that NSP from *Bean dwarf mosaic virus* (BDMV) and *Abutilon mosaic virus* (AbMV) complexes with ss-vDNA and ds-vDNA presumably in the nuclei of infected cells, whereas NSP from *Squash leaf curl virus* (SLCV) binds only ss-vDNA *in vitro* [9, 10, 11]. NSP also interacts with MP in the cytoplasm [12], which promotes the directionality of virus translocation to the cell surface [12, 13]. The mechanism for vDNA exit from the nucleus remains unsolved. It has been conceptually accepted that NSP facilitates the nuclear exit of newly-replicated vDNA via nuclear pores [10, 14-17]. However, the interactions with the host nuclear transport machinery have not been documented. A piece of this puzzle was obtained with the identification of an NSP-interacting GTPase (NIG), which interacts with NSP at the cytosolic side of the nuclear pore complex and facilitates the release of NSP-vDNA complex from the nuclear envelope into the cytosol [12, 18]. At this step, it has not been resolved whether NSP-vDNA is free in the cytosol or is transported via vesicles at their protoplasmic leaflet to the periphery. Consistent with this transport function, the cytosolic NIG (i) accumulates around the nuclear envelope, (ii) promotes the translocation of NSP-vDNA from the nucleus to the cytoplasm, (iii) functions as a pro-viral factor during begomovirus infection and (iv) shares structural features and transport properties with the human Rev-Interacting Protein (hRIP), which has been shown to be involved in the release of HIV-1 RNAs from the nuclear periphery to the cytoplasm [12, 18, 19]. Histone H3 has been shown to bind NSP and MP from BDMV as an additional component of the vDNA-protein complex that traffics intracellularly and intercellularly [20]. A nuclear acetyltransferase (AtNSI) interacts with the *Cabbage leaf curl virus* (CabLCV) NSP and may regulate the nuclear export of ss-vDNA-NSP via acetylation of CP and histones [21, 22]. Other NSP-interacting partners include the receptor-like kinases (RLKs) NSP-interacting kinase (NIK1) and NSP-associated kinase (NsAK), which are not involved in intracellular movement of vDNA but rather are implicated in antiviral immunity against begomoviruses [23-26].

For the cell-to-cell transport of vDNA, two distinct mechanisms, which basically differ in the nature of the vDNA-protein complex translocated into adjacent cells, have been suggested [11, 27]. In the first mechanism, designated “relay race” model, NSP facilitates the translocation of viral DNA from the nucleus to the cytoplasm and then is replaced by MP, which promotes the cell-to-cell movement of vDNA via plasmodesmata [10, 14]. The second mechanism represented by the “couple-skating” model preconizes that MP facilitates the vDNA-NSP nucleocytoplasmic trafficking and promotes the translocation of vDNA-NSP complex into the adjacent cells via endoplasmic-reticulum-derived tubules induced by the viral infection [11-13, 18, 28-32]. More recently, an alternative route for intracellular translocation and cell-to-cell spread of vDNA via chloroplasts and stromules has been proposed, which was based on experimental data of AbMV MP interactions (33,34). AbMV infection has been shown to induce the formation of a stromule network interconnecting different chloroplasts, chloroplasts with nuclei, and expanding to the cell periphery presumably at plasmodesmata [34]. This stromule network is stabilized by the plastid chaperone cpHSC70, which has also been shown to interact with MP [33]. The interaction between MP and cpHSC70 may bridge viral nucleoprotein complexes with stromules. Independent on the cell-to-cell transport models, the underlying mechanisms of the intracytoplasmic retrograde and anterograde movements of vDNA complexes remain to be determined. CabLCV and SLCV MPs have been shown to interact with synaptotagmin A (SYTA), which is located in the endosomes of plant cells and is required for the cell-to-cell trafficking of different MPs [35]. These findings raised the hypothesis that distinct viral MPs promote an intracytoplasmic transport of vDNA-complexes to plasmodesmata via an endocytic recycling pathway. Likewise, the AbMV MP has been shown to interfere with microtubule assembly and stability when co-expressed in plant tissues with a viral inducer of cell progression, which may suggest a microtubule-associated transit of MP or begomoviruses to the plasmodesmata [36]. However, the host components of the vDNA complex competent for intracytoplasmic translocation have not been identified, and the underlying mechanism for intracytoplasmic trafficking of vDNA remains to be elucidated.

In this investigation, we used a protein microarray assay to identify MP- and NSP-host protein-protein interactions (PPIs). From the NSP-MP-host PPI network map, we selected an intracellular transport-associated protein, designated NSP-interacting syntaxin domain-containing protein (NISP), as an NSP specific target, to gain further insight into the intracytoplasmic transport of begomoviruses. We showed that NISP, which was located in motile vesicle-like structures, many of which were targeted to the endosomes, interacted with NSP *in vivo* and displayed a pro-viral function. Furthermore, NISP also interacted with NIG, and the complex formed was enhanced by the presence of viral NSP. In addition, vDNA might have been recruited into the NISP complex. Our data indicate that NISP might be part of the vDNA complex that traffics intracellularly via endosomes.

## Results

### Identification of Arabidopsis proteins interacting with *in vitro* transcribed and translated MP and NSP

To search for the begomovirus movement proteins-Arabidopsis protein-protein interactions (PPIs), we used a nucleic acid programmable protein array (NAPPA)-based approach [37], in which we interrogated 4600 *in vitro* synthesized ORFs from Arabidopsis for their capacity to be targeted by MP and/or NSP from CabLCV. The *in situ* synthesized NAPPA protein microarray has previously generated a transcription factor (TF)-NAPPA dataset, which has been validated by pull-down assays and bimolecular fluorescence complementation (BiFC) assays in *N. benthamiana* leaves using a subset of newly identified interactions [37]. NSP and MP were *in vitro* transcribed and translated to probe this protein microarray (S1a and S1b Fig). A total of 40 candidate proteins were identified each for MP and NSP. The majority (35 proteins of 40) were found to interact with both proteins, which was not surprising because MP itself interacts with NSP and may be part of common multiprotein complexes as both participate in viral DNA movement (S2 Fig). We found five new NSP-specific protein interactions (dark orange) and five MP-specific protein interactions (light yellow), including a peptidyl-prolyl cis-trans isomerase family protein (AT1G73655) belonging to the same protein family as Pin4, which has been shown previously to interact with MP (S1 Table) [36]. As previously identified NSP-interacting proteins were absent in the protein array, the identification of known MP-specific interacting protein contributed to validate our current data.

The protein-protein interactions between the viral proteins and the Arabidopsis proteins were integrated into the protein-protein interactome experimentally determined for *Arabidopsis thaliana* (BioGRID database and the Arabidopsis interactome database), using Cytoscape software. This procedure identified the PPI network containing MP, NSP and all, directly and indirectly, interacting host proteins (Fig 1). AT1G68185, an ubiquitin-like protein, which formed a large hub (hub 20, degree 148), may represent a convergent targeting node among begomovirus viral proteins as it was found to be a direct target of NSP and MP independently (S1 Table). Furthermore, the ubiquitin-proteasome pathway (UPS) is targeted by many viruses to maintain suitable levels of viral proteins and to induce, inhibit or modify ubiquitin (Ub)-related host proteins [38]. Other MP-interacting proteins, including FKBP-like peptidyl-prolyl cis-trans isomerase family protein (this work), chloroplast heat shock protein 70-1 [33] and synaptotagmin A [35], which have been shown to assist the MP movement function, are connected to the AT1G68185 hub (hub 20), further supporting the notion that this hub may be biologically significant during begomovirus infection.

**Fig 1.**
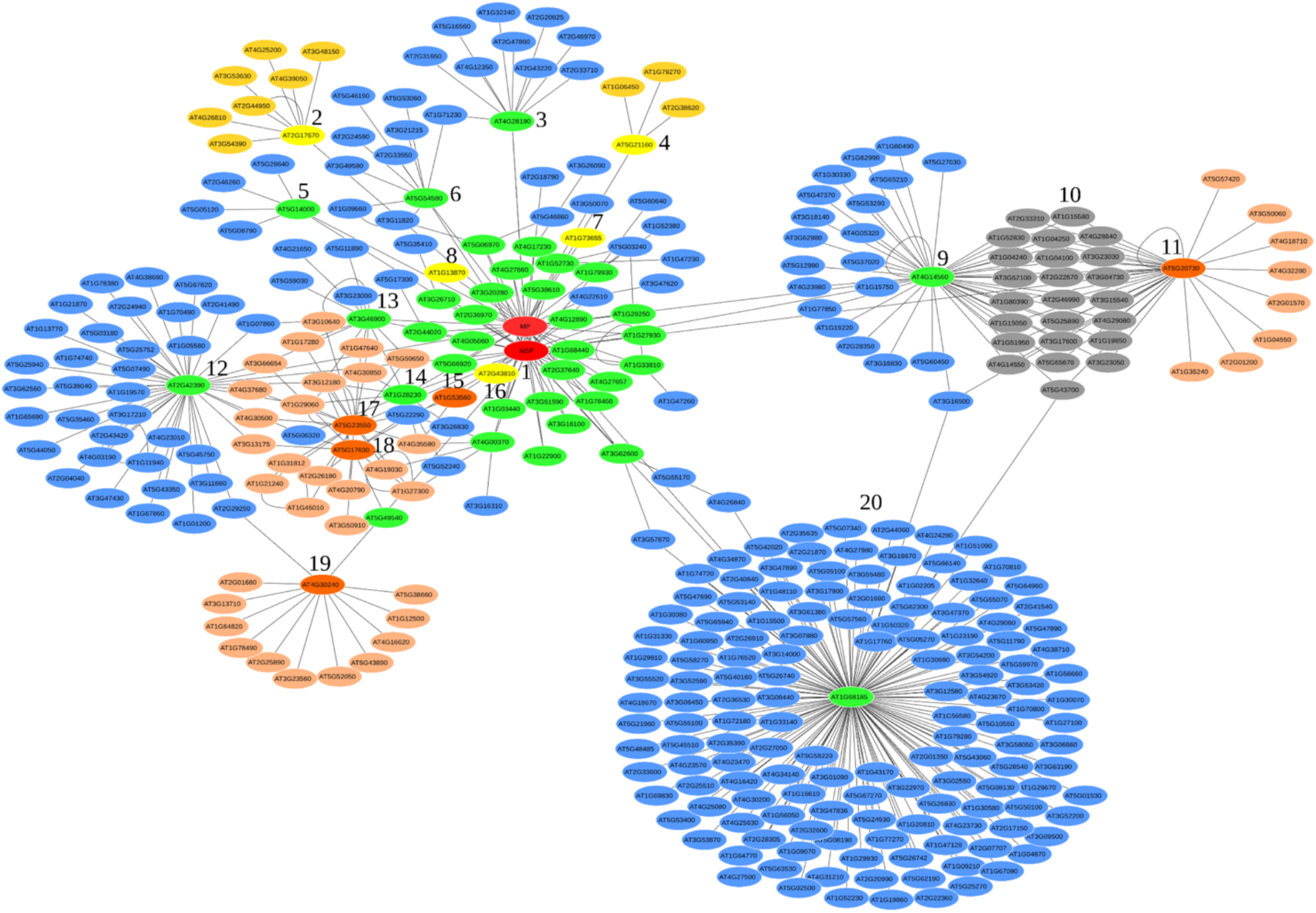
MP-NSP-Arabidopsis protein-protein interactions (PPI) network integrated into Arabidopsis interactome. The network displays a firework topology, which was assembled by the Cytoscape software. The viral proteins, MP and NSP, are represented in red. The NSP-specifically interacting proteins are in dark orange and connecting proteins from Arabidopsis in light orange. The MP-specifically interacting proteins are indicated in light yellow to which interacting dark yellow proteins converge. Arabidopsis proteins that interact with both MP and NSP are shown in green, forming hubs of convergent blue proteins. Gray proteins represent connecting points between two functionally similar hubs.

To gain further insights into the cellular processes affected by NSP or MP, we performed functional enrichment analyses of their direct and indirect interactors. Significantly enriched GO terms (with *p*-value < 0.05) were identified in all three categories, Biological Process, Molecular Function, or Cellular Component ontology (S2 Table). Some over-represented GO terms in this set of viral protein-interacting host proteins are consistent with NSP and MP function or localization. Consistent with their function as viral movement proteins, a subset of host interactors was significantly over-represented under the transport activity term and protein binding term in Molecular Function (S2 Table). Under the cellular component ontology, we observed an over-representation of proteins under membrane-bounded organelle term, intracellular vesicle term, and SNARE complex term, which may suggest an intracytoplasmic route for begomovirus proteins and vDNA trafficking (S2 Table). Accordingly, two NSP-interacting proteins, AT5G23550 and AT4G30240, form convergent hubs (17 and 19) enriched for membrane-bound proteins and transport activity function (Fig 1, S1 Table). The NSP-interacting protein AT5G23550 (hub 17) is annotated as a Got1/Sft2-like vesicle transport protein, which may be located to a late-Golgi compartment and may be involved in the fusion of retrograde transport vesicles derived from an endocytic compartment with the Golgi complex [39]. Likewise, AT4G30240 (hub 19) is a soluble N-ethylmaleimide-sensitive factor attachment protein receptor (SNARE)-like protein harboring a syntaxin-6 domain typical of proteins found in the endosomal transport vesicles [40].

Under the biological process ontology, the response to auxin term represented a significantly enriched GO term, which has not been previously associated with begomovirus infection (S2 Table). The NSP-MP-host PPI network uncovered two relevant hubs enriched for auxin response-related proteins, which are represented by an NSP-MP general interaction-derived hub (Fig 1, hub 9) and an NSP-specific host interaction-derived hub (hub 11). Remarkably, these two viral protein hubs are interconnected to each other via interactions with proteins (in gray), which are also over-represented under the auxin response term, forming a large auxin affected hub target by begomovirus (S1 Table). This hub may represent an undocumented biological process affected by begomovirus infection. The resulting PPI network map may provide a framework for future studies in detailing the begomovirus complex life cycle. One such example is described here, as following.

### A SNARE-like protein designated NSP-Interacting Syntaxin domain-containing Protein (NISP), but not its paralog, interacts with NSP via the syntaxin domain

The NSP-MP-Host PPI network uncovered enriched GO terms linked to intracellular vesicle and membrane-bound proteins, which may be associated with a mechanism for intracytoplasmic translocation of viral proteins and vDNA. To pursue further with this possible connection and to validate further the microarray data, we selected a new specific NSP-interacting protein, the SNARE-like protein AT4G30240, which harbors a syntaxin-6 domain at the N-terminus and a transmembrane segment at the C-terminus, for further analyses (Fig 2a). SNARE-complexes drive the fusion of membrane-bound vesicles with target membranes required for intracellular transport via the endosome system. We used the syntaxin domain superfamily as the prototype for the identification of homologs in the genomes of *Arabidopsis thaliana, Oryza sativa, Drosophila melanogaster, Homo sapiens*, and *Saccharomyces cerevisiae*. We then selected the most related proteins to construct phylogenetic trees using Bayesian inference method (S3a Fig). AT4G30240 was clustered in pair with AT2G18860 and formed a clade of syntaxin-6 domain-containing proteins, which also included a third Arabidopsis paralog AT1G27700. This AT4G30240-derived clade (in red), also designated Syntaxin-6-like subgroup, clustered only plant proteins, carrying an N-terminal syntaxin-6-like domain and a C-terminal transmembrane segment. AT4G30240 and AT2G18860 may be a result of genome duplication, as these syntaxin-6-like proteins clustered in pair even in phylogenetic analyses that include proteins from other organisms, share a high degree of sequence conservation and display similar intron/genomic DNA configuration (https://www.arabidopsis.org/).

**Fig 2.**
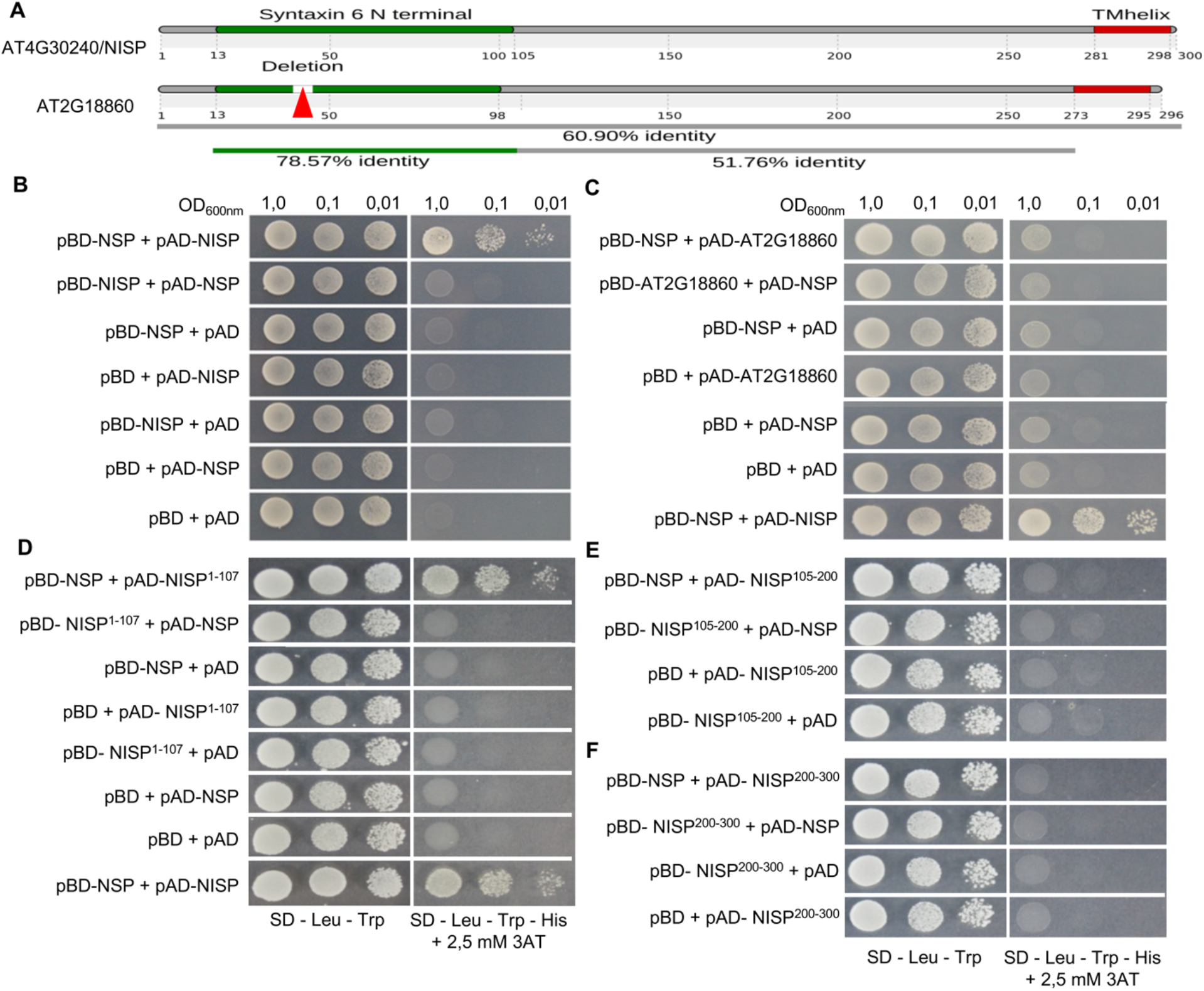
NISP, but not AT2G18860, interacts with NSP via the Syntaxin-6 domain. **(A)** Schematic representation of NISP and AT2G18860 sequence alignment. The syntaxin-6 domain is indicated in green and the transmembrane segment, in red. The identity of sequence and positions of amino acid residues are indicated. Sequence alignment was performed with CLUSTAL OMEGA. **(B)** NISP and NSP interact in yeast. NISP and NSP fused to the activation domain (AD), and binding domain (BD) of Gal4 and also in combination with empty vectors were expressed in yeast. The interactions between the tested proteins were monitored by histidine prototrophy in medium supplemented with 3-AT for 4 days at 28°C. **(C)** AT2G18860 failed to interact with NSP in yeast. AT2G18860 and NSP fused to both Gal4AD and BD were expressed in yeast, and possible interactions were assayed in all combinations. Co-expressed NSP-BD and NISP-AD were used as a positive control. **(D-F)**. Mapping the NSP-binding domain on NISP. Yeast cells expressing the indicated truncated version of NISP and NSP fusions were plated on selective medium lacking leucine, tryptophan, and histidine and supplemented with 2.5 mM 3-AT and grown for 4 days at 28°C. The truncated versions of NISP are delimited by the indicated position of the amino acid residues in the full-length protein. Protein interaction was only observed by co-expressing AD-NISP^1-107^ and BD-NSP.

The remaining syntaxin-6-like proteins clustered together in a distinct clade that included proteins from human, rice and the Arabidopsis AT1G28490, also named SYP61, a characterized trans-Golgi network (TGN)/endosome localized syntaxin [41-43]. This SYP61 second clade (in orange) was designated target (t)-syntaxin-6/SNARE-like subgroup, as it included structurally similar proteins carrying the N-terminal syntaxin-6 domain and also a t-SNARE coiled-coil homology domain at the C-terminus.

However, the syntaxin-6-like subgroup represents a newplant-specific class in the t-SNARE superfamily, since it lacks the coiled-coil t-SNARE domain at the C-terminus, which is responsible for vesicle fusions. A function of this subgroup has to be determined, yet. This hypothesis was further confirmed in a phylogenetic analysis of syntaxin-6 domain-containing proteins from *Arabidopsis thaliana, Oriza sativa, Glycine max, Zea mays, Zostera marina, Homo sapiens, Mus musculus, Bos taurus, Equus caballus, Danio rerio*, and *Drosophila melanogaster* using the full-length sequence of AT4G30240 as prototype (S3b Fig). The syntaxin-6 domain-containing proteins form two distinct clades with high internal stability (100%). The AT4G30240-derived clade, syntaxin-6-like clade, includes only plant syntaxin-6-like proteins.

To complement the protein microarray data, we used the yeast two-hybrid system to examine further the NSP-AT4G30240 interaction and compared the paralog AT2G18860 in our assays, which was not present in the protein microarray. The second most closely related paralog, AT1G28490, was present in the protein microarray but was not found to interact with NSP. Co-expression of the full-length AT4G30240 (NISP) ORF fused to the activation domain (AD), and NSP fused to the binding domain (BD) of GAL4 promoted histidine prototrophy (Fig 2b). The HIS3 marker gene was not activated in yeast cells cotransformed with the controls. In contrast, AT2G18860 failed to interact with NSP in yeast, as neither AD-AT2G18860 nor BD-AT2G18860 promoted his prototrophy when cotransformed with the respective cognate BD-NSP or AD-NSP (Fig 2c).

To map the NSP-interacting domain on NISP, we generated NISP deletion fragments and assay for protein interactions by two-hybrid assay. The N-terminal NISP fragment extending up to amino acid position 107 and containing an intact syntaxin-6 domain (NISP^1-107^, Fig 2a) was sufficient to bind NSP in yeast (Fig 2d). The interaction of the truncated NISP^1-107^ with NSP was specific because the HIS marker gene was not activated in yeast cells co-transformed with AD- or BD-truncated NISP^1-107^ and the empty vectors (Fig 2d). In contrast, the NISP truncated versions, comprising the NISP internal region, delimited by positions 105-200 (NISP^105-200^; Fig 2e) or the C-terminal portion (NISP^200-300^; Fig 2f), were not capable of recognizing NSP, confirming that NISP binds NSP via its syntaxin-6 domain. Although the syntaxin-6 domains of NISP and AT2G18860 are conserved (78.57% identity), NSP interacts specifically with NISP in yeast (Fig 2b). Sequence alignment revealed an 8-amino acid deletion at the syntaxin-6 domain of AT2G18860 as the most striking feature that may account for the differences in protein binding specificity (S4 Fig).

Although NISP lacks a typical coiled-coil t-SNARE domain, it harbors a syntaxin-6 domain at the N-terminus and a predicted transmembrane segment at the C-terminus, which suggests an endomembrane-bound localization (Fig 2a). Mammalian syntaxin-6 domain proteins are localized in the trans-Golgi network (TGN) membranes and endosomal vesicles [44]. To examine the subcellular localization of NISP, we transiently expressed AT4G30240 fused to a green fluorescent protein (GFP) under the control of the native and constitutive promoters, in *Nicotiana benthamiana* leaves and analyzed the localization of the fluorescent signal by confocal microscopy. NISP-GFP was predominantly localized in motile spherical structure, resembling vesicles (Fig 3a, 3b; S1 Video). The paralog AT2G18860 fused to GFP appeared concentrated predominantly in membrane-bound vesicles and plasma membrane (S5 Fig).

**Fig 3.**
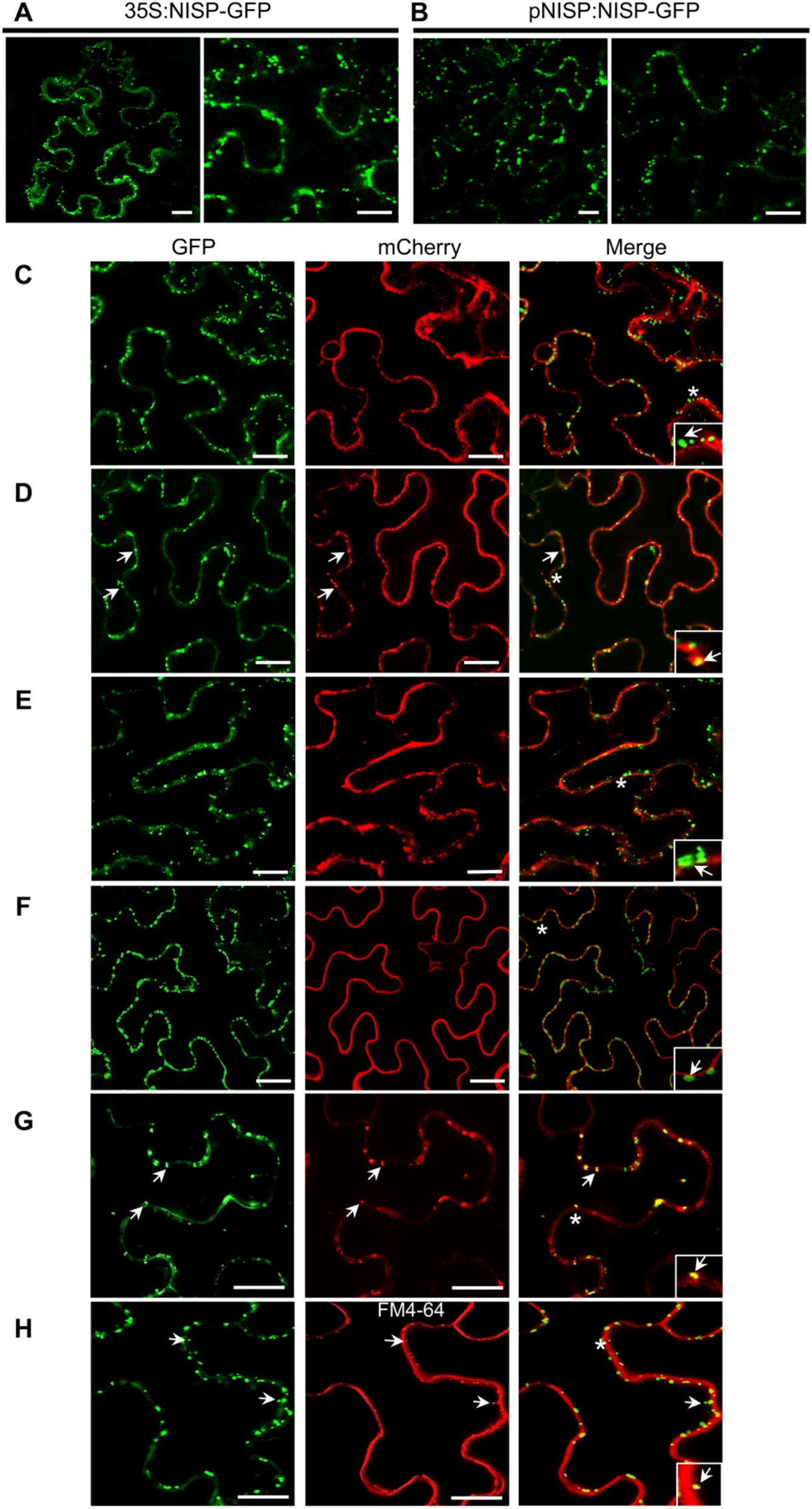
Subcellular localization of NISP. **(A)** The NISP-GFP distribution in intracellular vesicles. *N. benthamiana* leaves were infiltrated with *A. tumefaciens* carrying a DNA construct expressing NISP-GFP under the control of 35S promoter. Confocal images from three independent experiments were taken 36 hours post-infiltration. Approximately 200 cells were examined. Scale bars, 10 μm and 20 μm. **(B)** Confocal images of NISP-GFP transiently expressed under the control of the endogenous promoter. Scale bars, 10 μm and 20 μm. **(C)** Co-expression of the ER-associated ER-mCherry molecular marker and NISP-GFP in *N. benthamiana* leaves. *N. benthamiana* leaf cells expressing the indicated fusion proteins were examined by confocal microscopy 2 days after infiltration. The asterisks indicate the localization of the amplified insets, and the arrows indicate the merged pattern of GFP-associated vesicles in C, D, E, F, G, H. The confocal images are representative samples from three independent experiments. Scale bars, 10 μm. **(D)** Co-expression of the Golgi-associated G-mCherry molecular marker and NISP-GFP in *N. benthamiana* leaves 2 days after agro-infiltration. Scale bars, 20 μm. **(E)** Co-expression of the tonoplast-associated vac-mCherry molecular marker and NISP-GFP in *N. benthamiana* leaves 2 days after agro-infiltration. Scale bars, 20 μm. **(F)** Co-expression of the plasma membrane-associated FLS2-mCherry and NISP-GFP in *N. benthamiana* leaves 2 days after agro-infiltration. Scale bars, 20 μm. **(G)** Co-expression of endosome-associated SYTA-mCherry and NISP-GFP in *N. benthamiana* leaves 2 days after agro-infiltration. Scale bars, 10 μm. **(H)** Co-labeling of vesicles with NISP-GFP and FM4-64 in epidermal cells of *N. benthamiana*. Images were taken after 2 days post infiltration of *Agrobacterium* carrying a NISP-GFP expression vector and 1 h after infiltration of FM4-64. Scale bars, 20 μm.

To examine more precisely the NISP localization, we performed co-localization assays, using SYTA-mCherry as an endosome-associated marker [35], the plasma membrane-associated molecular marker FLS2-mCherry [45], ER-, TGN- and tonoplast-associated molecular markers constructed by Nelson et al. [46]. NISP-GFP did not co-localize with ER-mCherry-(Fig 3c, asterisks indicate the field of the insets) and tonoplast (vac-mCherry)-(Fig 3e) associated molecular markers. Likewise, NISP-containing vesicles did not co-localize with the plasma membrane FLS2-mCherry (Fig 3f). In contrast, a small population of NISP-GFP-containing vesicles co-localized with the TGN-associated molecular marker (Fig 3d) and a large fraction of NISP-GFP-containing vesicles precisely co-localized with the endosomal protein SYTA-mCherry (Fig 3g). Because many of the NISP-labeled vesicles were associated with the plasma membrane, we examined whether the NISP-associated vesicle membrane was derived from the plasma membrane by staining NISP-GFP-expressing cells with FM4-64, a marker for plasma membrane internalization that labels endosomes [47]. In addition to dye-labeled plasma membranes, some vesicles showed also NISP-GFP fluorescence (Fig 3H, arrows). Collectively, these results indicate that NISP localizes in TGN/early endosome vesicles.

### NISP interacts with NSP in intracellular vesicles

We next examined whether NISP interacts with NSP *in vivo* using the bimolecular fluorescence complementation (BiFC) assay. The formation of NISP/NSP complexes occurred atvesicles of *N. benthamiana* epidermal cells independent of the orientation of the NISP fusions (N-terminus or C-terminus of YFP; Fig 4a), and the reconstituted fluorescent signal was much higher than that of the background (control panels with combinations of the protein fusions with empty vectors). The reconstituted fluorescent signal was co-labeled by FM4-64, indicating that the complex was formed in endosomal vesicles derived from the plasma membrane (Fig 4b, see arrows). We further demonstrated that NISP and NSP interacted *in vivo* using co-immunoprecipitation assays. Both NISP-GFP and NSP-6HA, were co-immunoprecipitated from co-infiltrated extracts using anti-GFP antibody, demonstrating an association of NISP-GFP with NSP-6HA fusions (Fig 4c). Switching the protein tags and including MP did not alter the results because NISP-6HA also associated with NSP-GFP in co-infiltrated extracts (Fig 4d). Co-expression of MP in the assay was used to mimic infection, since CabLCV MP promotes the directionality of NSP to the cell periphery (12, 13). The Co-IP results also demonstrated that GFP alone did not interact with either NSP-6HA or NISP-6HA, demonstrating the specificity of the interactions between the viral protein and NISP.

**Fig 4.**
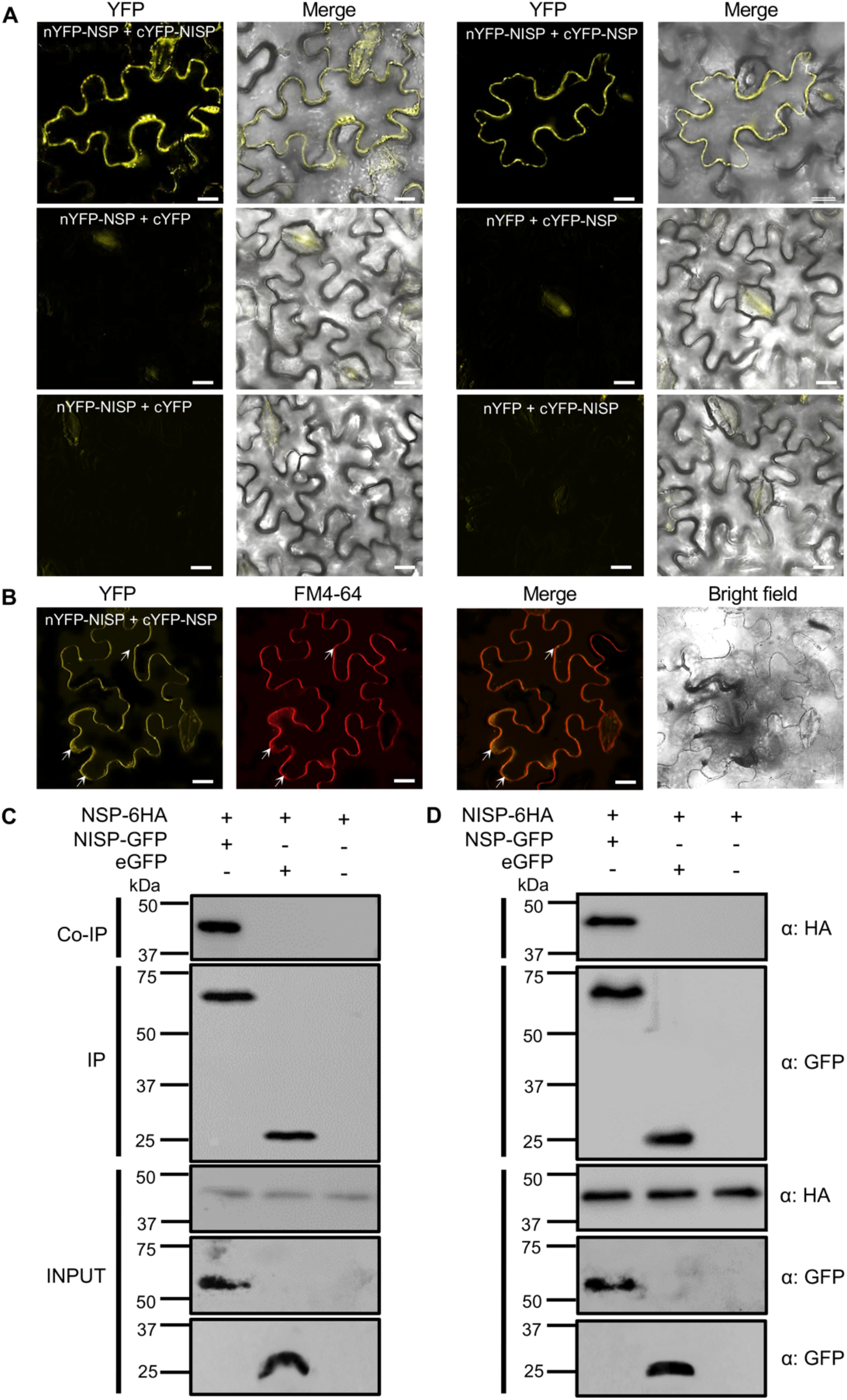
NISP interacts with NSP *in vivo*. **(A)** NISP interacts with NSP in vesicle-like compartments, as judged by BiFC. *N. benthamiana* leaf cells expressing NISP and NSP fused to the C-terminus (cYFP) or N-terminus (nYFP) of YFP, or in combination with the empty vectors, were observed by confocal microscopy 3 days after infiltration. Scale bars, 10 μm. The experiment was repeated at least four times with identical results. **(B)** NISP-NSP complex-containing vesicles are stained by FM4-64. *N. benthamiana* leaves were agroinfiltrated with nYFP-NISP- and cYFP-NSP-expressing DNA constructs. Images were taken 2 days post agroinfiltration and 30 min after infiltration of FM4-64. Arrows indicate examples of vesicle-associated reconstituted fluorescent signals that were co-stained by FM4-64. Scale bars, 10 μm. The experiment was repeated three times with similar results. **(C)** NISP interacts with NSP *in planta*. Whole cell protein extracts from N. benthamiana leaves expressing transiently NISP-GFP and NSP-6HA were used for co-immunoprecipitation assays using an anti-GFP serum. Input and IP show the levels of the expressed proteins NSP-6HA, NISP-GFP, and GFP. Anti-HA antibody was used to detect NSP-6HA from the immunoprecipitated complex. GFP was used as a negative control. Molecular mass markers are given on the left in kilodaltons. The experiment was repeated twice. **(D)** *In vivo* interaction of NISP with NSP. Co-immunoprecipitation was performed as described in C, except that the tags of the transiently expressed proteins were switched to NSP-GFP and NISP-6HA. The experiment was repeated twice.

### NISP, but not its paralog AT2G18860, displays a pro-viral function

The biological relevance of the NSP-NISP interactions in begomovirus infection was examined using several different approaches. We first analyzed the expression of NISP from publically available RNA-seq data, which showed that NISP was induced by begomovirus infection [26]. Quantitative RT-PCRs showed that NISP RNAs, but not the RNA from the NISP paralog AT2G18860, were induced by the begomovirus infection (S6 Fig). We then identified a T-DNA insertion mutant in the NISP gene (*nisp-1*) and the AT2G18860 gene (*at2g18860-1*). The *NISP* or AT2G18860 transcript levels in the respective T-DNA insertion mutant lines were much lower than in Col-0, indicating that *nisp-1* and *at2g18860-1* may be loss-of-function mutant lines (Fig 5a, 5c). These mutant lines were transformed with *NISP-GFP* or AT2G18860-GFP constructs to analyze a complementation of the defects in three independent lines each (Fig 5a, 5b, *nisp-1*/NISP lines, and *at2g18860-1/AT2G18860* lines). We also prepared transgenic lines overexpressing *NISP-GFP* and AT2G18860-GFP, under the control of the 35S constitutive promoter. The respective transcript (Fig 5a, 5c) levels were higher in NISP and AT2G18860 plants than in Col-0 and accumulation of the encoding proteins was determined by immunoblotting with an anti-GFP antibody (Fig 5b, 5d). The knockout and overexpressing lines were phenotypically indistinguishable from Col-0 (S8 Fig).

**Fig 5.**
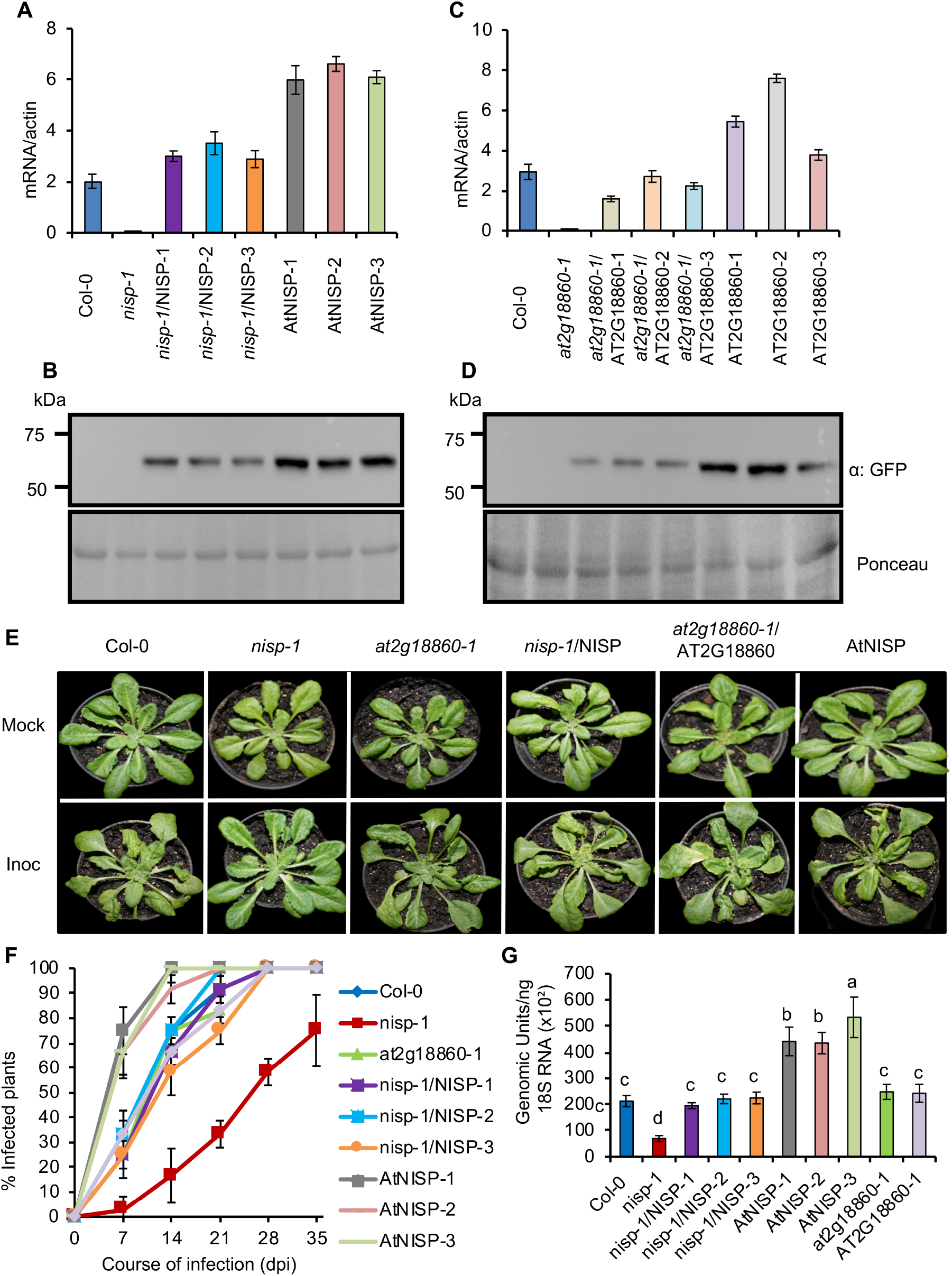
NISP displays a pro-begomoviral function. **(A)** Transcript accumulation of NISP in transgenic lines. NISP transcript levels in three independently complemented transgenic lines (*nisp-1*/NISP-1-3), three independently transformed *NISP*-overexpressing lines (AtNISP1-3), Col-0 and *nisp-1* were determined by qRT-PCR. Gene expression was calculated using the 2^-ΔCt^ method, and actin was used as an endogenous control. Error bars represent 95% confidence intervals based on replicated samples (n=3) from three independent experiments. **(B)** NISP-GFP protein accumulation in transgenic lines. Whole cell protein extracts from the complemented transgenic lines (*nisp-1*/NISP), *NISP*-overexpressing lines (AtNISP), Col-0 and *nisp-1* were separated by SDS-PAGE and immunoblotted using an anti-GFP serum. **(C)** AT2G18860 transcript accumulation in transgenic lines. The transcript levels of AT2G18860 in complemented transgenic lines (*at2g18860-1*/AT2G18860-1-3), AT2G18860-overexpressing lines (AT2G18860-1-3), Col-0 and *at2g18860-1* were quantified by qRT-PCR and normalized to the actin as an endogenous control. Error bars indicate 95% confidence intervals based on replicated samples (n=3) from three independent experiments. **(D)** AT2G18860-GFP protein accumulation in transgenic lines. Total protein extracts from complemented transgenic lines (*at2g18860-1*/AT2G18860), AT2G18860-overexpressing lines (AT2G18860), Col-0 and *at2g18860-1* were fractionated by SDS-PAGE and immunoblotted using anti-GFP antibody. **(E)** CabLCV infection-associated symptoms in Arabidopsis genotypes at 21 days post-inoculation (dpi). The figure shows representative samples of mock-inoculated and CabLCV -infected plants. The genotypes are indicated in the figure and are the same as specified in A-D. **(F)** The onset of infection is delayed in *nisp-1* knockout line. Ecotype Col-0, *nisp-1, at2g18860-1, nisp-1*/NISP complemented lines and AtNISP-overexpressing lines (as in a-d) were infected with CabLCV DNA, and the course of infection was monitored as the percent of systemically infected plants at different dpi. **(G)** Absolute quantification of CabLCV genomic units in infected lines at 21 dpi. The genotypes are the same as specified in a-d. Error bars indicate 95% confidence intervals based on replicated samples from three independent experiments. Different letters indicate significative differences among the groups (Duncan test, p <0,05, n= 3).

We next examined whether *NISP* and AT2G18860 participate in begomovirus infection. The knockout lines (*nisp-1* and *at2g18860-1)*, complemented lines (*nisp-1/NISP* and *at2g18860-1/AT2G18860), NISP*-overexpressing lines and Col-0 were inoculated with infectious clones of CabLCV and the course of infection was monitored by visualization of symptom development and PCR diagnosis. The accumulation of viral DNA was quantified by qPCR (Fig 5).

The *nisp-1* knockout line displayed attenuated symptoms (Fig 5e, S7 Fig), delayed course of infection (Fig 5f), and lower viral DNA as compared to Col-0 (Fig 5g). Expression of NISP-GFP in *nisp-1* restored the wild type susceptibility to begomovirus infection in all three independently transformed lines (*nisp-1/NISP*), confirming that the reduced susceptibility displayed by the *nisp-1* line was indeed due to loss of *NISP* function. In contrast, *NISP* overexpression conferred enhanced susceptibility to begomovirus infection; the progress of infection was accelerated (Fig 5f), and viral DNA load was significantly higher in the *NISP*-overexpressing lines than in Col-0 and *nisp-1* lines (Fig 5g). We predicted that if NISP-NSP binding was the basis for the NISP pro-viral function, the closest paralog of NISP, AT2G18860, which harbors the syntaxin-6-like domain but does not interact with NSP, would not affect begomovirus infection. Accordingly, the *at2g18860-1* knockout line, *at2g18860-1/AT2G18860* complemented lines, and an *AT2G18860*-overexpressing line displayed similar symptoms, course of infection and viral DNA load as Col-0, indicating that the presence or absence of the *AT2G18860* gene does not change the begomovirus infection (Fig 5e, 5f, 5g, S7 Fig). These results further substantiated a specific pro-viral function of NISP rather than of syntaxin-6 like domain proteins in general.

### NISP also interacts with NIG and complex formation is enhanced by viral NSP

There is a gap in our knowledge of how the vDNA-NSP complex, which binds to NIG in the cytosolic side of the nuclear pores, traffics to the cell periphery. From the PPI network map, we hypothesized that NIG could serve as a bridge to attach vDNA to the intracellular transport system of plant cells. The integration of our data with the Arabidopsis interactome uncovered a possible interaction between NIG and NISP, which was also shown here to bind specifically to viral NSP. The interaction between NIG and NISP was first demonstrated by two-hybrid assay (S9a Fig). The co-expression of the NISP ORF fused to BD and NIG ORF fused to AD from GAL4 promoted histidine prototrophy in medium supplemented with 2.5 mM 3AT, whereas co-expression with empty vectors did not.

To examine whether NIG and NISP interact *in vivo*, we used BiFC and co-immunoprecipitation assays. NIG has been shown to be unevenly distributed around the nuclear envelope, and cell periphery (S9b and S9c Fig) [25, 48]. In the BiFC analyses, nYFP-NIG as well as cYFP-NISP (and vice-versa) were localized at the cell periphery (Fig 6a), probably at the NISP endosomes, since the structures were co-stained by FM4-64 (Fig 6b; arrows). The fluorescent signal was barely detected in the controls (Fig 6a). The inclusion of NSP-6HA in the BiFC assay increased the NISP-NIG complex formation, as judged by the enhanced intensity of the reconstituted fluorescent signal, showing the fluorescent vesicles associated with the plasma membrane (S10 Fig). To provide further evidence of the NSP-mediated enhancement of NIG-NISP interaction, we used a quantitative co-immunoprecipitation assay (Fig 6, S9 Fig). We first confirmed that NIG interacts with NISP *in vivo* (Fig 6c, S10d). The Co-IP results indicated that NISP-GFP, but not GFP alone, interacted with NIG-HA. Likewise, NIG-GFP, but not GFP alone, interacted with NISP-HA. NSP promoted a stronger association of NISP-GFP with NIG-6HA, as judged by the amount of co-immunoprecipitated NIG by anti-GFP in the presence of co-expressed GST-NSP (Fig 6d, 6e). Likewise, GST-NSP enhanced NISP-6HA-NIG-GFP complex formation (S9e and S9f Fig). These results indicate that NSP/NIG and NISP might form a multiprotein complex at endosomes. Syntaxin-6 domain-containing proteins have been shown to participate in the endosome-plasma membrane route of protein cargos.

**Fig 6.**
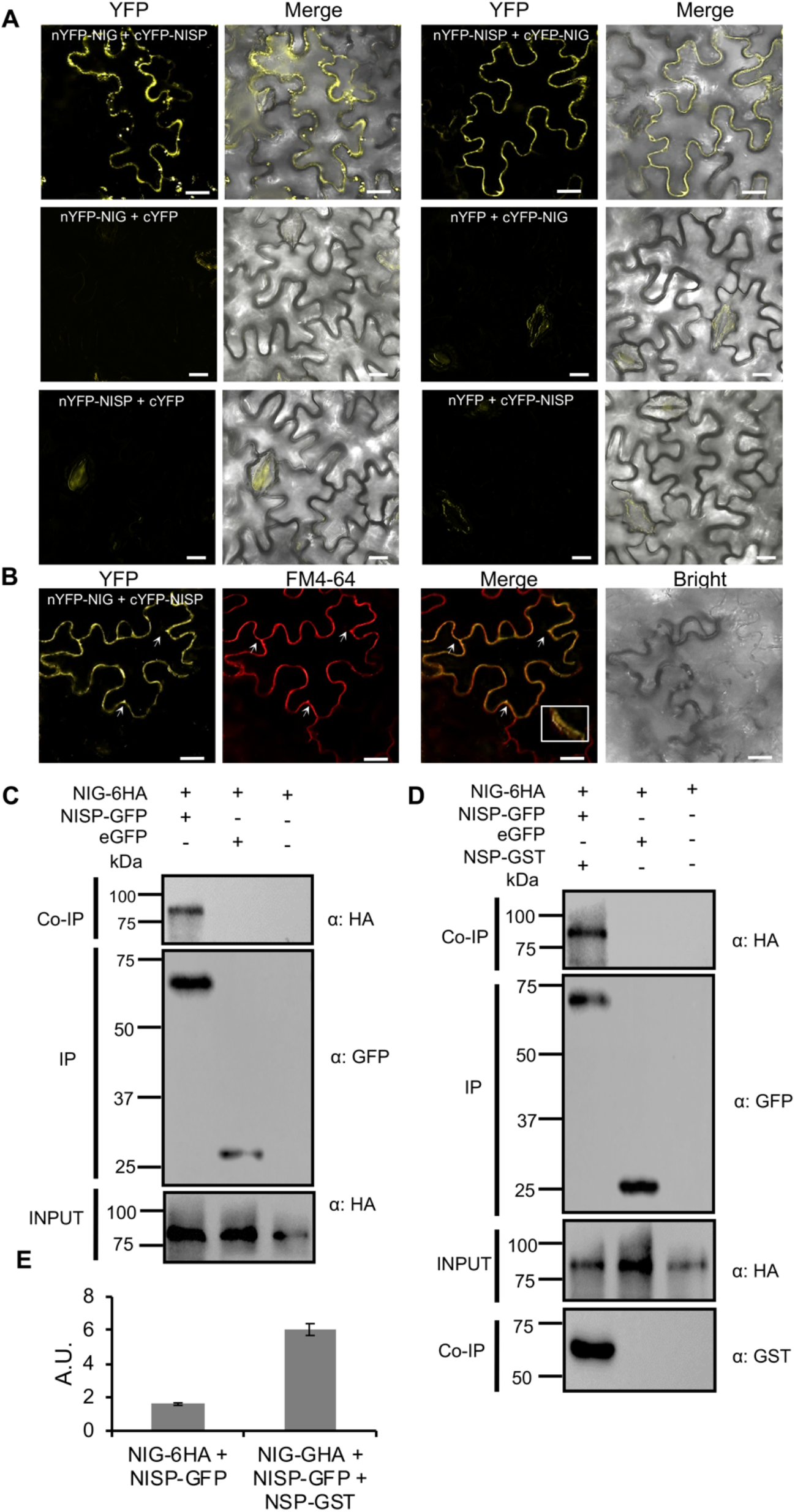
NISP interacts with NIG *in vivo* and NSP enhances NISP-NIG complex formation. **(A)** BIFC assay showing the interaction between NISP and NIG in vesicles of *N. benthamiana* leaf cells. *N. benthamiana* leaves transiently expressing NISP and NIG fused to the C-terminus (cYFP) or N-terminus (nYFP) of YFP were examined by confocal microscopy 3 days after infiltration. Scale bars, 10 μm and 20 μm. The experiment was repeated three times with similar results. **(B)** NISP-NIG complex-containing vesicles are stained by FM4-64. *N. benthamiana* leaves were agroinfiltrated with nYFP-NIG- and cYFP-NISP-expressing DNA constructs. Images were taken 2 days post infiltration and 1 h after infiltration of FM4-64. Arrows indicate examples of vesicle-associated reconstituted fluorescent signals that were co-stained by FM4-64 and the asterisk, the region of the amplified inset. Scale bars, 10 μm. The experiment was repeated three times with identical results. **(C)** In vivo interaction of NISP and NIG. Total protein extracts from *N. benthamiana* expressing NISP-GFP and NIG-6HA were used for co-immunoprecipitation assays using anti-GFP. Input and IP show the levels of the expressed proteins NISP-GFP and NIG-6HA. Anti-HA was used to detect NIG-6HA from the immunoprecipitated complex. GFP was used as an unrelated protein. The experiment was repeated twice with identical results. **(D)** NISP-NIG complex formation in the presence of viral NSP. The Co-IP assay was performed as described in A, except that co-expressed NSP-GST was included in the assay. NSP-GST was detected by immunoprecipitating it from infiltrated leaves and immunoblotting with anti-GST. The experiment was repeated twice with identical results. **(E)** Viral NSP enhances the interaction between NISP and NIG. NIG levels in the immunoprecipitated complex in the presence and absence of viral NSP were quantified using the Band Analysis tools of the ImageLab software (Bio-Rad). The signal values were normalized using the IP NISP-GFP band, revealed in the same blot as the Co-IP band. A.U. denotes arbitrary units.

### NISP complexes viral DNA in vivo

The current model of vDNA intracellular translocation informs that NSP binds to nascent viral DNA in the nucleus, facilitates its translocation to the cytosol via nuclear pores and moves vDNA intracytoplasmically to the cell periphery, where MP mediates the intercellular movement of vDNA-NSP or vDNA-MP via plasmodesmata. Because NSP also binds to NISP in vesicle-like structures, we asked whether the vDNA would be recruited into a NISP multi-protein complex using a modified chromatin immunoprecipitation (ChIP) assay (Fig 7). In our assay, proteins and DNA were first cross-linked with formaldehyde and then the protein-DNA complexes were immunoprecipitated with antibody (anti-GFP for NISP-GFP) from total protein extracts. The untransformed line (Col-0) and NISP-GFP-overexpressing (AtNISP-1, AtNISP-2, and AtNISP-3, Fig 7a) lines were infected with CabLCV infectious clone and the modified ChIP assay was performed using an anti-GFP antibody. We confirmed first that the inoculated plants were infected (IN) by performing PCR from input DNA using CabLCV-specific primers, which amplify a 770-bp fragment from CabLCV DNA-B (Fig 7b and S11b Fig, primers F1/R1) and a 389-bp fragment from DNA-A (Fig 7e and S11a Fig, primers F1/R1). This viral DNA-B and DNA-A fragments were also amplified from ChIP-DNA of infected (IN) AtNISP-overexpressing lines but not of infected (IN) or uninfected (UN) Col-0, which lacks GFP-tagged NISP (Fig 7a, 7e). Using two sets of DNA-B-specific primers (F2/R2 and F3/R3, S10b Fig), ChIP-qPCR further demonstrated significant enrichment of DNA-B-specific fragments in infected AtNISP-GFP-overexpressing samples, which had been precipitated with anti-GFP serum, but not in pulled down samples from infected Col-0 lines (Fig 7c and 7d). Likewise, the use of F2/R2 set of DNA-A primers (S10a Fig 10a) also resulted in significant enrichment of the DNA-A-specific fragment in ChIPed DNA from infected AtNISP-GFP-overexpressing samples but not from infected Col-0 (Fig 7f). Collectively, our results indicate that NISP is a functional component of the vDNA-complex that traffics along the cytoplasm to the cell periphery for the MP-mediated cell-to-cell movement of vDNA into adjacent cells.

**Fig 7.**
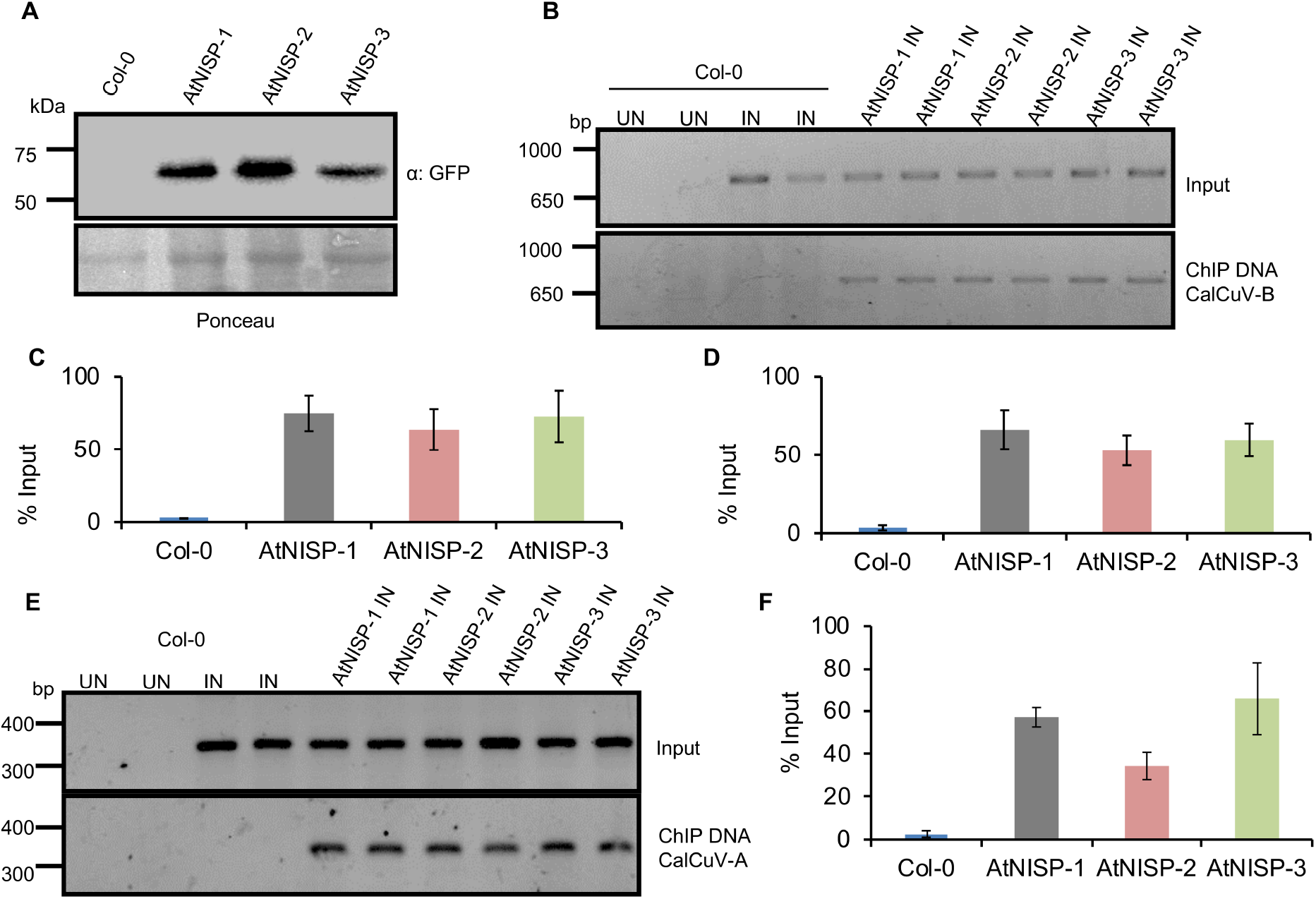
NISP complexes with vDNA *in vivo*. **(A)** NISP-GFP accumulation in overexpressing lines. Whole cell protein extracts from the NISP-GFP-overexpressing lines, as indicated in the figure, and Col-0 were separated by SDS-PAGE and immunoblotted using an anti-GFP serum. **(B)** NISP binds to viral DNA-B *in vivo*. NISP-GFP-overexpressing lines (AtNISP-1, AtNISP-2, and AtNISP-3), and Col-0 were inoculated with infectious CabLCV clones prior to the ChIP assay, which was performed with leaf samples of the indicated genotypes using anti-GFP. The infected plants were confirmed by PCR of input DNA from inoculated plants using CabLCV DNA-B-specific primers, which amplifies a 770-bp fragment (F1/R1; S10b Fig). IN denotes infected leaves and UN, uninfected leaves. **(C)** ChIP-qPCR assay of leaves from AtNISP-GFP transgenic lines and Col-0. The samples were immunoprecipitated with an anti-GFP antibody and ChIPed DNA was quantified by qPCR using the DNA-B-specific primers F3/R3 (S10b Fig). Data were normalized relative to the input of each sample and are expressed as the percentage of input. (D) ChIP-qPCR assay using a different set of DNA-B-specific primers (F2/R2) and samples from the indicated genotypes. The ChIP-qPCR was performed as described in C. **(E)** NISP binds to viral DNA-A *in vivo*. NISP-GFP-overexpressing lines (AtNISP-1, AtNISP-2 and AtNISP-3), and Col-0 were inoculated with infectious CabLCV clones prior to the ChIP assay, which was performed with leaf samples of the indicated genotypes using anti-GFP. The infected plants were confirmed by PCR of input DNA from inoculated plants using CabLCV DNA-A-specific primers, which amplifies a 389-bp fragment (F1/R1; S10a Fig). IN denotes infected leaves and UN, uninfected leaves. **(F)** ChIP-qPCR assay using a set of DNA-A-specific primers (F2/R2, S10a Fig) and samples from the indicated genotypes. The ChIP-qPCR was performed as described in C. The above experiments were repeated three times with similar results.

## Discussion

Viruses are obligate intracellular parasites and, once in the host cell cytoplasm, they must move intracellularly to and from the replication site to the plasma membrane for spreading. The cytoplasmic limitations on free diffusion have forced viruses to evolve efficient mechanisms to manipulate the host transport system for their benefit. Understanding how viruses hijack the host intracellular transport system using a limited repertoire of proteins provides an opportunity to uncover the molecular bases of the host transport machinery. Bipartite begomoviruses, plant ssDNA viruses, use two movement proteins, MP and NSP, to move intracellularly from the site of replication in the nucleus to the cell surface [17]. However, the host components of the vDNA complex competent for intracytoplasmic translocation have not been identified, and the underlying mechanism for intracytoplasmic trafficking of vDNA remains to be elucidated. Here, we used a previously fabricated, *in sit*u synthesized protein microarray containing 4600 Arabidopsis ORFs [37] to identify NSP- and MP-Arabidopsis protein-protein interactions (PPIs). Consistent with the movement function of the viral proteins, the identified NSP-MP-PPI network uncovered direct and indirect interactions, over-represented under transport activity, protein binding, membrane-bound organelles, intracellular vesicle, and SNARE complex ontology terms, which may define an intracellular route for vDNA trafficking. Precedents in the literature have implicated a microtubule-associated intracellular movement of viral proteins and vDNA in addition to an endocytic recycling pathway for the MP-mediated cell-to-cell movement of vDNA protein complexes (35; 36]. Accordingly, transport function-related proteins, including kinesin motor protein, plant syntaxin 121, Rab GTPase, Ras, Got1/Sft2-like vesicle transport protein, target SNARE coiled-coil domain protein, syntaxin-6 plant proteins, prenylated Rab acceptor, Heat shock protein 70 (Hsp 70), nuclear pore anchor, synaptotagmin A, were over-represented in the MP-NSP-PPI network. These studies thus provided a critical framework for future investigations to elucidate the molecular mechanisms for intracytoplasmic transport of begomoviruses.

### The NSP-interacting syntaxin domain-containing protein (NISP) is identified as a new functional cellular partner of begomovirus NSP

In the present investigation, we characterized an NSP-specifically interacting syntaxin-6 domain-containing protein, designated NISP and provided several lines of evidence indicating that NISP might be involved in the intracytoplasmic movement of NSP-vDNA. First, co-localization studies using organelle-associated molecular markers, and the FM4-64 marker demonstrated that NISP is associated with TGN/early endosomal vesicles. Second, we confirmed that NISP interacted with NSP *in planta* and showed that the complex formation occurred in vesicle-like structures, which resemble the NISP-associated TGN/endosomal vesicles, the typical subcellular localization of syntaxin-6-like proteins from Arabidopsis and humans [49]. Third, we showed that NISP exhibits a pro-viral function, and begomovirus infection required NISP-NSP interaction. The mutant *nisp-1* was less susceptible nt to begomovirus as the knockout lines displayed attenuated symptoms, a delayed course of infection, and accumulated much lower viral DNA as compared to Col-0. In contrast, the overexpressing lines were more susceptible to begomovirus. We took advantage of a highly conserved NISP paralog, AT2G18860, to show that NISP-NSP interaction was the molecular basis for the NISP pro-viral function. AT2G18860, which shares with NISP a 78,6% identical syntaxin-6 domain, did not interact with NSP and thus did not interfere with begomovirus infection. Consistent with these data, NISP, but not AT2G18860, is induced by begomovirus infection. Fourth, NISP also interacted with NIG, which facilitates the release of the NSP-DNA complex from the nuclear pores to the cytosol [12,18], and the presence of NSP enhanced the NISP-NIG complex formation, indicating that NSP, which interacts with NIG [12] and NISP (Fig 4), may be recruited by the complex. We used ChIP assay to demonstrate that NISP was also associated with vDNA in infected cells, which may be assembled into a NISP-NIG-NSP multiprotein complex. Finally, the NISP interactions relocated the dispersed cytosolic NIG and viral NSP to trafficking vesicle resembling the NISP-containing spherical structure associated with TGN/endosome. The NISP-mediated translocation of NSP-vDNA to endosomal vesicles favors the association of the viral complex with MP, which is brought to the endosome via interaction with the endosome-associated SYTA for the MP-mediated cell-to-cell movement of vDNA [35].

### NISP belongs to a new, previously unidentified plant-specific subfamily of syntaxin-6-like proteins

The phylogenetic reconstruction of syntaxin-6 domain-containing proteins from different eukaryotic organisms divided this family into two subclades, one containing proteins from all tested organisms, including SYP61 from Arabidopsis, and a new, previously unidentified plant-specific subclade, which includes NISP and AT2G18860 paralogs clustered in pair. SYP61 has been shown to form functional SNARE complexes involved in the vacuolar-TGN recycling pathway and in exocytotic trafficking to the plasma membrane [43,50]. The process of membrane fusion is the final step of vesicle transport, which is generally driven by a specific pairing of 1 v-SNAREs with the 2 or 3 cognate t-SNARE molecules into a trans-SNARE complex [49, 51]. The proteins from the SYP61 subclade contain the typical structural configuration of t-SNARES, including the N-terminal syntaxin-6 domain, followed by a C-terminal t-SNARE coiled-coil homology domain and a transmembrane segment at the C-terminus. The proteins from the NISP subclade harbor the N-terminal syntaxin-6 domain and the transmembrane segment but lack a C-terminal SNARE domain, which may impair the participation of these new syntaxin-6-like proteins in the assembly of a functional trans-SNARE complex for membrane fusion. Nevertheless, we have demonstrated that a representative of this new syntaxin-6 subclade, NISP, is functionally expressed in plant cells. In addition to being induced by begomovirus infection, NISP functions as a susceptibility factor to begomoviruses, forms complexes with viral and host proteins and is located in TGN/endosome-associated vesicles. We also mapped the NSP-binding domain on NISP to the syntaxin-6 domain. Therefore, we propose that NISP may function as an intracellular vesicle docking site to which the cargo proteins are specifically recruited through interactions with the syntaxin-6-like domain.

We also showed that members of the new syntaxin-6 subclade might diverge functionally concerning protein-protein binding specificity. The NISP paralog, AT218860, which shares with NISP a conserved syntaxin-6 domain (78.6% identity), fails to interact with viral NSP and hence does not affect begomovirus infection. Therefore, specificity of protein interactions may rely on marginal features of the syntaxin-6 domain. A striking difference between NISP and AT2G18860 is an eight-amino acid deletion toward the N-terminus of the syntaxin-6 domain, which may account for the NISP specificity to NSP (S4 Fig). Complementary studies are necessary to precisely define the determinants of a functional syntaxin-6 domain for NSP interaction, which will be useful for engineering resistance to begomoviruses.

### NISP may participate in the intracytoplasmic transport of NSP-vDNA

Based on the current data and others previously reported data, we propose a model for the NISP integration into an intracytoplasmic transport route used for the begomovirus anterograde movement. Begomoviruses replicate in the nuclei of infected cells, where NSP interacts with newly-synthesized vDNA and facilitates the translocation of vDNA to the cytoplasm via nuclear pores [13, 32]). This nuclear exportation process may be mediated by an exportin-like protein yet to be identified. At the cytosolic side of the nuclear envelope, NIG associates with the NSP-vDNA complex to accessorize the release of NSP-vDNA from the nuclear pores into the cytosol [12,18]. The pro-viral function of NIG is associated with its cytosolic localization because the WWP1-mediated recruitment of NIG into nuclear bodies impairs begomovirus infection [48]. At the cytosol, we showed here that NISP may recruit the NIG-NSP-vDNA complex to TGN/endosome-associated transport vesicles. The nature of the NISP-associated vesicle was identified using cell biology molecular markers and FM4-64, which demonstrated that NISP is predominantly localized at TGN and in the early endosome (Fig 3), which was consistent with the localization of syntaxin-6 proteins in most eukaryotic cells [40]. As MP has been demonstrated to associate with the endosomal SYTA, a possible NISP sequestration of NSP-DNA into TGN/endosome-associated vesicles may provide the means for the predicted interactions between endosomally localized MP-SYTA and NSP-DNA, which would favor the MP-mediated intercellular transport of vDNA via an endocytic recycling pathway [35]. In summary, our data support a model in which direct interactions between the host factor NISP and NIG/NSP modulate vDNA intracellular trafficking to endosomes where MP, recruited by the endosomal SYTA, opportunistically interacts with NSP-vDNA to mediate the cell-to-cell movement of vDNA via plasmodesmata.

## Material and Methods

### Phylogenetic analysis

The amino acid full-length sequences of plant proteins were retrieved from TAIR (http://arabidopsis.org/), Phytozome v12 database (https://phytozome.jgi.doe.gov/) and amino acid full-length sequences of other species from the Ensemble database (https://www.ensembl.org/index.html). The syntaxin domain superfamily was predicted using the software HMMER v3.2.1 (http://hmmer.org/) using Pfam v.30 database (https://pfam.xfam.org/). The syntaxin domain-containing protein sequences were selected to perform the phylogenetic reconstruction. The amino acid sequences were aligned using the MUSCLE algorithm [52]. Phylogenetic trees were constructed using Bayesian Inference with MrBayes v3.2.2 [53] and mix model of amino acid substitution (Wag). The analyses were performed running 10.000.000 generations and excluding the first 2.500.000 generations as burn-in and visualized with Figtree software (http://tree.bio.ed.ac.uk/software/figtree/). Plant NISP homologous proteins were identified using the BLAST algorithm implemented in the Ensemble and Phytozome databases. The amino acid sequences of the NISP homologs were selected for phylogenetic reconstruction of N-terminal Syntaxin-6 family using the same tools and criteria applied for the superfamily syntaxin. However, the amino acid substitution model Jones was used for the N-terminal Syntaxin-6 phylogenetic tree.

### PPI network analysis

NSP- and MP-interacting proteins from Arabidopsis were used as a query term to identify their respective interactions described in the BAR database (Bio-Analytic Resource for Plant Biology, http://bar.utoronto.ca/interactions/). The Biogrid and IntAct databases were selected for searching. The protein-protein interactions (PPI) were retrieved and imported into the Cytoscape software (https://cytoscape.org/) to visualize the topology of the PPI network and to calculate the network centrality metrics for each protein. The measured network centralities were betweenness, closeness, eccentricity, and degree. Briefly, the betweenness centrality in the PPI network of the graph G = (V,E) was calculated by observing the number of times a node v (protein) contributes as a link along the shorter paths among all nodes. The betweenness centrality of this node can be analyzed as:

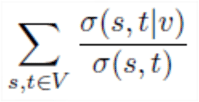

where V is the total set of nodes, *σ*(*s,t*) is the number of shorter paths of (*s,t*), and *σ*(*s,t*|*v*) is the number of paths crossing *v* [54]. The betweenness centrality can indicate the relevance of a protein as functionally capable of keeping together interacting proteins. Closeness centrality (of a node v is the sum of the shortest path distances from w to all other nodes and calculated as described by Freeman [55]:

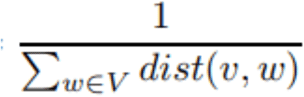

A high closeness value can be determined by the fact that all nodes are usually close to node v. The eccentricity centrality of a node v is the maximum distance from v to all other nodes in graph G and can be analyzed as described by Hage and Harary [56]:

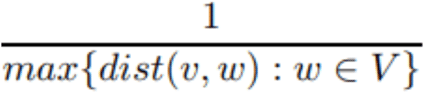

In addition, the degree centrality of node *v* was calculated by ∑ *i*=1*^K^di*, where *d* represents each adjacent node and *K*, the total number of adjacent nodes. Nodes with a high degree can be designated hubs and hold together different nodes with a lower degree.

### GO analysis of the PPI network

The gene set enrichment analysis (GSEA) of the proteins that compose the PPI network was performed using the Bioconductor GOstats package [57]. GSEA analysis computes hypergeometric test p values for over- or underrepresentation for each GO term, and the annotation of genes is based on the Bioconductor annotation package org.At.tair.

### Plasmid constructions

All recombinant plasmids were generated by the GATEWAY cloning system (Invitrogen, http://www.invitrogen.com/). The primers used for gene amplification are listed in S3 Table. The coding regions of *NISP* (AT4G30240) and AT2G18860, with or without a translational stop codon, were amplified from leaf cDNA of *Arabidopsis thaliana* ecotype Columbia (Col-0) and inserted into the entry vectors pDONR201 and pDONR207. The resulting clones were designated At4g30240-St-pDONR201 (pUFV2134), At4g30240-Ns-pDONR201 (pUFV2135), At2g18860-St-pDONR201 (pUFV3193), At2g18860-Ns-pDONR201 (pUFV3194), At4g30240-St-pDONR207 (pUFV2236), At4g30240-Ns-pDONR207 (pUFV2237), At2g18860-St-pDONR207 (pUFV3195), At2g18860-Ns-pDONR207 (pUFV3196). Then, the DNA inserts were transferred by recombination from the entry vector to different yeast and plant expression vectors.

For the yeast two-hybrid assay, the *NISP* and AT2G18860 coding regions were transferred from the respective entry vectors (At4g30240-St-pDONR201 and At2g18860-St-pDONR201) to the yeast expression vectors pDEST32 and pDEST22, generated the clones At4g30240-St-pDEST32 (pUFV2141), At4g30240-St-pDEST22 (pUFV2142), At2g18860-St-pDEST32 (pUFV3197) and At2g18860-St-pDEST22 (pUFV3198), fused to binding domain (BD) (pDEST32) or activation domain (AD) (pDEST22) of GAL4. The *NSP*- and *NIG*-derived clones used in the yeast two-hybrid assay have been previously described and designated NSP-St-pDEST32, NSP-Cl-pDEST22 (Fontes *et al*., 2004), and NIG-pDEST22 (pUFV2547; Calil *et al*., 2018).

For the subcellular localization experiments, the *NISP* and AT2G18860 coding region were fused to GFP and expressed in planta under the control of the CaMV 35S promoter in the vector pK7FWG2. For the co-immunoprecipitation assay, in addition to the GFP fusion, *NISP* ORF was also fused to hemagglutinin (HA) tag, using the binary destination vector pK7m34GWfor three fragment recombination. The resulting DNA constructions were At4g30240-pK7FWG2 (pUFV2576), At2g18860-pK7FWG2 (pUFV3199) and 2×35S::At4g30240-Ns-6HA-pH7m34GW (pUFV2586).

For the subcellular co-localization assays, the ER-associated molecular marker ER-mCherry (CD-959), the TGN-associated molecular marker G-mCherry (CD-967) and the tonoplast-associated molecular marker vac-mCherry (CD-975) were obtained from the Arabidopsis Biological Resource Center. The plasma membrane-associated molecular marker, FLS2-mCherry was obtained by amplifying the *FLS2* ORF from Arabidopsis cDNA with the primers AtFLS2-Fwd and AtFLS2-NS-Rvs (S3 Table), inserting the amplified fragment by recombination into pDON207 and transferring it by triple recombination to the destination vector pH7m34GW along with FLS2-Ns-pDON207, pDONR-P4-P1R+2×35S, and pDONR-P2R-P3+mCherry, resulting in the clone FLS2-mCherry (pUFV3034). Likewise, the SYTA ORF was amplified from leaf cDNA with the primers AtSYTA-Fwd and AtSYTA-NS-Rvs (S3 Table), cloned into the entry vector pDONR-207 and transferred via triple recombination to the destination vector pH7m34GW along with SYTA-NspDON207, pDONR-P4-P1R+2×35S, and pDONR-P2R-P3+mCherry, generating SYTA-mCherry (pUFV3296).

The *NISP* coding region was also fused to GFP and expressed in plants under the control of its endogenous promoter. For this, about 2000 bp of 5’ flanking sequence of At4g30240 was PCR-amplified using the primers described in S3 Table and cloned into pDONR-P4-P1R, yielding proAt4g30240-pDONR-P4-P1R (pUFV2436). Then, the 5’ flanking sequence was transferred by triple recombination to the destination vector pK7Fm34GW along with At4g30240-Ns-pDON201, proAt4g30240-pDONR-P4-P1R, and eGFP-pDONR-P2R-P3, resulting in the clone proAt4g30240::At4g30240-GFP (pUFV2849). The *NSP*- and *NIG*-derived clones used for co-immunoprecipitation assays have been previously described and designated as 2×35S-NSP-Cl-6HA-pH7m34GW (pUFV2424), NSP-Cl-pK7FWG2(pUFV2536), 2×35S-NIG-Ns-6HA-pH7m34GW (pUFV1946) and NIG-pK7FWG2 (pUFV1177) [48].

The SPYNE-GW and SPYCE-GW vectors, which contain the N-terminal (nYFP) and C-terminal (cYFP) of YFP, respectively, were used for the BiFC experiments. At4g30240-Ns-pDONR207 was transferred to SPYNE-GW and SPYCE-GW, yielding At4g30240-Ns-SPYNE (pUFV2238) and At4g30240-Ns-SPYCE (pUFV2239). The *NSP*- and *NIG*-derived clones used in the BiFC assays have been previously described and designated as NSP-Cl-Ns-SPYNE (pUFV1654), NSP-Cl-Ns-SPYCE (pUFV1655), NIG-Ns-SPYNE (pUFV1646), and NIG-Ns-SPYCE (pUFV1647) [48].

NISP truncated versions were obtained from At4g30240-St-pDONR201 by PCR using primers listed in S3 Table. The generated fragments were cloned into the entry vector pDONR201 and designated as At4g30240(1-107)-St-pDONR201 (pUFV2806), At4g30240(105-200)-St-pDONR201 (pUFV2807) and At4g30240(200-300)-St-pDONR201 (pUFV2808). The respective DNA inserts were then transferred to the yeast expression vectors, yielding At4g30240(1-107)-St-pDEST32 (pUFV2822), At4g30240(1-107)-St-pDEST22 (pUFV2819), At4g30240(105-200)-St-pDEST32 (pUFV2823), At4g30240(105-200)-St-pDEST22 (pUFV2820), At4g30240(200-200)-St-pDEST32 (pUFV2824) and At4g30240(200-300)-St-pDEST22 (pUFV2821).

### Detecting MP- and NSP-host protein interactions on high-density halo tag-NAPPAs

The identification of NSP and MP partners was carried out through a previously fabricated high-density HaloTag nucleic acid programmable protein array (HaloTag-NAPPA), as described by Yazaki *et al*. [37]. The protein microarray, which consisted of a set of *in situ* synthesized proteins from 4600 *Arabidopsis thaliana* ORFs, has been designated AtNAPPA01. 3×HA-fused CabLCV MP and NSP proteins were independently expressed using the TNT system (Promega) according to the manufacturer’s recommendations. The *in vitro* synthesized HA-fused MP, and HA-fused NSP were used to probe individually AtNAPPA01. The identification of MP- and NSP-interacting proteins with the NAPPA protein-protein interaction method was from a single experiment using duplicated spots on one glass slide. Interactions were considered positive only when both replicated spots displayed normalized signal intensity above background. Experiments, data scanning, quantification, and data processing were carried out as described by Yazaki *et al*. [37].

### Plant growth conditions and transformation

*Arabidopsis thaliana*, ecotype Columbia (Col-0), was used as the wild type. Homozygous seeds from the T-DNA insertional mutants *nisp-1* (Salk_117751) and *at2g18860-1* (Salk_099699) were obtained from the Arabidopsis Biological Resource Center. Arabidopsis seeds were surface-sterilized and cold treated at 4°C for 72 h in the dark and then exposed to white light. Seedlings were grown at 22°C on plates containing Murashige-Skoog (MS) medium for 2 weeks and then transferred to soil. Plants were grown in a growth chamber at 22 °C under short-day conditions (12h light/12h dark). Transgenic lines were obtained via *A. tumefaciens-mediated* transformation using the floral dip method [58]. Transformed plants with the indicated DNA constructions were selected in MS medium supplemented with the appropriate antibiotic (kanamycin, pK7FWG2).

Selected transformants were confirmed by PCR, transferred to soil, and grown in a growth chamber at 22 °C under long-day conditions (16h light/8h dark) to generate seeds. For segregation analysis, seeds were germinated on MS medium containing 50 μgmL^-1^ kanamycin. Homozygous T1 lines for the T-DNA loci were selected by determining the frequency of their kanamycin-resistant T2 seeds after self-pollination. Accumulation of NISP-GFP and AT2g18860-GFP transcript and protein levels were monitored in each generation by qRT-PCR using gene-specific primers (S3 Table) and immunoblotting using anti-GFP antibody. From the independently transformed lines (8-15 lines of each construct and genotype transformed), we selected three independently transformed lines which appeared to have an integrated T-DNA locus on a single chromosome, since 75% of their T1 segregating seedlings were resistant to kanamycin (S4 Table). The selected lines are presented in S4 Table. The nomenclature *nisp*-1/NISP-1-3 indicates that the genotype *nisp-1* was transformed with At4g30240-pK7FWG2, harboring the *NISP* ORF fused to the N-terminus of GFP under the control of the 35S promoter to generate NISP-complementing lines. The designations AtNISP-1, AtNISP-2 and AtNISP-3 indicate three independently transformed lines, which were generated by transformation of Col-0 with At4g30240-pK7FWG2 to obtain NISP-GFP-overexpressing lines. The same nomenclature was applied for the transformation of *at2g18860-1* mutant and Col-0 with At2g18860-pK7FWG2, containing the At2g18860 ORF fused to the N-terminus of GFP under the control of 35S promoter (S4 Table).

### RNA isolation and qRT-PCR analysis

Primers were designed using the Primer Express 3.0 (Life Technologies). Total RNA was extracted from Arabidopsis leaves using TRIzol reagent (Thermo Fisher Scientific) and quantified with a NanoDrop spectrophotometer. Total RNA (2 μg) was treated with RNase-free DNase I and reverse transcribed using oligo (dT) (0,5 *μ*M) and 1U of M-MLV (Thermo). Real-time RT-PCR was carried out using SYBR Green PCR Master Mix (Life Technologies), DNA template, and primers in 10 μL of reaction and an ABI7500 real-time PCR system (Life Technologies). The expression of each gene was normalized to the expression of *Actin* and quantified using the 2^-ΔCt^ method. The primers used to detect specific transcripts with real-time RT-PCR are listed in S3 Table.

### Yeast two-hybrid assay

Competent cells of *Saccharomyces cerevisiae*, strain AH109 (MATa, trp1-901, leu2-3, 112, ura3-52, his3-200, gal4Δ, gal80Δ, LYS2::GAL1UAS-GAL1TATA-HIS3, GAL2UAS-GAL2TATA-ADE2, URA3::MEL1UAS-MEL1 TATA-lacZ), were cotransformed with the yeast expression vectors pDEST22 (Gal4 AD) and pDEST32 (Gal4 BD), carrying the coding region of the tested proteins, along with 100 μg of salmon sperm carrier DNA (ssDNA), using the lithium acetate/polyethylene glycol (PEG) method. For the selection of double transformants, the cells were plated on medium deficient in leucine and tryptophan (SD, Synthetic Dropout, -Leu, -Trp) and cultured for 4 days at 28°C. Interactions were monitored by the ability of the reporter strain to grow on media lacking leucine, tryptophan, and histidine but supplemented with 2.5 mM 3-AT (3-Amino-1,2,4-triazole; Sigma) for 4 days at 28°C.

### Co-immunoprecipitation assay

The *in vivo* interactions between NISP and NSP or NIG were monitored by co-immunoprecipitation using the μMACS™ Epitope Tag Protein Isolation (MACS/Miltenyil Biotec) kit, according to manufacturer’s recommendations. *Agrobacterium tumefaciens* (*strain GV3101*)-mediated transient expression in *N. benthamiana* leaves was carried out as described (12). Total protein extracts were obtained from infiltrated leaves expressing the recombinant proteins. At 48 h after agro-infiltration, 200 mg of leaves were homogenized with 1 ml of lysis buffer (50 mM Tris-HCl pH 8.0, 1 mM PMSF, 2mM benzamidine, 1% (v/v) Nonidet P-40) and incubated for 2 h with anti-green fluorescent protein (GFP) magnetic beads (MACS / Miltenyi Biotec) at 4 °C under rotation. After incubation, the extracts were applied to a MACS column for the indicated protein tag, washed five times before elution with a pre-heated buffer at 95 °C. The immunoprecipitated proteins were separated by sodium dodecyl sulfate-polyacrylamide gel electrophoresis (SDS-PAGE) 10% (p/v), immunoblotted with the monoclonal antibodies anti-human influenza hemagglutinin (anti-HA) epitope tag, amino acids 98-106 (Miltenyi Biotec, 130-091-972; 1:10.000) or anti-GFP (Miltenyi Biotec, 130-091-833; 1:10.000), and detected using Signal West Pico Chemiluminescent Substrate (Thermo Scientific) according to the recommendations of the manufacturer.

### Bimolecular fluorescence complementation (BiFC) assay

Different combinations of *A. tumefaciens* strain GV3101 expressing the indicated proteins fused to N-YFP/C-YFP were co-infiltrated into the abaxial surface of *N. benthamiana* leaves at OD_600nm_ of 1:1 proportion. Three days after infiltration, fluorescence was examined in epidermal cells using a Zeiss inverted 510 META laser scanning microscope equipped with an argon laser and a helium laser as excitation sources. YFP was excited at 514 nm using the argon laser, and YFP emission was detected using a 560–615-nm filter. The images were processed using LSM Image Browser 4 (ZEISS) software.

### Subcellular localization

The subcellular localization of proteins was examined in epidermal cells of *N. benthamiana* leaves transiently expressing GFP fusions. Briefly, *N. benthamiana* leaves were agro-infiltrated with *A. tumefaciens* strain GV3101 carrying proAt4g30240-pDONR-P4-P1R (pUFV2436), At4g30240-pK7FWG2 (pUFV2576) or At2g18860-pK7FWG2 (pUFV3199) (Method S1) and fluorescence of the GFP fusions were visualized in epidermal cells 3 days after infiltration by confocal microscopy using a Zeiss inverted 510 META laser scanning microscope equipped with an argon laser and a helium laser as excitation sources. GFP was excited by the argon/helium-neon laser system in 488 nm wavelength, and emission was collected in 500–530 nm band-pass filter. Images were captured and processed with LSM Image Browser 4 (Carl-Zeiss) software.

For co-localization assays, NISP-GFP was co-expressed with ER-, Golgi-, tonoplast-, plasma membrane- and endosome-associated molecular markers fused to mCherry (see in plasmid constructions) and fluorescence was examined by confocal microscopy, using a Zeiss inverted LSM510 META laser scanning microscope, on whose system, the default scanning mode is sequential scanning (multitrack mode). For imaging GFP, the 488 nm excitation line and the 500 to 530 nm band pass filter were used. Excitation of mCherry was at 540 nm and emission of 608-680 nm. For the FM4-64 staining, 48 h after agro-infiltration with the DNA construct expressing NISP-GFP, the leaves were infiltrated with a solution of 50 μM FM4-64 (Molecular Probes, Eugene, OR) and examined by confocal microscopy 1h post-infiltration.

### Infectivity assay

The generation of transgenic lines overexpressing NISP (AtNISp) and At2g18860 (AT2G18860), knockout lines (*nisp-1* and *at2g18860-1)*, NISP complementing lines are described in Method S1. Transgenic lines and Columbia at the seven-leaf stage were inoculated with plasmids containing partial tandem repeats of CabLCV DNA-A and DNA-B by biolistic delivery using a microprojectile bombardment model PDS-1000/He accelerator at 900 psi. (24, 26, 59). The inoculated plants were transferred to a growth chamber and monitored for symptoms development and progress of infection [23, 60]. Total DNA was extracted from systemically infected leaves [61], and viral DNA accumulation was detected by PCR using universal DNA-B specific primers PBL1v2040 and PCRC1 [62] (S3 Table). In each experiment, 15 plants of each line were inoculated with 2 μg of tandemly repeated DNA-A plus DNA-B per plant. The course of infection was examined using data from three independent experiments.

### Quantification of viral DNA in infected plants

The accumulation of viral DNA in infected plants was quantified by qPCR [26]. The reactions were prepared in 10 μL using SYBR Green Master Mix (Life Technologies), 100 ng of total DNA from systemically infected leaves, and CabLCV-DNA-B-specific primers (S3 Table), and analyzed on a real-time PCR system. The genomic copies of CabLCV were normalized against an internal control (18S rRNA) to account for template input variation between tubes. For viral DNA quantification, standard curves were prepared using serial dilutions of DNA-B of CabLCV (100 to 10^6^ copies of viral genome per reaction).

### Formaldehyde fixation and immunoprecipitation of NISP-GFP:vDNA complexes

A modified ChIP assay evaluated the in vivo interaction between NISP and viral DNA. NISP-overexpressing transgenic lines (AtNIS) and Columbia (Col-0) at the six-leaf stage were inoculated by biolistic delivery with CabLCV DNA-A and DNA-B infectious clones. Tissue harvest, formaldehyde fixation, and chromatin breaks from 45 days-old infected plants were conducted according to Alves et al. [63] and Yamaguchi et al. [64]. For cross-linking proteins to DNA with formaldehyde, 600 mg of infected leaves were washed with 10 mL of PBS (10 mM Na2HPO4, 1.8 mm KH2PO4, 137 mM NaCl, 2.7 mM KCl, pH 7.4) and fixed in 10 mL PBS by adding formaldehyde to 1% final concentration. The infiltration of formaldehyde into vegetable cells was carried out under vacuum at −500 Pa for 1.5 min, then 1 min twice with 30 s intervals. After washing first with 0.125 M glycine under vacuum for 1.5 min and then with PBS, the tissue was air-dried. The fixed samples were homogenized in liquid N2 and 1 mL of extraction buffer (50 mM Tris-HCl pH 8.0, 10 mM EDTA, 1% (v/v) Nonidet-P40). After incubation for 30 min on ice, cell debris was removed by centrifugation at 12000 x g for 10 min. The supernatant (300 μL) was diluted with 700 μL of the ChIP dilution buffer (16.7 mM Tris-HCl pH 8,0, 167 mM NaCl, 1.2 mM EDTA, 0.01% SDS) and DNA was fragmented by sonication at an 80 μm amplitude level, six pulses for 30 min, 30 s on/off on the ice. Then, 700 μL of the sonicated suspension was mixed with 200 μL of the ChIP dilution buffer, supplemented with 1.1% (v/v) Triton X-100 and centrifuged at 13200 x g for 5 min at 4° C. The supernatant was incubated with 50 μL of protein-A-garose beads for 2 h. After centrifugation at 900 x g for 3 min, 18 μL of the supernatant were stored at −80° C, as input control. The remaining supernatant was incubated with 1 μl of polyclonal anti-GFP serum (Life Technologies A11122; 3x diluted) for 4h at 4° C. Anti-GFP-NISP-GFP:vDNA complexes were captured by incubation with 50 μL of potein-A-garose beads for 4 h at 4°C, under gentle agitation. After centrifugation at 900 x g for 3min, the pellet was washed five times with different washing buffers in the order: twice with the washing buffer A (0.1% (w/v) SDS 1% (v/v) Triton X-100, 2 mM EDTA, 20 mM Tris-HCl, pH 8,1, 150 mM NaCl), once with the buffer B (0.1% (w/v) SDS 1% (v/v) Triton X-100, 2 mM EDTA, 20 mM Tris-HCl, pH 8,1, 500 mM NaCl), followed by the buffer C (250 mM LiCl, 1% (v/v) IGEPAL-CA630, 1% (w/v) sodium deoxycholate, 1 mM EDTA, 10 mM Tris-HCl pH 8,1) and finally with 0,5 X TE (10 mM Tris-HCl pH 8,0, 1 M EDTA). The DNA pellet was reverse cross-linked from proteins and purified using the QIAprep^®^ Spin Miniprep kit.

### ChIP-qPCR

NSP primers (S3 Table) were initially used to amplify viral DNA-B from the immunoprecipitated DNA and input DNA to ensure DNA-B-specific amplification in samples. Likewise, DNA-A-specific primers were used to examine initial amplification of the DNA-A component. Each reaction contained 1 μL of immunoprecipitated DNA or 20-fold diluted input DNA, 0.2 mM CabLCV-DNA-B-specific primers (S3 Table), and 5 μL of SYBR Green PCR Master Mix (Life Technologies) in 10 μL total volume. The thermal cycling conditions of the reactions consisted of an initial step at 95°C for 10 min, followed by 40 cycles at 94°C for 15 s and 58°C for 45 s. The cycle threshold (Ct) values from Col-0 and AtNISP were normalized with the Ct values of the respective inputs, and the results were expressed as a percentage of the input.

### Author Contribution

B.C.G.M performed most of the experiments; L.G.C.M. performed the co-localization experiments and protein localization assays; M.D.-B., A.Y.K. and J.Y. performed protein microarray assay; B.C.G.M. and J.P.B.M. performed two hybrid assays; J.C.F.S. performed *in silico* and statistical analyses; B.C.G.M. and A.A.S. performed BiFC assays; E.P.B.F. and J.R.E. conceived experiments; B.C.G.M. and E.P.B.F. designed all experiments, analyzed the data and wrote the manuscript.

## Supporting information

S1 Table- Protein-protein interactions between the viral proteins and the Arabidopsis proteins.

S2 Table- Enriched GO terms in three categories, Biological Process, Molecular Function, or Cellular Component ontology

S1 Video/Movie -Vesicle-associated localization of NISP

## Acknowledgments

The authors are grateful to the microscopy facility at Universidade Federal de Viçosa.

## Supporting information

**S1 Table-** Protein-protein interactions between the viral proteins and the Arabidopsis proteins.

**S2 Table-** Enriched GO terms in three categories, Biological Process, Molecular Function, or Cellular Component ontology

**S3 Table.** List of primers used for cloning, PCR, RT-qPCR and qPCR

**S4 Table.** Expression of kanamycin resistance in the T1 generation of transgenic Arabidopsis plants.

**S1 Video/Movie** -Vesicle-associated localization of NISP

**S1 Fig. Identification of MP- and NSP-interacting proteins.**

*In vitro* synthesized 3xHA-NSP and 3xHA-MP. In vitro translated HA-NSP and HA-MP were electrophoresed by SDS-PAGE and immunoblotted using an anti-HA antibody. **(B)** Protein interaction between *in situ* synthesized Arabidopsis hallo-proteins and 3xHA-MP or 3xHA-NSP. The AtNAPPA02 high-density array containing 4600 Arabidopsis ORFS spotted in duplicates was probed with 3xHA-MP or 3xHA-NSP, and candidate interactors were detected with an anti-HA antibody. The bright spots are signals from candidate interactors.

**S2 Fig. A network of NSP- and MP-directly interacting Arabidopsis proteins.**

The network was assembled by the Cytoscape software. The viral proteins are indicated in red, MP-specifically interacting proteins in yellow, NSP-specifically interacting proteins in orange and proteins that associate with both MP and NSP are indicated in green.

**S3 Fig. *In silico* analyses of NISP.**

**(A)** Phylogenetic tree of syntaxin superfamily domain-containing proteins from the species indicated. The phylogenetic tree was constructed using Bayesian inference performed with MrBayes v3.2.2 and the mixed amino acid substitution model Wag. The syntaxin-6 domain-containing proteins from Arabidopsis are indicated in blue, and NISP (AT4G30240) was clustered with plant-specific syntaxin-6 proteins forming the red clade. **(B)** Phylogenetic tree of N-terminal Syntaxin-6 domain-containing proteins. NISP (AT4G30240) clusters together with plant-specific N-terminal syntaxin-6 proteins, which do not harbor a typical t-SNARE domain at the C-terminus. The syntaxin-6 domain proteins from Arabidopsis are in blue. The phylogenetic tree was constructed using Bayesian inference performed with MrBayes v3.2.2 and the mixed amino acid substitution model Jones.

**S4 Fig. Sequence alignment of NISP and AT2G18860**

The N-terminal syntaxin-6 domain is indicated in red and the transmembrane segment in black. The sequence alignment was performed using CLUSTAL OMEGA. Identical amino acids are indicated with asterisks, highly conserved residues with (:) and lower conservation with (.).

**S5 Fig. Subcellular localization of AT2G18860**

AT2G18860-GFP distribution in plasma membrane-associated vesicles. *N. benthamiana* leaves were infiltrated with *A. tumefaciens* carrying a DNA construct expressing AT2G18860-GFP under the control of 35S promoter. Confocal images were taken 36 h post-infiltration. Approximately 200 cells were examined. Scale bars, 10 μm.

**S6 Fig**. ***NISP*, but not AT2G18860, expression is induced by CabLCV**

NISP and AT2G18860 transcript levels were quantified by qRT-PCR in uninfected Col-0 leaves (Col-0), tungsten-inoculated Col-0 leaves (Col-0 T), and systemically infected Col-0 leaves with CabLCV (Col-0 IN). Gene expression was calculated using the 2^-ΔCt^ method, and actin was used as an endogenous control. Error bars indicate 95% confidence intervals based on replicated samples (n = 3) from three independent experiments.

**S7 Fig. CabLCV infection-associated symptoms in Arabidopsis genotypes at 21 days post-inoculation (dpi).** The figure shows representative samples of mock-inoculated and CabLCV - infected plants. The genotypes are indicated in the figure.

**S8 Fig. The transgenic lines are visibly indistinguishable from the wild-type plants.**

Columbia (Col-0) ecotype of *Arabidopsis thaliana* was used as the wild-type control for phenotype comparison, and the Col-0 ecotype was used for generating almost all the transgenic plants except for the complementation test in *nik1-1* and *at2g18860-1* mutants. The genotypes are the same as in Fig. 5. Plants were grown in a growth chamber at 22°C under long-day conditions (16 h light/8 h dark). **(A)** Developmental phenotypes associated with inactivation of NISP gene and overexpression of NISP-GFP in the R3 generation of transgenic lines at 45 days after germination. **(B)** Developmental phenotypes associated with inactivation of NISP gene and overexpression of NISP-GFP in the R3 generation of transgenic lines at 60 days after germination. **(C)** Developmental phenotypes associated with inactivation of AT2G18860 gene and overexpression of AT2G18860-GFP in the R2 generation of transgenic lines at 45 days after germination. **(D)** Developmental phenotypes associated with inactivation of AT2G18860 gene and overexpression of AT2G18860-GFP in the R2 generation of transgenic lines at 60 days after germination

**S9 Fig. NSP enhances NISP-NIG complex formation.**

**(A)** NISP interacts with NIG in yeast. AD-NIG and BD-NISP fusions were expressed in yeast and interactions between the recombinant proteins were monitored by His prototrophy in selective medium (-Leu,-Trp,-His) and supplemented with 2.5 mM 3-AT. The co-expression of PRNIG fused to BD and CSN5A fused to AD was used as a positive control. **(B)** Cytosolic and perinuclear distribution of NIG-GFP. *N. benthamiana* leaves were infiltrated with A. tumefaciens carrying NIG-GFP constructs with expression driven by the 35S constitutive promoter. NIG-GFP was imaged by confocal microscopy 36 h after infiltration. **(C)** Cytosolic localization of NIG in transgenic lines. Confocal fluorescence image of Arabidopsis root cells stably transformed with 35S:NIG-GFP. **(D)** NISP interacts with NIG *in planta*. Total protein extracts from *N. benthamiana* expressing NISP-6HA and NIG-GFP were used for co-immunoprecipitation assays using anti-GFP. Input and IP show the levels of the expressed proteins NISP-6HA and NIG-GFP. Anti-HA was used to detect NISP-6HA from the immunoprecipitated complex. GFP was used as an unrelated protein. The experiment was repeated three times. **(E)** NISP-NIG complex formation in the presence of viral NSP. The Co-IP assay was performed as described in A, except that co-expressed NSP-GST was included in the assay. NSP-GST was detected by immunoprecipitating it from transfected leaves and immunoblotting with anti-GST. The experiment was repeated three times. **(F)** The interaction of NISP and NIG is increased by the presence of viral NSP. NIG-GFP levels in the immunoprecipitated complex in the presence and absence of viral NSP were quantified using the Band Analysis tools of the ImageLab software (Bio-Rad). The signal values were normalized using the IP NIG-GFP band, revealed in the same blot as the Co-IP band. A.U. denotes arbitrary unit.

**S10 Fig. NISP interacts with NIG *in vivo* and NSP enhances NISP-NIG complex formation.**

**(A)** BIFC assay showing the interaction between NISP and NIG in vesicles of *N. benthamiana* leaf cells. *N. benthamiana* leaves transiently expressing NISP and NIG fused to the C-terminus (cYFP) or N-terminus (nYFP) of YFP, were examined by confocal microscopy 3 days after infiltration. Scale bars, 10 μm and 20 μm. **(B)** Co-expression of NSP-6HA strengthens interaction between NIG and NISP. The BiFC assay was performed by agro-infiltration of *N. benthamiana* leaves with nYFP-NIG, cYFP-NISP and vice-versus along with the expression of NSP-6HA. The reconstituted fluorescence signal was observed by confocal microscopy, 3 days after agroinfiltration. The figure displays representative samples from three independent biological repeats. Scale bars, 10 μm. **(C)** Confocal fluorescent image of NISP and NIG fused to the C-terminus (cYFP) or N-terminus (nYFP) of YFP in combination with empty vectors, as indicated in the figure.

**S11 Fig. Schematic representation of CabLCV DNA-A, DNA-B and ChIP-DNA primers.**

**(A)** Schematic representation of DNA-A. The number 1 corresponds to the 5’ end of the nick site (TAATATT/AC) within the conserved nonanucleotide sequence located in the intergenic region. The ORFs are indicated with arrows. The positions of the primers used for PCR (F1/R1), and qRT-PCR (F2/R2) are indicated. **(B)** Schematic representation of DNA-B. The numbering scheme is the same as in a. The ORFs are indicated with arrows. The positions of the primers used for PCR (F1/R1), and qRT-PCR (F2/R2 and F3/R3) are indicated.

