## Supporting Information, S1-S11 Figures, S3 Table and S4 Table for "A plant-specific syntaxin-6 protein contributes to the intracytoplasmic route for begomoviruses"

**Molecular Plant Supporting Information**

Article title: **A new plant-specific syntaxin-6 protein contributes to the intracytoplasmic route for begomoviruses**

**This file contains:**

Supplementary Figures

S1Fig

S2 Fig

S3 Fig

S4 Fig

S5 Fig

S6 Fig

S7 Fig

S8 Fig

S9 Fig

S10 Fig

S11 Fig

Supplementary Tables

S3 Table

S4 Table

**Additional supplementary files provided separately**

S1 Table

S2 Table

S1 Video/Movie -Vesicle-associated localization of NISP

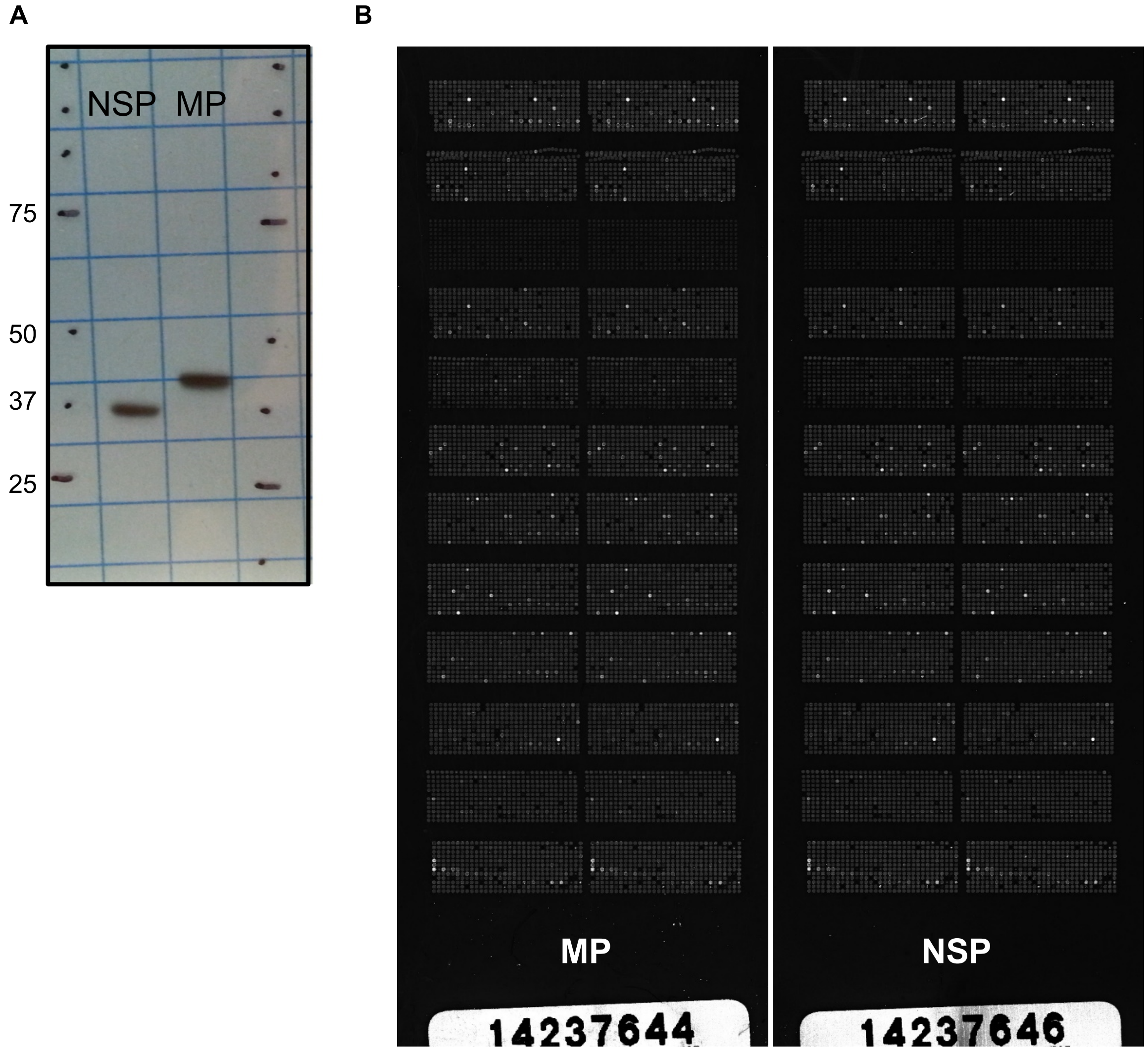


**S1 Fig. Identification of MP- and NSP-interacting proteins.**

**(A)** *In vitro* synthesized 3xHA-NSP and 3xHA-MP. In vitro translated HA-NSP and HA-MP were electrophoresed by SDS-PAGE and immunoblotted using an anti-HA antibody. **(B)** Protein interaction between *in situ* synthesized Arabidopsis hallo-proteins and 3xHA-MP or 3xHA-NSP. The AtNAPPA02 high-density array containing 4600 Arabidopsis ORFS spotted in duplicates was probed with 3xHA-MP or 3xHA-NSP, and candidate interactors were detected with an anti-HA antibody. The bright spots are signals from candidate interactors.

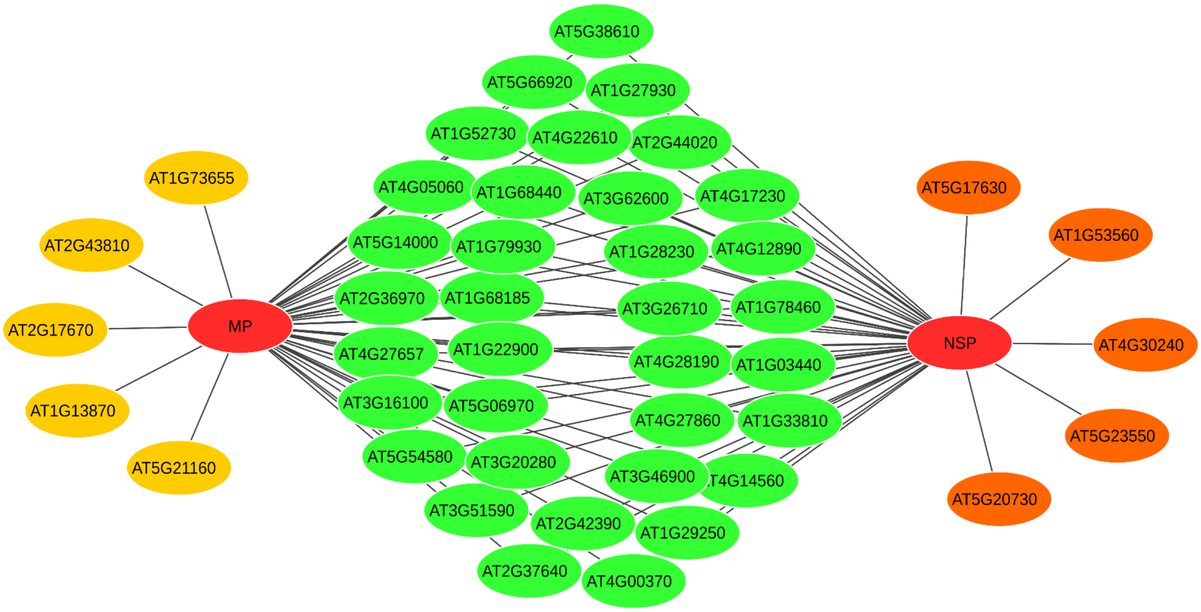


**S2 Fig. A network of NSP- and MP-directly interacting Arabidopsis proteins.**

The network was assembled by the Cytoscape software. The viral proteins are indicated in red, MP-specifically interacting proteins in yellow, NSP-specifically interacting proteins in orange and proteins that associate with both MP and NSP are indicated in green.

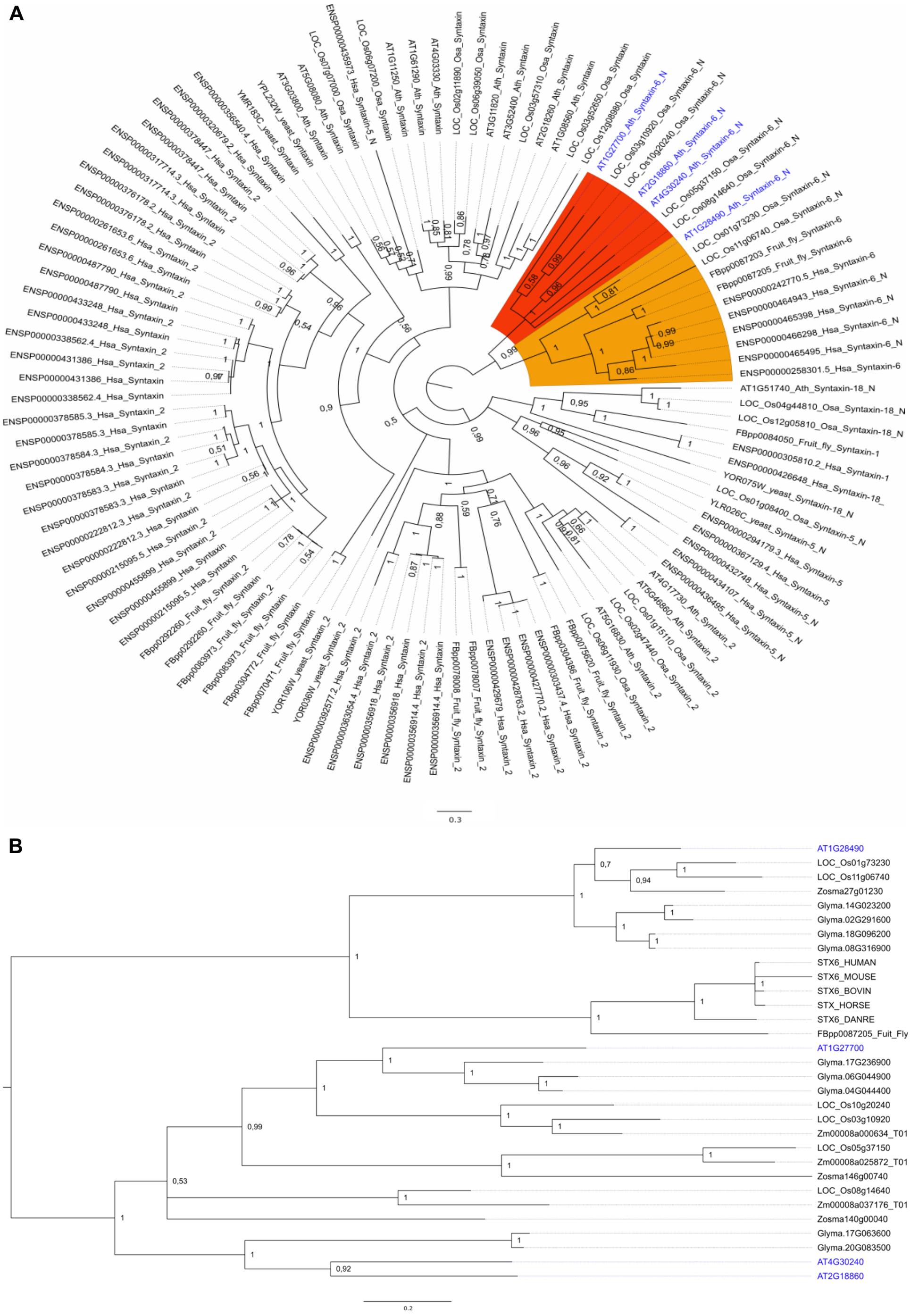


**S3 Fig.** ***In silico*** **analyses of NISP.**

**(A)** Phylogenetic tree of syntaxin superfamily domain-containing proteins from the species indicated. The phylogenetic tree was constructed using Bayesian inference performed with MrBayes v3.2.2 and the mixed amino acid substitution model Wag. The syntaxin-6 domain-containing proteins from Arabidopsis are indicated in blue, and NISP (AT4G30240) was clustered with plant-specific syntaxin-6 proteins forming the red clade. **(B)** Phylogenetic tree of N-terminal Syntaxin-6 domain-containing proteins. NISP (AT4G30240) clusters together with plant-specific N-terminal syntaxin-6 proteins, which do not harbor a typical tSnare domain at the C-terminus. The syntaxin-6 domain proteins from Arabidopsis are in blue. The phylogenetic tree was constructed using Bayesian inference performed with MrBayes v3.2.2 and the mixed amino acid substitution model Jones.

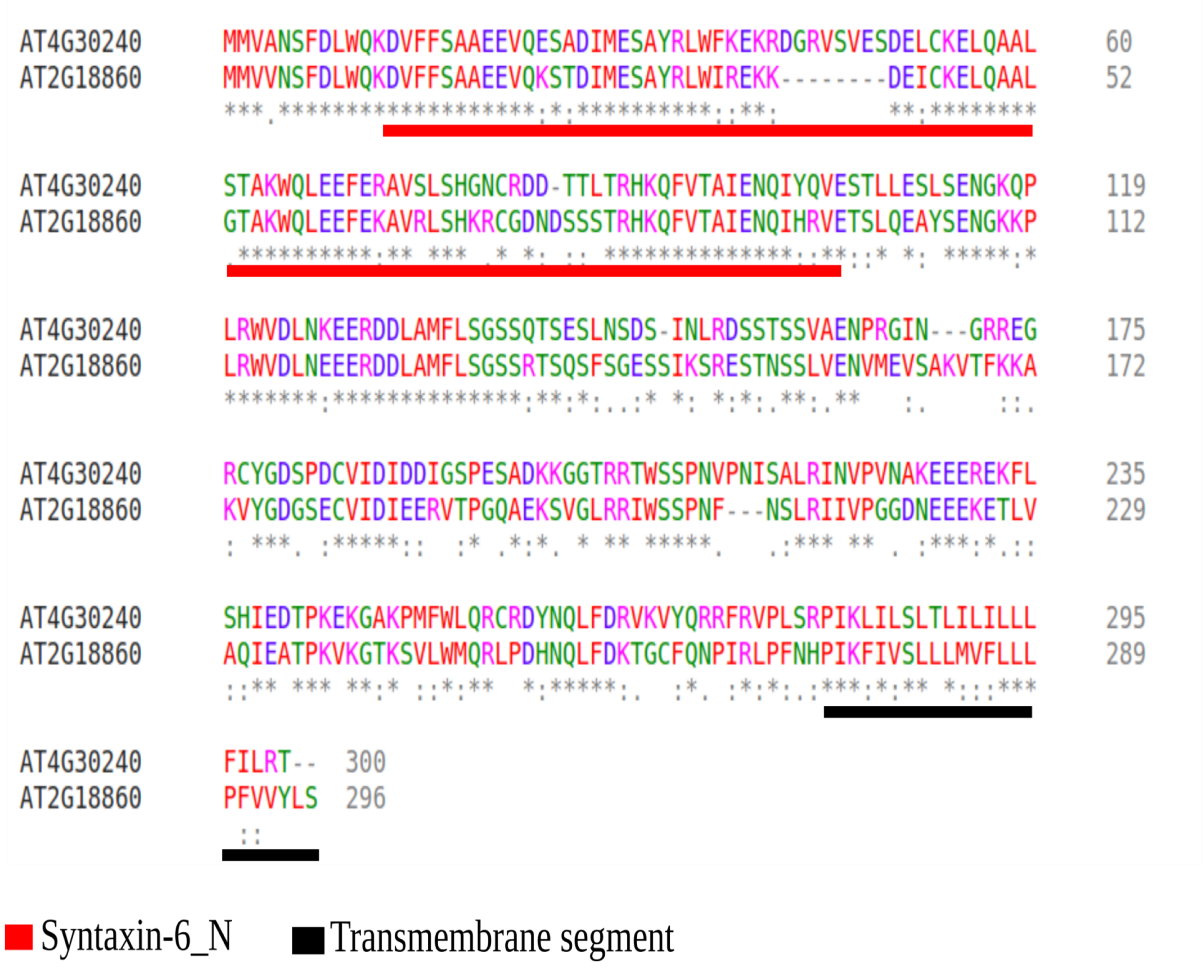


**S4 Fig. Sequence alignment of NSP and AT2G18860**

The N-terminal syntaxin-6 domain is indicated in red and the transmembrane segment in black. The sequence alignment was performed using CLUSTAL OMEGA. Identical amino acids are indicated with asterisks, highly conserved residues with (:) and lower conservation with (.).

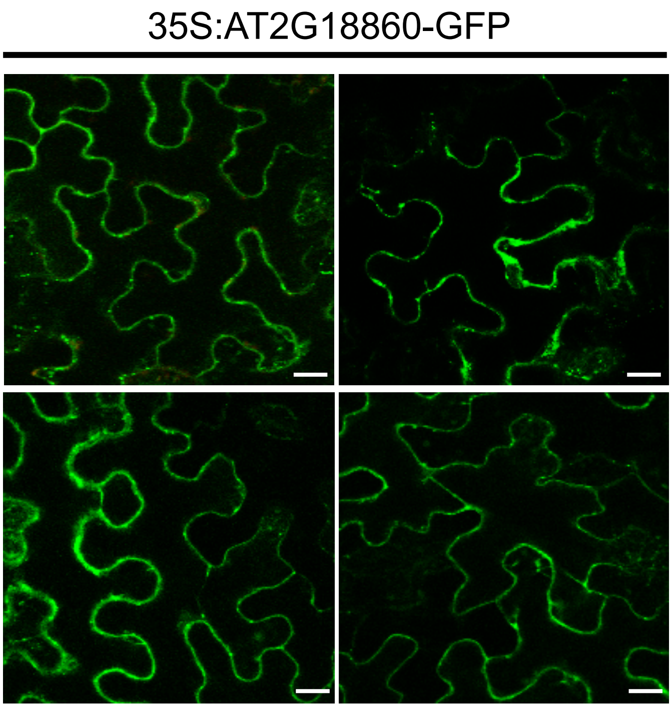


**S5 Fig. Subcellular localization of AT2G18860**

AT2G18860-GFP distribution in plasma membrane-associated vesicles. *N. benthamiana* leaves were infiltrated with *A. tumefaciens* carrying a DNA construct expressing AT2G18860-GFP under the control of 35S promoter. Confocal images were taken 36 h post-infiltration. Approximately 200 cells were examined. Scale bars, 10 µm.

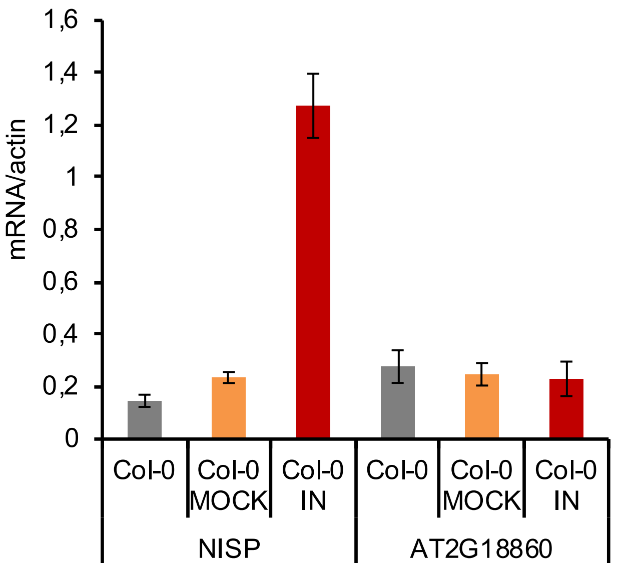


**S6 Fig**. ***NISP*, but not AT2G18860, expression is induced by CabLCV**

NISP and AT2G18860 transcript levels were quantified by qRT-PCR in uninfected Col-0 leaves (Col-0), tungsten-inoculated Col-0 leaves (Col-0 T), and systemically infected Col-0 leaves with CabLCV (Col-0 IN). Gene expression was calculated using the $2^{-\Delta Ct}$ method, and actin was used as an endogenous control. Error bars indicate 95% confidence intervals based on replicated samples (n = 3) from three independent experiments.

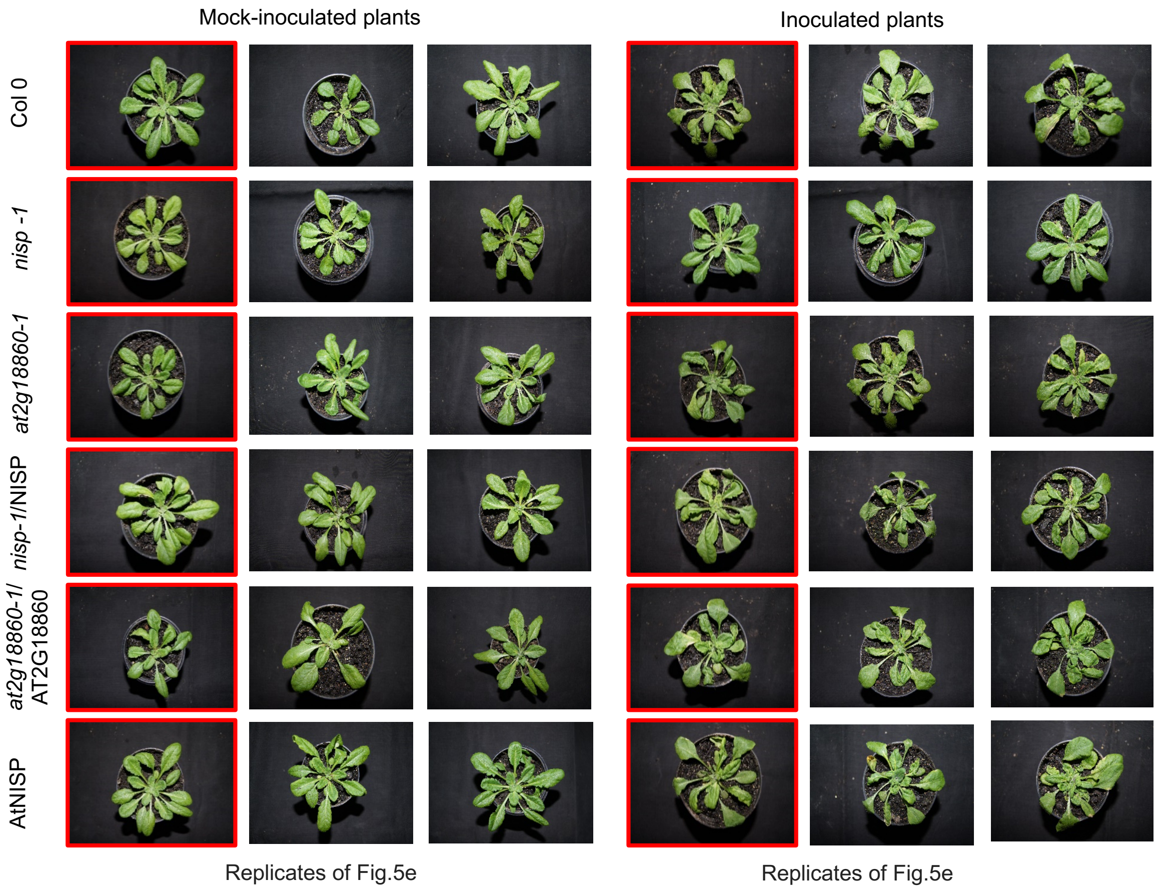


**S7 Fig. CabLCV infection-associated symptoms in Arabidopsis genotypes at 21 days post-inoculation (dpi).** The figure shows representative samples of mock-inoculated and CabLCV -infected plants. The genotypes are indicated in the figure; *nisp-1*/NISP and *at2g18860-1*/AT2G18860 are complemented lines and AtNISP is a NISP-overexpressing line, as indicated in Figure 5A-5D.

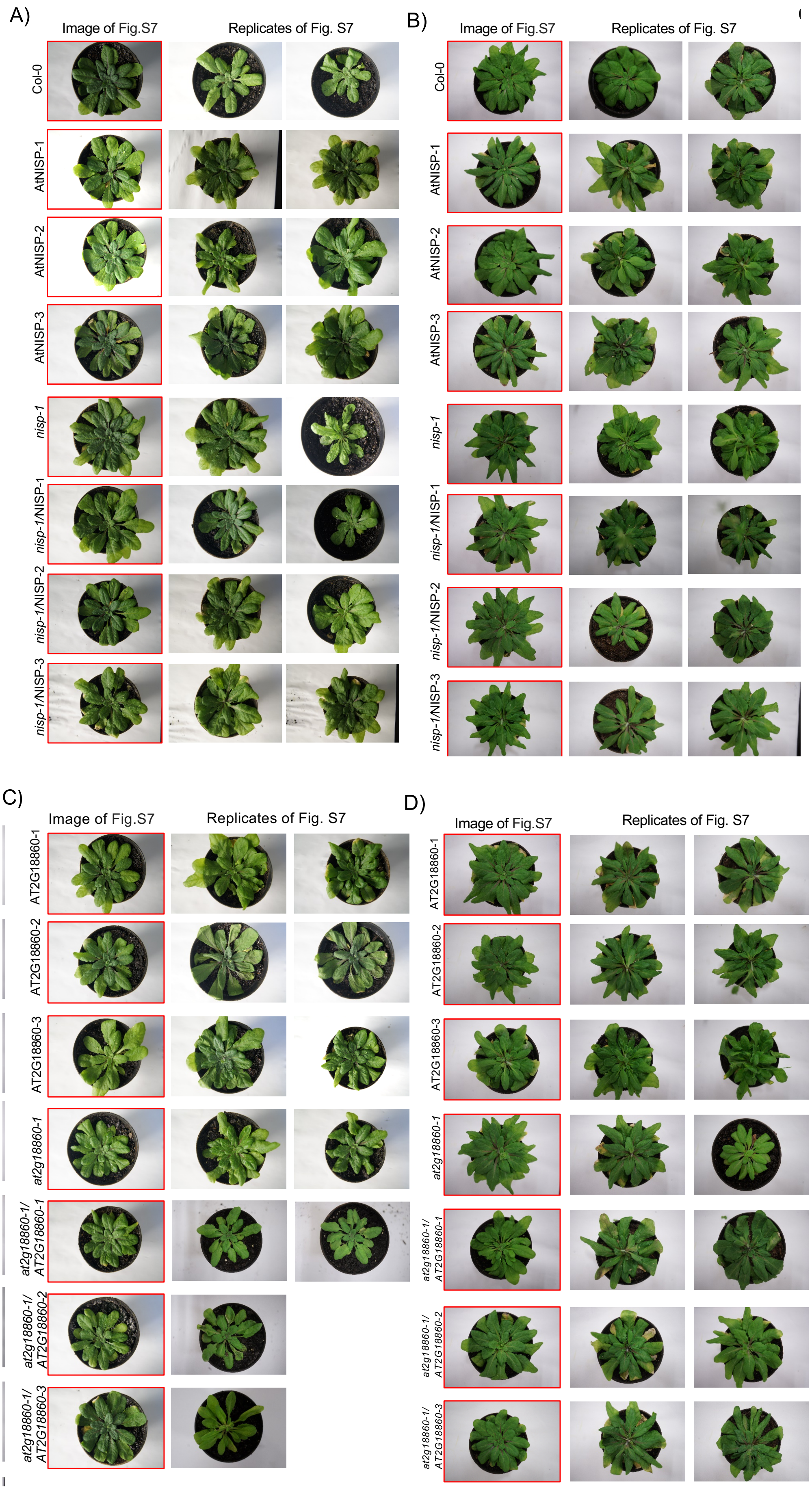
A)
Image of Fig.S7
Replicates of Fig. S7
Col-0
AtNISP-1
AtNISP-2
AtNISP-3
nisp-1
nisp-1/NISP-1
nisp-1/NISP-2
nisp-1/NISP-3
B)
Image of Fig.S7
Replicates of Fig. S7
Col-0
AtNISP-1
AtNISP-2
AtNISP-3
nisp-1
nisp-1/NISP-1
nisp-1/NISP-2
nisp-1/NISP-3
C)
Image of Fig.S7
Replicates of Fig. S7
AT2G18860-1
AT2G18860-2
AT2G18860-3
at2g18860-1
at2g18860-1/ AT2G18860-1
at2g18860-1/ AT2G18860-2
at2g18860-1/ AT2G18860-3
D)
Image of Fig.S7
Replicates of Fig. S7
AT2G18860-1
AT2G18860-2
AT2G18860-3
at2g18860-1
at2g18860-1/ AT2G18860-1
at2g18860-1/ AT2G18860-2
at2g18860-1/ AT2G18860-3

**S8 Fig. The transgenic lines are visibly indistinguishable from the wild-type plants.** Columbia (Col-0) ecotype of *Arabidopsis thaliana* was used as the wild-type control for phenotype comparison, and the Col-0 ecotype was used for generating almost all the transgenic plants except for the complementation test in *nik1-1* and *at2g18860-1* mutants. The genotypes are the same as in Fig. 5. Plants were grown in a growth chamber at 22°C under long-day conditions (16 h light/8 h dark). **(A)** Developmental phenotypes associated with inactivation of NISP gene and overexpression of NISP-GFP in the R3 generation of transgenic lines at 45 days after germination. **(B)** Developmental phenotypes associated with inactivation of NISP gene and overexpression of NISP-GFP in the R3 generation of transgenic lines at 60 days after germination. **(C)** Developmental phenotypes associated with inactivation of AT2G18860 gene and overexpression of AT2G18860-GFP in the R2 generation of transgenic lines at 45 days after germination. **(D)** Developmental phenotypes associated with inactivation of AT2G18860 gene and overexpression of AT2G18860-GFP in the R2 generation of transgenic lines at 60 days after germination

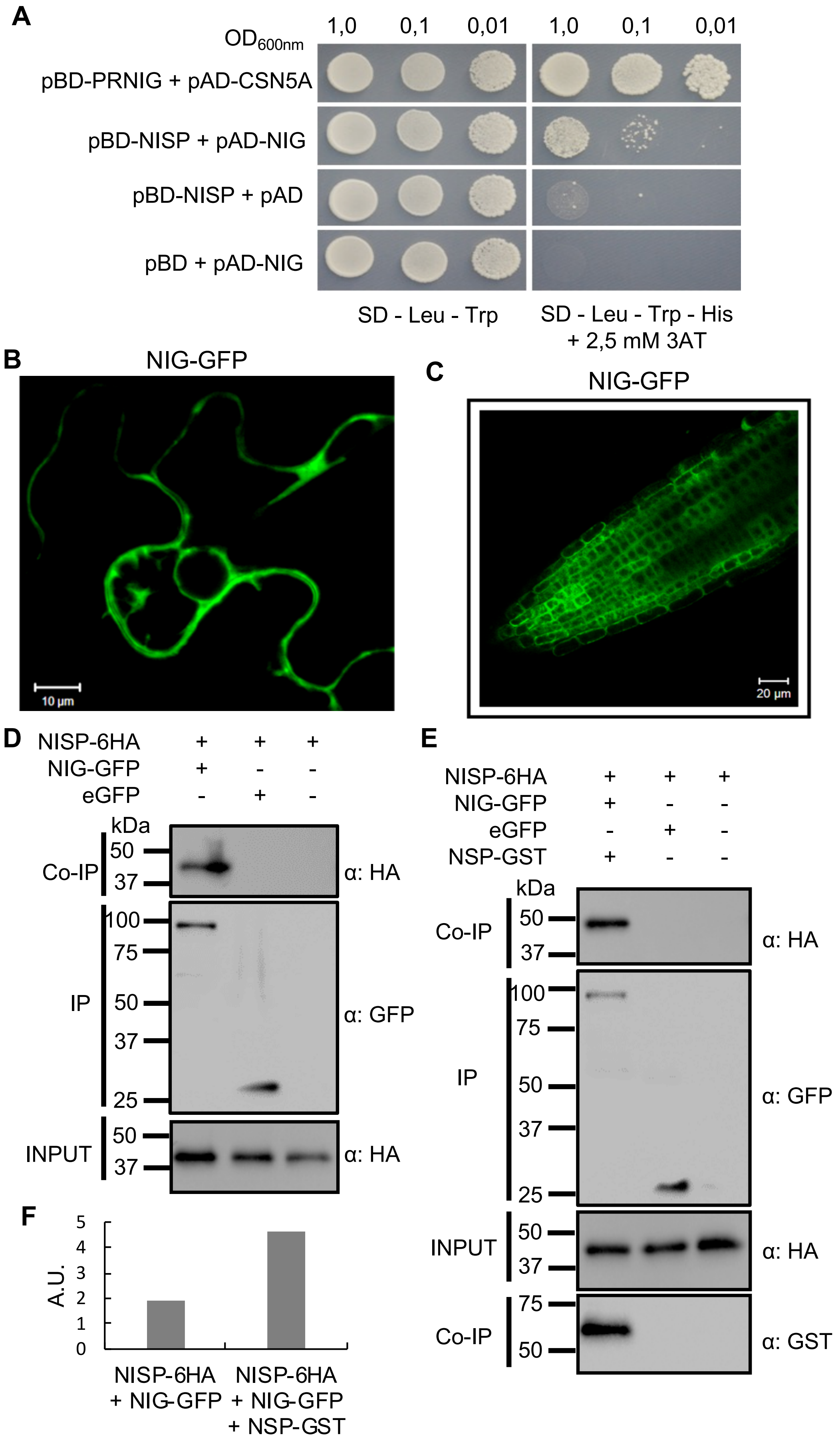

A
OD600nm
1,0
0,1
0,01
1,0
0,1
0,01
pBD-PRNIG + pAD-CSN5A
pBD-NISP + pAD-NIG
pBD-NISP + pAD
pBD + pAD-NIG
SD - Leu - Trp
SD - Leu - Trp - His
+ 2,5 mM 3AT
B
NIG-GFP
10 µm
C
NIG-GFP
20 µm
D
NISP-6HA + + +
NIG-GFP + - -
eGFP - + -
kDa
Co-IP
50
37
α: HA
IP
100
75
50
37
25
α: GFP
INPUT
50
37
α: HA
E
NISP-6HA + + +
NIG-GFP + - -
eGFP - + -
NSP-GST + - -
kDa
Co-IP
50
37
α: HA
IP
100
75
50
37
25
α: GFP
INPUT
50
37
α: HA
Co-IP
75
50
α: GST
F
A.U.
5
4
3
2
1
0
NISP-6HA
+ NIG-GFP
NISP-6HA
+ NIG-GFP
+ NSP-GST

**S9 Fig. NSP enhances NISP-NIG complex formation.**

**(A)** NISP interacts with NIG in yeast. AD-NIG and BD-NISP fusions were expressed in yeast and interactions between the recombinant proteins were monitored by His prototrophy in selective medium (-Leu,-Trp,-His) and supplemented with 2.5 mM 3-AT. The co-expression of PRNIG fused to BD and CSN5A fused to AD was used as a positive control. **(B)** Cytosolic and perinuclear distribution of NIG-GFP. *N. benthamiana* leaves were infiltrated with A. tumefaciens carrying NIG-GFP constructs with expression driven by the 35S constitutive promoter. NIG-GFP was imaged by confocal microscopy 36 h after infiltration. **(C)** Cytosolic localization of NIG in transgenic lines. Confocal fluorescence image of Arabidopsis root cells stably transformed with 35S:NIG-GFP. **(D)** NISP interacts with NIG *in planta*. Total protein extracts from *N. benthamiana* expressing NISP-6HA and NIG-GFP were used for co-immunoprecipitation assays using anti-GFP. Input and IP show the levels of the expressed proteins NISP-6HA and NIG-GFP. Anti-HA was used to detect NISP-6HA from the immunoprecipitated complex. GFP was used as an unrelated protein. The experiment was repeated three times. **(E)** NISP-NIG complex formation in the presence of viral NSP. The Co-IP assay was performed as described in A, except that co-expressed NSP-GST was included in the assay. NSP-GST was detected by immunoprecipitating it from transfected leaves and immunoblotting with anti-GST. The experiment was repeated three times. **(F)** The interaction of NISP and NIG is increased by the presence of viral NSP. NIG-GFP levels in the immunoprecipitated complex in the presence and absence of viral NSP were quantified using the Band Analysis tools of the ImageLab software (Bio-Rad). The signal values were normalized using the IP NIG-GFP band. A.U. denotes arbitrary unit.

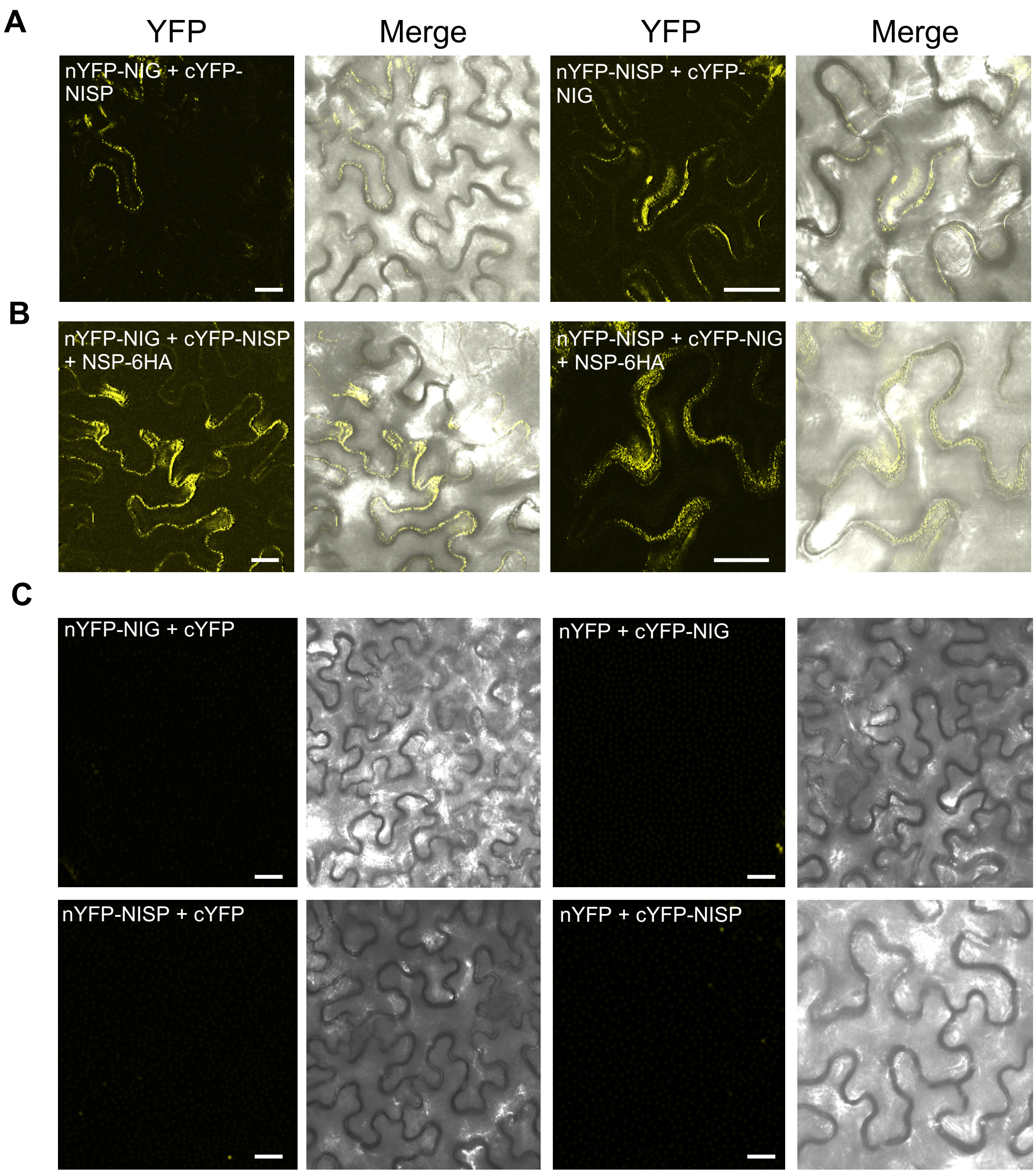


**S10 Fig. NISP interacts with NIG *in vivo* and NSP enhances NISP-NIG complex formation.**

**(A)** BIFC assay showing the interaction between NISP and NIG in vesicles of *N. benthamiana* leaf cells. *N. benthamiana* leaves transiently expressing NISP and NIG fused to the C-terminus (cYFP) or N-terminus (nYFP) of YFP, were examined by confocal microscopy 3 days after infiltration. Scale bars, 10 µm and 20 µm. **(B)** Co-expression of NSP-6HA strengthens interaction between NIG and NISP. The BiFC assay was performed by agro-infiltration of *N. benthamiana* leaves with nYFP-NIG, cYFP-NISP and vice-versus along with the expression of NSP-6HA. The reconstituted fluorescence signal was observed by confocal microscopy, 3 days after agroinfiltration. The figure displays representative samples from three independent biological repeats. Scale bars, 10 µm. **(C)** Confocal fluorescent image of NISP and NIG fused to the C-terminus (cYFP) or N-terminus (nYFP) of YFP in combination with empty vectors, as indicated in the figure.

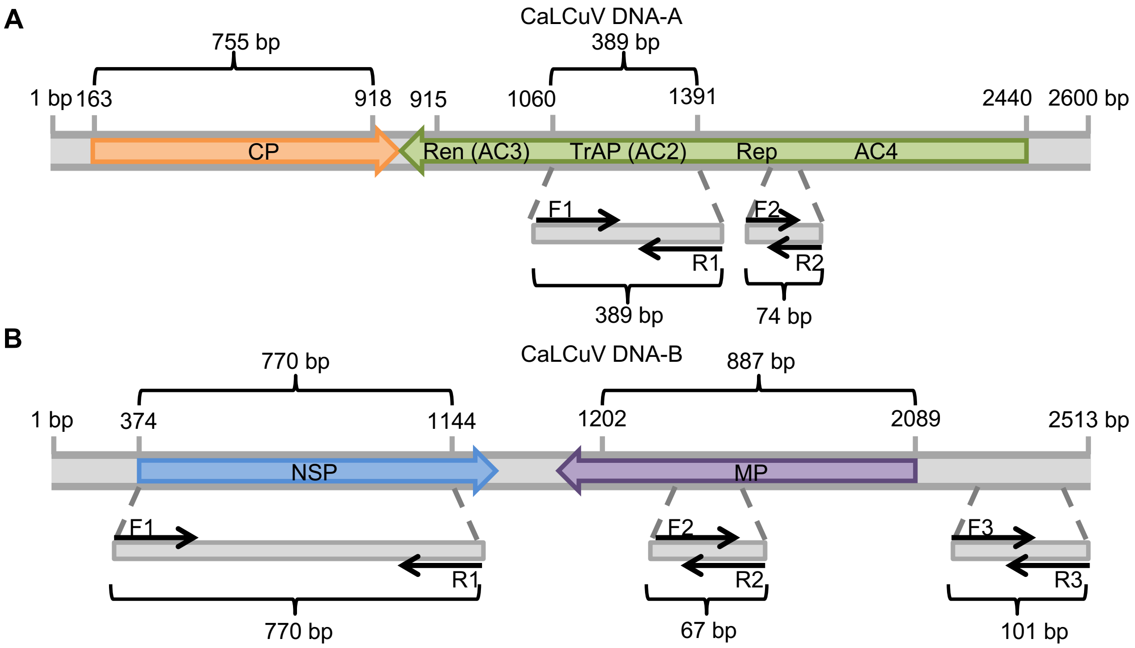


**S11 Fig. Schematic representation of CabLCV DNA-A, DNA-B and ChIP-DNA primers.** **(A)** Schematic representation of DNA-A. The number 1 corresponds to the 5' end of the nick site (TAATATT/AC) within the conserved nonanucleotide sequence located in the intergenic region. The ORFs are indicated with arrows. The positions of the primers used for PCR (F1/R1), and qRT-PCR (F2/R2) are indicated. **(B)** Schematic representation of DNA-B. The numbering scheme is the same as in a. The ORFs are indicated with arrows. The positions of the primers used for PCR (F1/R1), and qRT-PCR (F2/R2 and F3/R3) are indicated.

### S3 Table. List of primers used for cloning, PCR, RT-qPCR and qPCR

| Primer | Gene | Sequência 5'–3' |
|---|---|---|
| attB1_2942-Fwd | Gateway AttB1 site sequence | GGGGACAAGTTTGTACAAAAAAGCAGGCT |
| attB1_2943-Rvs | Gateway AttB1 site sequence | GGGGACCACTTTGTACAAGAAAGCTGGGT |
| attB4 | Gateway AttB4 site sequence | GGGGACAACTTTGTATAGAAAAGTTG |
| pDONR201/ 207_3397-Fwd | pDONR207 | TCGCGTTAACGCTAGCATGGATC |
| pDONR201/ 207_3398-Rvs | pDONR201 | TGTAACATCAGAGATTTTGAGACAC |
| 35S_MC36-Fwd | 35S promoter | TCCTTCGCAAGACCCTTCCTC |
| GFP_4799-Rvs | Green Fluorescent Protein (GFP) | CGCCCTCGCCCTCGCCGGACAC |
| DEST32-Fwd | pDEST32 | AACCGAAGTGCGCCAAGTGTCTG |
| DEST22-Fwd | pDEST22 | TATAACGCGTTTGGAATCACT |
| DEST22-Rvs | pDEST22 | AGCCGACAACCTTGATTGGAGAC |
| At4g30240-Fwd | At4g30240/NIS | AAAAAGCAGGCTTCACAATGATGGTAGCGAATAG |
| At4g30240-Ns-Rvs | At4g30240/NISP | AGAAAGCTGGGTCAGTTCTCAAGATGAATAATAG |
| At4g30240-St-Rvs | At4g30240/NISP | AGAAAGCTGGGTCTTAAGTTCTCAAGATGAATAA |
| At2g18860-Fwd | At2g18860 | AAAAAGCAGGCTTCACAATGATGGTAGTGAACAG |
| At2g18860-Ns-Rvs | At2g18860 | AGAAAGCTGGGTCAGACAAATACACTACGAAGGG |
| At2g18860-St-Rvs | At2g18860 | AGAAAGCTGGGTCTCAAGACAAATACACTACG |
| At4g30240(1-107)-Fwd | At4g30240/NIS (1-107aa) | AAAAAGCAGGCTTCACAATGATGGTAGCGAATAG |
| At4g30240(1-107)-Rvs | At4g30240/NIS (1-107aa) | AGAAAGCTGGGTCTTACCACCCAACGCAAAG |
| At4g30240(105-200)-Fwd | At4g30240/NIS (105-200aa) | AAAAAGCAGGCTTCACAATGTGGGTGGATCTTAA |
| At4g30240(105-200)-Rvs | At4g30240/NIS (105-200aa) | AGAAAGCTGGGTCTTACACTGATATTTGGTAC |
| At4g30240(200-300)-Fwd | At4g30240/NIS (200-300aa) | AAAAAGCAGGCTTCACAATGGCGTTGAGGATTAA |
| At4g30240(200-300)-Rvs | At4g30240/NIS (200-300aa) | AGAAAGCTGGGTCTTAAGTTCTCAAGATGAATAA |
| pAt4g30240-Fwd | At4g30240/NIS promoter | AGAAAAGTTGTCTGGGAGTGAGAAGATAT |
| pAt4g30240-Rvs | At4g30240/NIS promoter | GTACAAACTTGCCTTCGAAACCACACCTAAA |
| NIG_4263-Fwd | At4G13350/NIG | AAAAAGCAGGCTTCACAATGGCGGGTCGAGTTAA |
| NIG_4264-Ns-Rvs | At4G13350/NIG | AGAAAGCTGGGTCTTACCCAAATGGGTTTCCTCC |
| NIG_4265-St-Rvs | At4G13350/NIG | AGAAAGCTGGGTCCCCAAATGGGTTTCCTCCTGA |
| AtSYTA-Fwd | SYTA | GGGGACAAGTTTGTACAAAAAAGCAGGCTTCACAA TGGGCTTTTTCAGTACGATACTAG |
| AtSYTA-NS-Rvs | SYTA | GGGGACCACTTTGTACAAGAAAGCTGGGTCAGAGG CAGTTCGCCAC |
| AtFLS2-Fwd: | FLS2 | AAAAAGCAGGCTTCACAATGAAGTTACTCTCA |
| AtFLS2-NS-Rvs | FLS2 | AGAAAGCTGGGTCAACTTCTCGATCCTCGTT |
| PBL1v2040 | Begomovirus-specific primers | GCCTCTGCAGCARTGRTCKATCTTCATACA |
| PCRC1 | Begomovirus-specific primers | CTAGCTGCAGCATATTTACRARWATGCCA |
| NSP-CLCV-Fwd | Begomovirus NSP | AAAAAGCAGGCTTCACAATGTATCCT |
| NSP-CLCV-Rvs | Begomovirus NSP | AGAAAGCTGGGTCTTAACCTAAATAA |
| AC2-CaLCuV-Fwd* | Begomovirus TrAP (AC2) | AAAAAGCAGGCTTCACAATGCAAAATTCATCACT |
| AC2-CaLCuV-Rvs: | Begomovirus TrAP (AC2) | AGAAAGCTGGGTCCTACTTAAATATGTCGGCCCA |
| qRT_CaLCuV-Fwd | CaLCuV DNA-B | CCTGGGCCTGTTAGT |
| qRT_CaLCuV-Rvs | CaLCuV DNA-B | TCTTCCTCTCCCATCTTCCGT |
| qRT_CaLCuV-CompB-Fwd | CaLCuV DNA-B | TTCATGGACACCAGGAGA |
| qRT_CaLCuV-CompB-Rvs | CaLCuV DNA-B | TACACGTGTCCTATGGAGTC |
| RtCaLcuVA1 Fwd | CaLCuV DNA-A | ACAGCACGATTGAGGGTATG |
| RtCaLcuVA1 Rvs | CaLCuV DNA-A | AAAGGGACTGGCAATCAAAC |
| qRT_18SRNA-Fwd | *Arabidopsis thaliana* 18SRNA | TTTGCGCGCCTGCTGCC |
| qRT_18SRNA-Rvs | *Arabidopsis thaliana* 18SRNA | TGTGCTGGCGACGCATCATT |
| qRT_At2g18860-Fwd | At2g18860 | ATGGCAGTTGGAGGAGTTTGA |
| qRT_At2g18860-Rvs | At2g18860 | GCCGTGTGGATGAAGAATCA |
| qRT_At4g30240-Fwd | At4g30240/NIS | CAAGCTGCTTTGAGCACTGCTA |
| qRT_At4g30240-Rvs | At4g30240/NIS | TCTCGACAATTTCCATGGCTTA |
| qRT_actin-Fwd | *Arabidopsis thaliana* actin | ATGTCGTGAGCCATCCTGTC |
| qRT_actin-Rvs | *Arabidopsis thaliana* actin | ACACCGGATTCGTGCGGCAT |

*CaLCuV refers to CabLCV

**S4 Table. Expression of kanamycin resistance in the T1 generation of transgenic Arabidopsis plants.**

| Transgenic Lines Selected | Kanamycin-Resistant Seedlings | Ratio | $X^{2*}$ |
|---|---|---|---|
| *nisp-1*/NISP-1 | $126^{+}/50^{-}$ | 3:1 | 0.272 |
| *nisp-*1/NISP-2 | $130^{+}/54^{-}$ | 3:1 | 0.463 |
| *nisp-1*/NISP-3 | $122^{+}/49^{-}$ | 3:1 | 0.281 |
| AtNISP-1 | $120^{+}/48^{-}$ | 3:1 | 0.285 |
| AtNISP-2 | $129^{+}/53^{-}$ | 3:1 | 0.467 |
| AtNISP-3 | $124^{+}/53^{-}$ | 3:1 | 0.609 |
| *at2g18860-1*/AT2G18860-1 | $108^{+}/45^{-}$ | 3:1 | 0.426 |
| *at2g18860-1*/AT2G18860-2 | $112^{+}/53^{-}$ | 3:1 | 1.161 |
| *at2g18860-1*/AT2G18860-3 | $121^{+}/57^{-}$ | 3:1 | 1.261 |
| AT2G18860-1 | $118^{+}/54^{-}$ | 3:1 | 0.937 |
| AT2G18860-2 | $107^{+}/53^{-}$ | 3:1 | 1.408 |
| AT2G18860-3 | $102^{+}/51^{-}$ | 3:1 | 1.469 |

* x2 tests indicate good agreement with the segregation ratio indicated.
