## Supplementary material for "A plant-specific syntaxin-6 protein contributes to the intracytoplasmic route for begomoviruses": S2 Table- Enriched GO terms in three categories, Biological Process, Molecular Function, or Cellular Component ontology

| Molecular function | | | | |
|---|---|---|---|---|
| **GOMFID** | **Pvalue** | **Count** | **Size** | **Term** |
| GO:0005515 | 4.00887556284974e-17 | 78 | 1984 | protein binding |
| GO:0003700 | 1.88487791837318e-14 | 67 | 1670 | DNA-binding transcription factor activity |
| GO:0042802 | 0.000433420750868995 | 9 | 153 | identical protein binding |
| GO:0003755 | 0.00124285259789491 | 5 | 55 | peptidyl-prolyl cis-trans isomerase activity |
| GO:0031386 | 0.00125664493447685 | 2 | 4 | protein tag |
| GO:0015297 | 0.00257470178896631 | 7 | 126 | antiporter activity |
| GO:0097159 | 0.00304424651209625 | 61 | 2954 | organic cyclic compound binding |
| GO:0004930 | 0.00308117505324363 | 2 | 6 | G protein-coupled receptor activity |
| GO:0010011 | 0.00308117505324363 | 2 | 6 | auxin binding |
| GO:0016853 | 0.00366330286207615 | 9 | 208 | isomerase activity |
| GO:1901363 | 0.00414698612148083 | 60 | 2935 | heterocyclic compound binding |
| GO:0002020 | 0.00427202625297532 | 2 | 7 | protease binding |
| GO:0005528 | 0.00442642621807514 | 3 | 23 | FK506 binding |
| GO:0044212 | 0.00562611467127364 | 3 | 25 | transcription regulatory region DNA binding |
| GO:1901505 | 0.00701774035334296 | 4 | 52 | carbohydrate derivative transmembrane transporter activity |
| GO:0031625 | 0.0071830917010464 | 2 | 9 | ubiquitin protein ligase binding |
| GO:0060589 | 0.00889254830114986 | 2 | 10 | nucleoside-triphosphatase regulator activity |
| GO:0016168 | 0.0112547155087199 | 3 | 32 | chlorophyll binding |
| GO:0000822 | 0.0146377165162235 | 1 | 1 | inositol hexakisphosphate binding |
| GO:0004514 | 0.0146377165162235 | 1 | 1 | nicotinate-nucleotide diphosphorylase (carboxylating) activity |
| GO:0005096 | 0.0146377165162235 | 1 | 1 | GTPase activator activity |
| GO:0010298 | 0.0146377165162235 | 1 | 1 | dihydrocamalexic acid decarboxylase activity |
| GO:0010314 | 0.0146377165162235 | 1 | 1 | phosphatidylinositol-5-phosphate binding |
| GO:0031210 | 0.0146377165162235 | 1 | 1 | phosphatidylcholine binding |
| GO:0031517 | 0.0146377165162235 | 1 | 1 | red light photoreceptor activity |
| GO:0033971 | 0.0146377165162235 | 1 | 1 | hydroxyisourate hydrolase activity |
| GO:0046406 | 0.0146377165162235 | 1 | 1 | magnesium protoporphyrin IX methyltransferase activity |
| GO:0051997 | 0.0146377165162235 | 1 | 1 | 2-oxo-4-hydroxy-4-carboxy-5-ureidoimidazoline decarboxylase activity |
| GO:0071771 | 0.0146377165162235 | 1 | 1 | aldehyde decarbonylase activity |
| GO:0005375 | 0.0149745413418038 | 2 | 13 | copper ion transmembrane transporter activity |
| GO:0005315 | 0.019774436662646 | 2 | 15 | inorganic phosphate transmembrane transporter activity |
| GO:0022804 | 0.0220144954938841 | 13 | 471 | active transmembrane transporter activity |
| GO:0015932 | 0.0264452042117183 | 3 | 44 | nucleobase-containing compound transmembrane transporter activity |
| GO:1901681 | 0.0279981243399283 | 2 | 18 | sulfur compound binding |
| GO:0005215 | 0.028984280143765 | 25 | 1140 | transporter activity |
| GO:0003975 | 0.0290618740767358 | 1 | 2 | UDP-N-acetylglucosamine-dolichyl-phosphate N-acetylglucosaminephosphotransferase activity |
| GO:0004134 | 0.0290618740767358 | 1 | 2 | 4-alpha-glucanotransferase activity |
| GO:0010297 | 0.0290618740767358 | 1 | 2 | heteropolysaccharide binding |

| | | | | |
|---|---|---|---|---|
| GO:0015211 | 0.0290618740767358 | 1 | 2 | purine nucleoside transmembrane transporter activity |
| GO:0015421 | 0.0290618740767358 | 1 | 2 | oligopeptide-transporting ATPase activity |
| GO:0017151 | 0.0290618740767358 | 1 | 2 | DEAD/H-box RNA helicase binding |
| GO:0030515 | 0.0290618740767358 | 1 | 2 | snoRNA binding |
| GO:0031516 | 0.0290618740767358 | 1 | 2 | far-red light photoreceptor activity |
| GO:0043495 | 0.0290618740767358 | 1 | 2 | protein membrane anchor |
| GO:0015301 | 0.0309945198512035 | 2 | 19 | anion:anion antiporter activity |
| GO:0003713 | 0.0373444742851494 | 2 | 21 | transcription coactivator activity |
| GO:0004634 | 0.0432755781599338 | 1 | 3 | phosphopyruvate hydratase activity |
| GO:0004829 | 0.0432755781599338 | 1 | 3 | threonine-tRNA ligase activity |
| GO:0005094 | 0.0432755781599338 | 1 | 3 | Rho GDP-dissociation inhibitor activity |
| GO:0008469 | 0.0432755781599338 | 1 | 3 | histone-arginine N-methyltransferase activity |
| GO:0016277 | 0.0432755781599338 | 1 | 3 | [myelin basic protein]-arginine N-methyltransferase activity |
| GO:0019789 | 0.0432755781599338 | 1 | 3 | SUMO transferase activity |
| GO:0035241 | 0.0432755781599338 | 1 | 3 | protein-arginine omega-N monomethyltransferase activity |
| GO:0035242 | 0.0432755781599338 | 1 | 3 | protein-arginine omega-N asymmetric methyltransferase activity |
| GO:0008514 | 0.0499035167572346 | 5 | 136 | organic anion transmembrane transporter activity |

| **Cellular component** | | | | |
|---|---|---|---|---|
| **GOCCID** | **Pvalue** | **Count** | **Size** | **Term** |
| GO:0044446 | 5.87144883390372e-08 | 59 | 2203 | intracellular organelle part |
| GO:0005634 | 5.0722792776277e-07 | 170 | 9449 | nucleus |
| GO:0005829 | 5.49418404918242e-07 | 48 | 1688 | cytosol |
| GO:0005886 | 1.23160609690527e-06 | 75 | 3262 | plasma membrane |
| GO:0005623 | 2.73259892475091e-06 | 314 | 21616 | cell |
| GO:0005773 | 3.31954023771593e-06 | 27 | 756 | vacuole |
| GO:0043227 | 3.62323530219752e-05 | 228 | 15770 | membrane-bounded organelle |
| GO:0031974 | 8.39230058702016e-05 | 19 | 522 | membrane-enclosed lumen |
| GO:0009506 | 8.41601053186774e-05 | 26 | 852 | plasmodesma |
| GO:0030054 | 8.74018391894894e-05 | 26 | 854 | cell junction |
| GO:0005730 | 0.000152518357892879 | 13 | 290 | nucleolus |
| GO:0005788 | 0.000262260794736382 | 3 | 10 | endoplasmic reticulum lumen |
| GO:0005622 | 0.00035878312216202 | 287 | 19708 | intracellular |
| GO:0008180 | 0.000606763562613532 | 3 | 13 | COP9 signalosome |
| GO:0031976 | 0.000635737973665555 | 15 | 425 | plastid thylakoid |
| GO:0043229 | 0.000661848252813021 | 189 | 14265 | intracellular organelle |
| GO:0019005 | 0.000819279639568185 | 4 | 32 | SCF ubiquitin ligase complex |
| GO:0031090 | 0.000921869699116923 | 24 | 890 | organelle membrane |
| GO:0016020 | 0.00107967727187362 | 22 | 914 | membrane |
| GO:0009535 | 0.00137507961273567 | 12 | 322 | chloroplast thylakoid membrane |
| GO:0070013 | 0.00139426101826737 | 16 | 512 | intracellular organelle lumen |
| GO:0042651 | 0.00211137931743623 | 12 | 339 | thylakoid membrane |
| GO:0005774 | 0.00220173326312835 | 15 | 482 | vacuolar membrane |
| GO:0000151 | 0.00265721139541112 | 8 | 177 | ubiquitin ligase complex |

| | | | | |
|---|---|---|---|---|
| GO:0005654 | 0.00277392730711284 | 7 | 140 | nucleoplasm |
| GO:0030076 | 0.00433552987425912 | 3 | 25 | light-harvesting complex |
| GO:0044436 | 0.00474295591203549 | 11 | 330 | thylakoid part |
| GO:0005794 | 0.00686721351123819 | 22 | 928 | Golgi apparatus |
| GO:0005783 | 0.00688789123024271 | 15 | 546 | endoplasmic reticulum |
| GO:0016592 | 0.00728738490936164 | 3 | 30 | mediator complex |
| GO:0022626 | 0.00794528877603799 | 10 | 304 | cytosolic ribosome |
| GO:0009538 | 0.00898761215174485 | 2 | 11 | photosystem I reaction center |
| GO:0031967 | 0.0098942220598519 | 21 | 902 | organelle envelope |
| GO:0009941 | 0.0104354962724681 | 15 | 573 | chloroplast envelope |
| GO:0032991 | 0.0105216364859542 | 29 | 1423 | protein-containing complex |
| GO:0005681 | 0.0106908582056233 | 2 | 12 | spliceosomal complex |
| GO:0005618 | 0.0122395862462402 | 15 | 584 | cell wall |
| GO:0009536 | 0.0127340270384156 | 69 | 3997 | plastid |
| GO:0010287 | 0.0129129504493737 | 4 | 68 | plastoglobule |
| GO:0048046 | 0.0130428799484902 | 10 | 328 | apoplast |
| GO:0043224 | 0.0133210656852553 | 1 | 1 | nuclear SCF ubiquitin ligase complex |
| GO:0043228 | 0.0171462843051214 | 21 | 952 | non-membrane-bounded organelle |
| GO:0044434 | 0.0189036767217474 | 11 | 413 | chloroplast part |
| GO:0009532 | 0.0213092636554229 | 16 | 682 | plastid stroma |
| GO:0016604 | 0.0246719372265525 | 3 | 47 | nuclear body |
| GO:0000307 | 0.0264652063872122 | 1 | 2 | cyclin-dependent protein kinase holoenzyme complex |
| GO:0019773 | 0.0264652063872122 | 1 | 2 | proteasome core complex, alpha-subunit complex |
| GO:0031201 | 0.0264652063872122 | 1 | 2 | SNARE complex |
| GO:0009521 | 0.0304869302868607 | 3 | 51 | photosystem |
| GO:0005768 | 0.0323265717026583 | 8 | 276 | endosome |
| GO:0044425 | 0.0337219576019642 | 19 | 901 | membrane part |
| GO:0000502 | 0.0369574007257079 | 3 | 55 | proteasome complex |
| GO:1905368 | 0.0369574007257079 | 3 | 55 | peptidase complex |
| GO:0016363 | 0.0394347649675068 | 1 | 3 | nuclear matrix |
| GO:0031519 | 0.0394347649675068 | 1 | 3 | PcG protein complex |
| GO:1902911 | 0.0394347649675068 | 1 | 3 | protein kinase complex |
| GO:0000325 | 0.0462380007296454 | 4 | 101 | plant-type vacuole |

| **Biological process** | | | | |
|---|---|---|---|---|
| **GOBPID** | **Pvalue** | **Count** | **Size** | **Term** |
| GO:0009733 | 4.97716598806653e-30 | 50 | 424 | response to auxin |
| GO:0010583 | 8.97417535147843e-10 | 16 | 148 | response to cyclopentenone |
| GO:0042221 | 4.10032317321509e-09 | 30 | 729 | response to chemical |
| GO:0009741 | 1.75198718687499e-08 | 13 | 114 | response to brassinosteroid |
| GO:0048364 | 1.91812704424336e-08 | 15 | 164 | root development |
| GO:0046686 | 2.27612337310151e-07 | 24 | 470 | response to cadmium ion |
| GO:0009416 | 1.77769309091465e-06 | 40 | 1190 | response to light stimulus |
| GO:0009605 | 3.49450350944116e-06 | 56 | 2009 | response to external stimulus |
| GO:0010154 | 9.11375003153309e-06 | 28 | 743 | fruit development |
| GO:0006457 | 9.40940875549832e-06 | 16 | 290 | protein folding |
| GO:0019363 | 1.59518903511332e-05 | 13 | 206 | pyridine nucleotide biosynthetic process |

| | | | | |
|---|---|---|---|---|
| GO:0061458 | 3.12283217327476e-05 | 48 | 1736 | reproductive system development |
| GO:0009638 | 3.26585316721751e-05 | 4 | 13 | phototropism |
| GO:0006096 | 4.53841266783276e-05 | 12 | 196 | glycolytic process |
| GO:0006165 | 4.53841266783276e-05 | 12 | 196 | nucleoside diphosphate phosphorylation |
| GO:0010072 | 4.902039903449e-05 | 6 | 44 | primary shoot apical meristem specification |
| GO:0009734 | 4.95439687587058e-05 | 7 | 64 | auxin-activated signaling pathway |
| GO:0009135 | 5.00889787643229e-05 | 12 | 198 | purine nucleoside diphosphate metabolic process |
| GO:0009185 | 5.00889787643229e-05 | 12 | 198 | ribonucleoside diphosphate metabolic process |
| GO:0046031 | 5.00889787643229e-05 | 12 | 198 | ADP metabolic process |
| GO:1901292 | 5.00889787643229e-05 | 12 | 198 | nucleoside phosphate catabolic process |
| GO:0046496 | 5.54559379761659e-05 | 17 | 372 | nicotinamide nucleotide metabolic process |
| GO:0009793 | 6.98866400758898e-05 | 22 | 576 | embryo development ending in seed dormancy |
| GO:0072524 | 7.92796246064452e-05 | 17 | 383 | pyridine-containing compound metabolic process |
| GO:0009206 | 8.8180580616966e-05 | 12 | 210 | purine ribonucleoside triphosphate biosynthetic process |
| GO:0009142 | 0.000100901481387967 | 12 | 213 | nucleoside triphosphate biosynthetic process |
| GO:0009630 | 0.000119945357425803 | 10 | 154 | gravitropism |
| GO:0032870 | 0.000124690904113881 | 28 | 869 | cellular response to hormone stimulus |
| GO:0009168 | 0.000125614445269351 | 12 | 218 | purine ribonucleoside monophosphate biosynthetic process |
| GO:0048731 | 0.000132127031310198 | 17 | 440 | system development |
| GO:0009124 | 0.000136862105128763 | 12 | 220 | nucleoside monophosphate biosynthetic process |
| GO:0010467 | 0.000165122551831446 | 76 | 3381 | gene expression |
| GO:0009408 | 0.000187990890550127 | 14 | 298 | response to heat |
| GO:0009199 | 0.000198796639189016 | 12 | 229 | ribonucleoside triphosphate metabolic process |
| GO:0046034 | 0.000198796639189016 | 12 | 229 | ATP metabolic process |
| GO:0009144 | 0.000215411461041235 | 12 | 231 | purine nucleoside triphosphate metabolic process |
| GO:0031347 | 0.000219779473021766 | 20 | 541 | regulation of defense response |
| GO:0051188 | 0.00025791335232165 | 20 | 546 | cofactor biosynthetic process |
| GO:0006733 | 0.000280743999302022 | 17 | 426 | oxidoreduction coenzyme metabolic process |
| GO:0034654 | 0.000285220619282841 | 62 | 2656 | nucleobase-containing compound biosynthetic process |
| GO:0009126 | 0.00029423546343727 | 12 | 239 | purine nucleoside monophosphate metabolic process |
| GO:0009152 | 0.00029423546343727 | 12 | 239 | purine ribonucleotide biosynthetic process |
| GO:0009161 | 0.000329523369596703 | 12 | 242 | ribonucleoside monophosphate metabolic process |
| GO:0072522 | 0.000342874992759109 | 13 | 279 | purine-containing compound biosynthetic process |
| GO:0051179 | 0.000361232944972185 | 69 | 3104 | localization |
| GO:0009651 | 0.000364023768470993 | 25 | 782 | response to salt stress |
| GO:0046364 | 0.000482061855222544 | 10 | 183 | monosaccharide biosynthetic process |
| GO:0002376 | 0.000490892965086498 | 29 | 986 | immune system process |
| GO:0050789 | 0.000533600382232124 | 81 | 3999 | regulation of biological process |
| GO:0009644 | 0.000627867652618181 | 11 | 224 | response to high light intensity |
| GO:0010015 | 0.000638832499529958 | 17 | 458 | root morphogenesis |
| GO:0006090 | 0.000661250636464564 | 16 | 418 | pyruvate metabolic process |
| GO:0080090 | 0.0007864731152411 | 13 | 324 | regulation of primary metabolic process |
| GO:0010051 | 0.000804787184871281 | 7 | 100 | xylem and phloem pattern formation |
| GO:0048527 | 0.000867337112215913 | 6 | 75 | lateral root development |

| | | | | |
|---|---|---|---|---|
| GO:0006094 | 0.000888390846011083 | 9 | 164 | gluconeogenesis |
| GO:0009889 | 0.00089542650310783 | 44 | 1826 | regulation of biosynthetic process |
| GO:1901576 | 0.000952584494523197 | 116 | 6000 | organic substance biosynthetic process |
| GO:0009963 | 0.000958711324163015 | 7 | 103 | positive regulation of flavonoid biosynthetic process |
| GO:0009607 | 0.000998643879053542 | 37 | 1434 | response to biotic stimulus |
| GO:0009723 | 0.00101168194405963 | 14 | 353 | response to ethylene |
| GO:0006833 | 0.00104821136040308 | 8 | 136 | water transport |
| GO:0051171 | 0.00107740426452198 | 9 | 179 | regulation of nitrogen compound metabolic process |
| GO:0019318 | 0.00115311909989046 | 12 | 279 | hexose metabolic process |
| GO:0034645 | 0.00135873969906413 | 75 | 3568 | cellular macromolecule biosynthetic process |
| GO:0051049 | 0.00140053836185885 | 9 | 175 | regulation of transport |
| GO:0016070 | 0.00146217800128918 | 67 | 3115 | RNA metabolic process |
| GO:0046390 | 0.00153682202494342 | 14 | 369 | ribose phosphate biosynthetic process |
| GO:0009628 | 0.00158928234972594 | 27 | 1095 | response to abiotic stimulus |
| GO:0090696 | 0.00164914601436766 | 6 | 85 | post-embryonic plant organ development |
| GO:0010311 | 0.00165048880305198 | 4 | 34 | lateral root formation |
| GO:0071695 | 0.00172446512286255 | 17 | 502 | anatomical structure maturation |
| GO:0048438 | 0.00183576830358636 | 17 | 505 | floral whorl development |
| GO:0009615 | 0.00197681997142452 | 9 | 184 | response to virus |
| GO:0009743 | 0.00208616017830384 | 7 | 120 | response to carbohydrate |
| GO:0009955 | 0.00216342884729418 | 6 | 88 | adaxial/abaxial pattern specification |
| GO:0022414 | 0.00217785064206821 | 53 | 2369 | reproductive process |
| GO:0010148 | 0.0022023826808173 | 2 | 5 | transpiration |
| GO:0051090 | 0.0022023826808173 | 2 | 5 | regulation of DNA-binding transcription factor activity |
| GO:0016925 | 0.00232629445452831 | 3 | 18 | protein sumoylation |
| GO:0048583 | 0.00238008983221033 | 12 | 311 | regulation of response to stimulus |
| GO:0007165 | 0.00249139842178076 | 31 | 1222 | signal transduction |
| GO:0009269 | 0.00250583052534263 | 4 | 38 | response to desiccation |
| GO:0009057 | 0.00250987792136539 | 20 | 658 | macromolecule catabolic process |
| GO:0007155 | 0.0027078243939711 | 6 | 92 | cell adhesion |
| GO:0042542 | 0.00282432271867728 | 9 | 194 | response to hydrogen peroxide |
| GO:0009791 | 0.00303867113360698 | 27 | 1090 | post-embryonic development |
| GO:0090558 | 0.00313141819446908 | 20 | 671 | plant epidermis development |
| GO:0034976 | 0.00318213121082329 | 13 | 357 | response to endoplasmic reticulum stress |
| GO:0019439 | 0.00320648412930211 | 16 | 488 | aromatic compound catabolic process |
| GO:0044270 | 0.00320648412930211 | 16 | 488 | cellular nitrogen compound catabolic process |
| GO:0042176 | 0.00327067166251161 | 2 | 6 | regulation of protein catabolic process |
| GO:0046700 | 0.00333615237515155 | 16 | 490 | heterocycle catabolic process |
| GO:0010035 | 0.00335642611074 | 21 | 755 | response to inorganic substance |
| GO:0009409 | 0.00347115220345317 | 19 | 630 | response to cold |
| GO:0060255 | 0.00347532058474022 | 40 | 1741 | regulation of macromolecule metabolic process |
| GO:0048439 | 0.00363518135235425 | 5 | 68 | flower morphogenesis |
| GO:0000413 | 0.00369976000448799 | 3 | 21 | protein peptidyl-prolyl isomerization |
| GO:0006855 | 0.00369976000448799 | 3 | 21 | drug transmembrane transport |
| GO:1901361 | 0.00375139037427478 | 16 | 496 | organic cyclic compound catabolic process |
| GO:0032501 | 0.00387118284571784 | 34 | 1576 | multicellular organismal process |

| | | | | |
|---|---|---|---|---|
| GO:0010218 | 0.00389927925584597 | 6 | 99 | response to far red light |
| GO:0006163 | 0.00392436410075207 | 13 | 366 | purine nucleotide metabolic process |
| GO:0090066 | 0.00393338191966936 | 8 | 167 | regulation of anatomical structure size |
| GO:0048513 | 0.00407739328387982 | 8 | 168 | animal organ development |
| GO:0009886 | 0.00423654232184118 | 3 | 22 | post-embryonic animal morphogenesis |
| GO:0010158 | 0.00453339231014114 | 2 | 7 | abaxial cell fate specification |
| GO:0010014 | 0.0045339238858717 | 8 | 171 | meristem initiation |
| GO:0010114 | 0.00466298340092407 | 6 | 103 | response to red light |
| GO:1901701 | 0.0046793879416626 | 21 | 752 | cellular response to oxygen-containing compound |
| GO:0045010 | 0.00473382062481921 | 6 | 103 | actin nucleation |
| GO:0032273 | 0.00496151170162504 | 6 | 104 | positive regulation of protein polymerization |
| GO:0071229 | 0.00499251682776117 | 20 | 700 | cellular response to acid chemical |
| GO:0009880 | 0.00504186968708695 | 4 | 46 | embryonic pattern specification |
| GO:0009965 | 0.00536543821860449 | 9 | 214 | leaf morphogenesis |
| GO:0090567 | 0.0054318007518676 | 25 | 954 | reproductive shoot system development |
| GO:0044089 | 0.00544087761810676 | 6 | 106 | positive regulation of cellular component biogenesis |
| GO:0051495 | 0.00544087761810676 | 6 | 106 | positive regulation of cytoskeleton organization |
| GO:1902905 | 0.00544087761810676 | 6 | 106 | positive regulation of supramolecular fiber organization |
| GO:0048523 | 0.0057341856202804 | 21 | 758 | negative regulation of cellular process |
| GO:0009751 | 0.00575390574192411 | 15 | 473 | response to salicylic acid |
| GO:0009753 | 0.00586316469844461 | 15 | 474 | response to jasmonic acid |
| GO:0019637 | 0.00590975809530124 | 29 | 1167 | organophosphate metabolic process |
| GO:0006091 | 0.00594823701256159 | 18 | 615 | generation of precursor metabolites and energy |
| GO:0010155 | 0.00617581727890846 | 5 | 77 | regulation of proton transport |
| GO:0030833 | 0.00622227737678641 | 6 | 109 | regulation of actin filament polymerization |
| GO:0005975 | 0.00637036716304114 | 18 | 634 | carbohydrate metabolic process |
| GO:0030832 | 0.00650000715477068 | 6 | 110 | regulation of actin filament length |
| GO:0032970 | 0.00650000715477068 | 6 | 110 | regulation of actin filament-based process |
| GO:0110053 | 0.00650000715477068 | 6 | 110 | regulation of actin filament organization |
| GO:0048532 | 0.00704241250597826 | 10 | 264 | anatomical structure arrangement |
| GO:0006979 | 0.00730978815001429 | 15 | 486 | response to oxidative stress |
| GO:0043254 | 0.00738720220068824 | 6 | 113 | regulation of protein complex assembly |
| GO:0015802 | 0.0076123227049012 | 3 | 27 | basic amino acid transport |
| GO:0048367 | 0.00793878439726216 | 14 | 461 | shoot system development |
| GO:0097659 | 0.00804568369693227 | 47 | 2184 | nucleic acid-templated transcription |
| GO:0010090 | 0.00840189181564488 | 7 | 152 | trichome morphogenesis |
| GO:0032268 | 0.00880953488316344 | 10 | 273 | regulation of cellular protein metabolic process |
| GO:0008154 | 0.00905528072609012 | 6 | 118 | actin polymerization or depolymerization |
| GO:0034765 | 0.00905528072609012 | 6 | 118 | regulation of ion transmembrane transport |
| GO:0016310 | 0.00933656092354678 | 24 | 946 | phosphorylation |
| GO:0006810 | 0.00969844840128636 | 34 | 1560 | transport |
| GO:0045087 | 0.0102480355213813 | 11 | 329 | innate immune response |
| GO:0048481 | 0.0102704271345608 | 7 | 158 | plant ovule development |
| GO:0048765 | 0.0103324655935503 | 11 | 323 | root hair cell differentiation |
| GO:0048469 | 0.0105566567998059 | 11 | 324 | cell maturation |
| GO:2001141 | 0.010810791625008 | 35 | 1580 | regulation of RNA biosynthetic process |

| | | | | |
|---|---|---|---|---|
| GO:0070887 | 0.0108848077284432 | 11 | 340 | cellular response to chemical stimulus |
| GO:0090697 | 0.010898655523811 | 10 | 282 | post-embryonic plant organ morphogenesis |
| GO:0015696 | 0.0111920361898402 | 3 | 31 | ammonium transport |
| GO:0010325 | 0.0114091778565285 | 2 | 11 | raffinose family oligosaccharide biosynthetic process |
| GO:1901700 | 0.0114793876677908 | 18 | 746 | response to oxygen-containing compound |
| GO:0006612 | 0.0117711052061691 | 12 | 374 | protein targeting to membrane |
| GO:0009259 | 0.0118868945454564 | 14 | 467 | ribonucleotide metabolic process |
| GO:0090627 | 0.0120015213181777 | 12 | 375 | plant epidermal cell differentiation |
| GO:0050776 | 0.0120319639993915 | 13 | 421 | regulation of immune response |
| GO:0009812 | 0.0121199104521229 | 9 | 244 | flavonoid metabolic process |
| GO:0030243 | 0.012259209187742 | 6 | 126 | cellulose metabolic process |
| GO:0072657 | 0.0124727502329178 | 12 | 377 | protein localization to membrane |
| GO:0009737 | 0.012557616237491 | 17 | 615 | response to abscisic acid |
| GO:0009725 | 0.0125997123013608 | 3 | 40 | response to hormone |
| GO:0019344 | 0.0128824792301812 | 8 | 205 | cysteine biosynthetic process |
| GO:0010152 | 0.0135556754075667 | 2 | 12 | pollen maturation |
| GO:0006073 | 0.0137129899126579 | 12 | 382 | cellular glucan metabolic process |
| GO:0030036 | 0.0144547074550275 | 7 | 169 | actin cytoskeleton organization |
| GO:0009744 | 0.0147077433743487 | 8 | 210 | response to sucrose |
| GO:0044267 | 0.015001281798713 | 49 | 2428 | cellular protein metabolic process |
| GO:0009612 | 0.0150867624933634 | 4 | 63 | response to mechanical stimulus |
| GO:0001778 | 0.0150890975282622 | 1 | 1 | plasma membrane repair |
| GO:0006574 | 0.0150890975282622 | 1 | 1 | valine catabolic process |
| GO:0009594 | 0.0150890975282622 | 1 | 1 | detection of nutrient |
| GO:0010081 | 0.0150890975282622 | 1 | 1 | regulation of inflorescence meristem growth |
| GO:0010184 | 0.0150890975282622 | 1 | 1 | cytokinin transport |
| GO:0010202 | 0.0150890975282622 | 1 | 1 | response to low fluence red light stimulus |
| GO:0019428 | 0.0150890975282622 | 1 | 1 | allantoin biosynthetic process |
| GO:0033234 | 0.0150890975282622 | 1 | 1 | negative regulation of protein sumoylation |
| GO:0043547 | 0.0150890975282622 | 1 | 1 | positive regulation of GTPase activity |
| GO:0043619 | 0.0150890975282622 | 1 | 1 | regulation of transcription from RNA polymerase II promoter in response to oxidative stress |
| GO:0045014 | 0.0150890975282622 | 1 | 1 | carbon catabolite repression of transcription by glucose |
| GO:0045990 | 0.0150890975282622 | 1 | 1 | carbon catabolite regulation of transcription |
| GO:0046015 | 0.0150890975282622 | 1 | 1 | regulation of transcription by glucose |
| GO:0046827 | 0.0150890975282622 | 1 | 1 | positive regulation of protein export from nucleus |
| GO:0061985 | 0.0150890975282622 | 1 | 1 | carbon catabolite repression |
| GO:0070207 | 0.0150890975282622 | 1 | 1 | protein homotrimerization |
| GO:0072660 | 0.0150890975282622 | 1 | 1 | maintenance of protein location in plasma membrane |
| GO:1900088 | 0.0150890975282622 | 1 | 1 | regulation of inositol biosynthetic process |
| GO:1900091 | 0.0150890975282622 | 1 | 1 | regulation of raffinose biosynthetic process |
| GO:1990778 | 0.0150890975282622 | 1 | 1 | protein localization to cell periphery |
| GO:0048229 | 0.0161690087416735 | 15 | 534 | gametophyte development |
| GO:0045892 | 0.0161801748472776 | 12 | 391 | negative regulation of transcription, DNA-templated |

| | | | | |
|---|---|---|---|---|
| GO:1902679 | 0.0161801748472776 | 12 | 391 | negative regulation of RNA biosynthetic process |
| GO:0019219 | 0.0167545109733366 | 36 | 1686 | regulation of nucleobase-containing compound metabolic process |
| GO:0009646 | 0.018109101120796 | 3 | 37 | response to absence of light |
| GO:0050832 | 0.0181425162295137 | 11 | 351 | defense response to fungus |
| GO:0007030 | 0.0181786361161756 | 7 | 177 | Golgi organization |
| GO:0048443 | 0.0181786361161756 | 7 | 177 | stamen development |
| GO:0036211 | 0.0182169108484553 | 38 | 1776 | protein modification process |
| GO:0071407 | 0.0182371816326785 | 11 | 354 | cellular response to organic cyclic compound |
| GO:0046483 | 0.0183132031328147 | 72 | 3925 | heterocycle metabolic process |
| GO:0015780 | 0.0183234062384548 | 2 | 14 | nucleotide-sugar transmembrane transport |
| GO:0009056 | 0.0187400987051903 | 37 | 1723 | catabolic process |
| GO:0009863 | 0.0188332448183096 | 11 | 353 | salicylic acid mediated signaling pathway |
| GO:0051707 | 0.0188350585316931 | 28 | 1249 | response to other organism |
| GO:0055086 | 0.0189453001447958 | 21 | 850 | nucleobase-containing small molecule metabolic process |
| GO:0044419 | 0.0190413017121883 | 6 | 139 | interspecies interaction between organisms |
| GO:0010054 | 0.0191858463661779 | 11 | 354 | trichoblast differentiation |
| GO:2000026 | 0.0193286438720727 | 15 | 546 | regulation of multicellular organismal development |
| GO:1901564 | 0.0196232474496549 | 73 | 4002 | organonitrogen compound metabolic process |
| GO:0009653 | 0.0203030253204579 | 20 | 839 | anatomical structure morphogenesis |
| GO:0009641 | 0.0208135982019617 | 2 | 15 | shade avoidance |
| GO:0006862 | 0.020837111546033 | 3 | 39 | nucleotide transport |
| GO:0009742 | 0.020837111546033 | 3 | 39 | brassinosteroid mediated signaling pathway |
| GO:0071383 | 0.020837111546033 | 3 | 39 | cellular response to steroid hormone stimulus |
| GO:0043933 | 0.0212232196899022 | 17 | 653 | protein-containing complex subunit organization |
| GO:0010050 | 0.0214096586507564 | 4 | 70 | vegetative phase change |
| GO:0046173 | 0.0214096586507564 | 4 | 70 | polyol biosynthetic process |
| GO:0018193 | 0.0221039808646057 | 12 | 409 | peptidyl-amino acid modification |
| GO:0045934 | 0.0221039808646057 | 12 | 409 | negative regulation of nucleobase-containing compound metabolic process |
| GO:0009750 | 0.0222210819758679 | 6 | 144 | response to fructose |
| GO:0048856 | 0.0222555686983299 | 18 | 824 | anatomical structure development |
| GO:0071236 | 0.0229861213744943 | 11 | 364 | cellular response to antibiotic |
| GO:0009266 | 0.0230267224751948 | 4 | 76 | response to temperature stimulus |
| GO:0044271 | 0.0233513378113603 | 17 | 719 | cellular nitrogen compound biosynthetic process |
| GO:0010187 | 0.0236894885045011 | 2 | 16 | negative regulation of seed germination |
| GO:2000652 | 0.0236894885045011 | 2 | 16 | regulation of secondary cell wall biogenesis |
| GO:0048767 | 0.0243340119482866 | 7 | 188 | root hair elongation |
| GO:0006725 | 0.024396082573756 | 79 | 4388 | cellular aromatic compound metabolic process |
| GO:1900140 | 0.0245473213382688 | 4 | 73 | regulation of seedling development |
| GO:0006355 | 0.0246537541619134 | 33 | 1573 | regulation of transcription, DNA-templated |
| GO:0005976 | 0.0256872507965066 | 17 | 668 | polysaccharide metabolic process |
| GO:0098662 | 0.0257409021206122 | 6 | 149 | inorganic cation transmembrane transport |
| GO:0010363 | 0.025956064870567 | 11 | 371 | regulation of plant-type hypersensitive response |
| GO:0015979 | 0.0260198756281003 | 12 | 419 | photosynthesis |
| GO:0010102 | 0.0260402853296895 | 2 | 17 | lateral root morphogenesis |
| GO:0009890 | 0.0264376550675167 | 12 | 420 | negative regulation of biosynthetic process |

| | | | | |
|---|---|---|---|---|
| GO:0005983 | 0.0265843743984608 | 2 | 17 | starch catabolic process |
| GO:0009825 | 0.02659237225398 | 5 | 111 | multidimensional cell growth |
| GO:0070727 | 0.0274833617319266 | 23 | 991 | cellular macromolecule localization |
| GO:0009415 | 0.0275864681816309 | 12 | 424 | response to water |
| GO:0030003 | 0.0279592043873693 | 9 | 282 | cellular cation homeostasis |
| GO:0006811 | 0.0286287007714681 | 25 | 1104 | ion transport |
| GO:0010182 | 0.0293864910427875 | 5 | 114 | sugar mediated signaling pathway |
| GO:0010450 | 0.0298582828594031 | 1 | 2 | inflorescence meristem growth |
| GO:0000025 | 0.0299512261164581 | 1 | 2 | maltose catabolic process |
| GO:0001560 | 0.0299512261164581 | 1 | 2 | regulation of cell growth by extracellular stimulus |
| GO:0009413 | 0.0299512261164581 | 1 | 2 | response to flooding |
| GO:0009729 | 0.0299512261164581 | 1 | 2 | detection of brassinosteroid stimulus |
| GO:0010080 | 0.0299512261164581 | 1 | 2 | regulation of floral meristem growth |
| GO:0010390 | 0.0299512261164581 | 1 | 2 | histone monoubiquitination |
| GO:0032388 | 0.0299512261164581 | 1 | 2 | positive regulation of intracellular transport |
| GO:0046822 | 0.0299512261164581 | 1 | 2 | regulation of nucleocytoplasmic transport |
| GO:0051222 | 0.0299512261164581 | 1 | 2 | positive regulation of protein transport |
| GO:0051245 | 0.0299512261164581 | 1 | 2 | negative regulation of cellular defense response |
| GO:0080171 | 0.0299512261164581 | 1 | 2 | lytic vacuole organization |
| GO:0080178 | 0.0299512261164581 | 1 | 2 | 5-carbamoylmethyl uridine residue modification |
| GO:1900111 | 0.0299512261164581 | 1 | 2 | positive regulation of histone H3-K9 dimethylation |
| GO:1902448 | 0.0299512261164581 | 1 | 2 | positive regulation of shade avoidance |
| GO:1903320 | 0.0299512261164581 | 1 | 2 | regulation of protein modification by small protein conjugation or removal |
| GO:1903829 | 0.0299512261164581 | 1 | 2 | positive regulation of cellular protein localization |
| GO:2000068 | 0.0299512261164581 | 1 | 2 | regulation of defense response to insect |
| GO:2000082 | 0.0299512261164581 | 1 | 2 | regulation of L-ascorbic acid biosynthetic process |
| GO:0009867 | 0.0302133236227664 | 9 | 286 | jasmonic acid mediated signaling pathway |
| GO:0051235 | 0.0303574569035653 | 5 | 115 | maintenance of location |
| GO:0009611 | 0.0315530178857144 | 10 | 335 | response to wounding |
| GO:0009944 | 0.031617321429223 | 4 | 79 | polarity specification of adaxial/abaxial axis |
| GO:0006950 | 0.0319512179909594 | 36 | 1962 | response to stress |
| GO:0044237 | 0.0321815067258644 | 57 | 3375 | cellular metabolic process |
| GO:1901360 | 0.0326468957007177 | 77 | 4355 | organic cyclic compound metabolic process |
| GO:0000096 | 0.0326643898148772 | 10 | 337 | sulfur amino acid metabolic process |
| GO:0006825 | 0.0327737757797424 | 2 | 19 | copper ion transport |
| GO:0051603 | 0.0328122609752537 | 12 | 434 | proteolysis involved in cellular protein catabolic process |
| GO:0010162 | 0.0338506598838323 | 6 | 159 | seed dormancy process |
| GO:0040007 | 0.0338779607180417 | 22 | 958 | growth |
| GO:0009814 | 0.0346541824321327 | 14 | 538 | defense response, incompatible interaction |
| GO:0051649 | 0.0347079836499763 | 25 | 1125 | establishment of localization in cell |
| GO:0035690 | 0.0354062546267919 | 11 | 390 | cellular response to drug |
| GO:0032989 | 0.035560130099704 | 18 | 748 | cellular component morphogenesis |
| GO:0071555 | 0.035577024216424 | 14 | 540 | cell wall organization |
| GO:0031400 | 0.0357405551326426 | 3 | 48 | negative regulation of protein modification process |

| | | | | |
|---|---|---|---|---|
| GO:0070647 | 0.0358008444693641 | 9 | 296 | protein modification by small protein conjugation or removal |
| GO:0046394 | 0.0367539903151117 | 26 | 1187 | carboxylic acid biosynthetic process |
| GO:0006511 | 0.0367705841827754 | 10 | 344 | ubiquitin-dependent protein catabolic process |
| GO:0010243 | 0.0374514648824485 | 12 | 443 | response to organonitrogen compound |
| GO:0042335 | 0.0376593011484245 | 3 | 49 | cuticle development |
| GO:0071322 | 0.0377183609768817 | 5 | 122 | cellular response to carbohydrate stimulus |
| GO:0016043 | 0.037757308278471 | 32 | 1578 | cellular component organization |
| GO:0055085 | 0.0379548577957795 | 10 | 352 | transmembrane transport |
| GO:0055082 | 0.0390980726535066 | 9 | 300 | cellular chemical homeostasis |
| GO:0009773 | 0.0396297289085993 | 3 | 50 | photosynthetic electron transport in photosystem I |
| GO:0043632 | 0.0399144925505794 | 10 | 349 | modification-dependent macromolecule catabolic process |
| GO:0009069 | 0.0403117346008656 | 8 | 255 | serine family amino acid metabolic process |
| GO:0003002 | 0.0406017257453014 | 4 | 88 | regionalization |
| GO:0015833 | 0.0408656852115269 | 24 | 1088 | peptide transport |
| GO:0045184 | 0.0413842131183136 | 22 | 979 | establishment of protein localization |
| GO:0048507 | 0.0416124165498851 | 13 | 501 | meristem development |
| GO:0034050 | 0.041870159861401 | 11 | 401 | host programmed cell death induced by symbiont |
| GO:0048440 | 0.0418827649175524 | 8 | 257 | carpel development |
| GO:0042742 | 0.0424956936916541 | 11 | 402 | defense response to bacterium |
| GO:0010197 | 0.0429881097319888 | 2 | 22 | polar nucleus fusion |
| GO:0051259 | 0.0429881097319888 | 2 | 22 | protein complex oligomerization |
| GO:0010941 | 0.0437660843224772 | 11 | 404 | regulation of cell death |
| GO:0006886 | 0.0437733208803176 | 21 | 933 | intracellular protein transport |
| GO:0010605 | 0.0438097037960778 | 17 | 714 | negative regulation of macromolecule metabolic process |
| GO:0043903 | 0.0444566666146036 | 1 | 3 | regulation of symbiosis, encompassing mutualism through parasitism |
| GO:0006435 | 0.0445897891026286 | 1 | 3 | threonyl-tRNA aminoacylation |
| GO:0009660 | 0.0445897891026286 | 1 | 3 | amyloplast organization |
| GO:0010617 | 0.0445897891026286 | 1 | 3 | circadian regulation of calcium ion oscillation |
| GO:0018027 | 0.0445897891026286 | 1 | 3 | peptidyl-lysine dimethylation |
| GO:0019388 | 0.0445897891026286 | 1 | 3 | galactose catabolic process |
| GO:0019919 | 0.0445897891026286 | 1 | 3 | peptidyl-arginine methylation, to asymmetrical-dimethyl arginine |
| GO:0032456 | 0.0445897891026286 | 1 | 3 | endocytic recycling |
| GO:0033157 | 0.0445897891026286 | 1 | 3 | regulation of intracellular protein transport |
| GO:0033530 | 0.0445897891026286 | 1 | 3 | raffinose metabolic process |
| GO:0035246 | 0.0445897891026286 | 1 | 3 | peptidyl-arginine N-methylation |
| GO:0035672 | 0.0445897891026286 | 1 | 3 | oligopeptide transmembrane transport |
| GO:0040014 | 0.0445897891026286 | 1 | 3 | regulation of multicellular organism growth |
| GO:0043447 | 0.0445897891026286 | 1 | 3 | alkane biosynthetic process |
| GO:0043620 | 0.0445897891026286 | 1 | 3 | regulation of DNA-templated transcription in response to stress |
| GO:0046740 | 0.0445897891026286 | 1 | 3 | transport of virus in host, cell to cell |
| GO:0048833 | 0.0445897891026286 | 1 | 3 | specification of floral organ number |
| GO:0050792 | 0.0445897891026286 | 1 | 3 | regulation of viral process |
| GO:0051289 | 0.0445897891026286 | 1 | 3 | protein homotetramerization |

| | | | | |
|---|---|---|---|---|
| GO:0051453 | 0.0445897891026286 | 1 | 3 | regulation of intracellular pH |
| GO:0072334 | 0.0445897891026286 | 1 | 3 | UDP-galactose transmembrane transport |
| GO:1901684 | 0.0445897891026286 | 1 | 3 | arsenate ion transmembrane transport |
| GO:2000072 | 0.0445897891026286 | 1 | 3 | regulation of defense response to fungus, incompatible interaction |
| GO:0006970 | 0.0450429772922111 | 3 | 55 | response to osmotic stress |
| GO:0022607 | 0.0452634574972824 | 20 | 879 | cellular component assembly |
| GO:0048511 | 0.0455611287425748 | 6 | 171 | rhythmic process |
| GO:0006888 | 0.0458009359811377 | 4 | 89 | ER to Golgi vesicle-mediated transport |
| GO:0007034 | 0.0460843760639176 | 5 | 129 | vacuolar transport |
| GO:0001510 | 0.0466375347311756 | 6 | 172 | RNA methylation |
| GO:0034284 | 0.0488374278922391 | 6 | 174 | response to monosaccharide |
| GO:0034622 | 0.0489563092242564 | 15 | 618 | cellular protein-containing complex assembly |
| GO:0048193 | 0.0490987110961924 | 7 | 221 | Golgi vesicle transport |
| GO:0009735 | 0.0494533793763484 | 8 | 266 | response to cytokinin |
